# Increased seed carbohydrate reserves associated with domestication influence the optimal seminal root number of *Zea mays*

**DOI:** 10.1101/2020.12.09.417691

**Authors:** Alden C. Perkins, Jonathan P. Lynch

**Affiliations:** Department of Plant Science, The Pennsylvania State University, University Park, PA 16802, USA

**Keywords:** domestication, roots, seed reserves, abiotic stress, nitrogen, phosphorus, *Zea mays* ssp. *mays*, *Zea mays* ssp. *parviglumis*

## Abstract

**Background and Aims:** Domesticated maize (*Zea mays* ssp. *mays*) generally forms between two and six seminal roots, while its wild ancestor, Mexican annual teosinte (*Zea mays* ssp. *parviglumis*), typically lacks seminal roots. Maize also produces larger seeds than teosinte, and it generally has higher growth rates as a seedling. Maize was originally domesticated in the tropical soils of southern Mexico, but it was later brought to the Mexican highlands before spreading to other parts of the continent, where it experienced different soil resource constraints. The aims of this study were to understand the impact of increased seminal root number on seedling nitrogen acquisition and to model how differences in maize and teosinte phenotypes might have contributed to increased seminal root number in domesticated maize.

**Methods:** Seedling root architectural models of a teosinte accession and a maize landrace were constructed by parameterizing the functional-structural plant model *OpenSimRoot* using plants grown in mesocosms. Seedling growth was simulated in a low-phosphorus environment, multiple low-nitrogen environments, and at variable planting densities. Models were also constructed to combine individual components of the maize and teosinte phenotypes.

**Key Results:** Seminal roots contributed about 35% of the nitrogen and phosphorus acquired by maize landrace seedlings in the first 25 days after planting. Increased seminal root number improved plant N acquisition under low-N environments with varying precipitation patterns, fertilization rates, soil textures, and planting densities. Models suggested that the optimal number of seminal roots for nutrient acquisition in teosinte is constrained by its limited seed carbohydrate reserves.

**Conclusions:** Seminal roots can improve the acquisition of both nitrogen and phosphorus in maize seedlings, and the increase in seed size associated with maize domestication may have facilitated increased seminal root number.

## Introduction

The collection of phenotypic traits that differentiate cultivated plants from their wild relatives is referred to as domestication syndrome (Hammer, 1984). Common domestication traits in plants include the loss of seed dispersal mechanisms, increased size of seeds or fruits, decreased photoperiod sensitivity, and changes in secondary metabolite production (Doebley *et al*., 2006; Meyer *et al*., 2012). While the transition from natural to agricultural ecosystems is associated with many changes in the rhizosphere, such as supplemental irrigation, the application of chemical fertilizers, control of edaphic pests and diseases, and decreased interspecific competition (Schmidt *et al*., 2016), little is known about how domestication influences root phenotypes.

Maize (*Zea mays* ssp. *mays*) was domesticated approximately 9,000 years ago from Mexican annual teosinte (*Z. mays* ssp. *parviglumis*), and the crop has several major morphological differences from its progenitor (Matsuoka *et al*., 2002). Teosintes may form many tillers and lateral branches that terminate in male inflorescences, while maize does not typically branch or tiller. The female inflorescences of teosinte are small and have two rows of seeds, while domesticated maize forms large ears with many ranks of seeds (Doebley *et al*., 1990). Teosinte kernels are small and covered by hard cupulate fruitcases that may constitute half the dry weight of the seed, while maize seeds are exposed and may be ten times larger (Doebley, 2004). Maize landrace seeds contain about 58% less protein and 34% more carbohydrates than teosinte kernels as a percentage of kernel mass (Flint-Garcia *et al*., 2009). The embryo contains about 26% of the seed protein, which is in the form of globulins, and the remaining protein is found in the endosperm (Watson, 2003; Flint-Garcia, 2017). Most of the endosperm protein is in the form of ethanol-soluble proteins called zeins, which constitute a greater percentage of the total seed protein in teosinte than maize (Paulis and Wall, 1977).

The maize root system is composed of the primary root, a variable number of seminal roots, and shoot-borne nodal roots that form in whorls initiated both below ground (crown roots) and above ground (brace roots) (Kiesselbach, 1949) as well as lateral roots on all axial root classes. Seminal root primordia are initiated around the scutellar node in the seed starting about 25 days after pollination (Erdelska and Vidovencova, 1993). They are sometimes classified as dorsal, ventral, and intermediary based on their orientation to the embryo axis, and their development begins in approximately that order. Seminal roots emerge from the seed following the primary root, often before the coleoptile has emerged from the soil, and the embryonic roots comprise the majority of the root system for the first 2-3 weeks as the adventitious roots are developing (Feldman, 1994; Hochholdinger *et al*., 2004). Maize seminal roots are of smaller diameter than the primary root and crown roots, and they have a smaller stele, fewer cortical cell files, and fewer metaxylem vessels (Burton *et al*., 2013 *b*; Tai *et al*., 2015).

Variation in seminal root number is found within maize and other monocot species, and domestication appears to have increased the average seminal root number in some species. Burton *et al*. (2013 *a*) characterized root anatomical and architectural diversity in a panel of 195 maize landraces and 61 teosinte accessions from diverse geographic origins. They observed that maize landraces had 3.9 seminal roots on average, while teosinte accessions had 0.5 seminal roots on average. The opposite pattern was observed for crown roots; landraces had 20.6 crown roots on average, while teosinte accessions had 24.0 at 28 days after planting. Increased seminal root number following domestication has also been reported in other Poales such as barley (*Hordeum vulgare*) (Grando and Ceccarelli, 1995). In wheat (*Triticum* spp.), five seminal root primordia are found in the seeds of both cultivars and wild relatives (Robertson *et al*., 1979), and all five primordia typically develop in cultivated wheat, while only three develop in wild accessions under normal conditions (Golan *et al*., 2018). Other grains, such as rice (*Oryza sativa*) and sorghum (*Sorghum bicolor*), form a primary root but no seminal roots (Hochholdinger *et al*., 2004; Singh *et al*., 2010). Sorghum is a close relative of maize; the two lineages diverged about 12 million years ago (Salvi, 2017), and sorghum was domesticated in Ethiopia and Sudan (Wendorf *et al*., 1992). Molecular evidence suggests that many of the genes involved in the formation of maize seminal root primordia may be non-syntenic to sorghum (Tai *et al*., 2017).

Teosinte and early maize landraces grew in soils with limited nutrient availability. The domestication of maize took place in the lowlands of southern Mexico, a region with tropical deciduous forests, wet summers, and dry winters (Hastorf, 2009). Highly weathered soils with low phosphorus availability are found in this region (Krasilnikov *et al*., 2013), and high levels of precipitation contribute to nitrate leaching. Early maize was brought to the Mexican highlands, where it diversified before spreading to the north and south (Matsuoka *et al*., 2002). In the highlands, it probably experienced lower temperatures (Eagles and Lothrop, 1994) and low-phosphorus due to the presence of Andisols, which have high P fixation (Bayuelo-Jiménez *et al*., 2011; Bayuelo-Jiménez and Ochoa-Cadavid, 2014). In other parts of the world, including the central United States, low soil nitrogen may have been a more significant limitation to maize growth (Schlegel and Havlin, 1995), especially before the widespread use of synthetic fertilizers in the 1940s.

Root architecture has implications for soil resource acquisition (Lynch, 1995; Lynch 2019). While phosphorus is highly immobile in soils, nitrate, the dominant form of inorganic nitrogen in most agroecosystems, is prone to leaching (Barber, 1995; Di and Cameron, 2002; Kabala *et al*., 2017). In the absence of rainfall, soil water may also become relatively more available at depth due to evaporation at the soil surface as well greater root activity in shallow soil (Asbjornsen *et al*., 2008). Nitrogen leaching is affected by soil texture and precipitation levels, among other factors (Gaines and Gaines, 1994; Dodd *et al*., 2000). Therefore, while maize root angles become steeper relative to the soil surface under low-nitrogen conditions (Trachsel *et al*., 2013), the ideal root angle for nitrogen capture depends on the environment (Dathe *et al*., 2016). Tradeoffs exist between root architectures that improve the acquisition of phosphorus and those useful for mobile resources like nitrates and water (Ho *et al*., 2005; Rangarajan *et al*., 2018). For example, low crown root number is useful for the acquisition of nitrogen (Saengwilai *et al*., 2014) and water (Gao and Lynch, 2016), but high crown root number improves phosphorus capture (Sun *et al*., 2018) in maize when those resources are limited. Similarly, low lateral root branching density improves maize growth with low nitrogen (Zhan and Lynch, 2015) and water (Zhan *et al*., 2015), but high lateral root branching density improves phosphorus acquisition (Jia *et al*., 2018).

While increased seminal root number improves phosphorus capture (Zhu *et al*., 2006), it is unclear how the increase in seminal root number associated with domestication influences seedling nitrogen acquisition. At the beginning of the growing season, when seminal roots are likely to be the most important, nitrogen concentrations may be greater in the topsoil if inorganic nitrogen fertilizer has recently been applied (Jobbágy and Jackson, 2001). In the corn belt region of the United States, nitrogen mineralization from organic matter may also be greater in the topsoil at the time of planting due to organic matter and soil temperature gradients (Gupta *et al*., 1982). Therefore, the distributions of nitrogen and phosphorus in the soil profile may be similar during early growth. The ‘steep, cheap, and deep’ maize root ideotype proposes two phenotypes for seminal roots in order to complement the nodal roots (Lynch, 2013). If the first crown roots are slow to develop, seminal roots should grow at shallow angles relative to the soil surface and be highly branched in order to capture shallow soil resources. If the first crown roots develop quickly and are sufficient for capturing shallow soil resources, seminal roots should grow at a steeper angle and have less lateral branching in order to capture deep resources. Therefore, it is possible that0020seminal roots perform multiple functions for domesticated maize.

Functional-structural plant models, which combine three-dimensional representations of plant architecture with physiological models, can provide useful insights into systems that are challenging to study empirically (Vos *et al*., 2010; Dunbabin *et al*., 2013). Understanding the influences of domestication on seminal root number in maize requires the consideration of plant performance in diverse environments and phenotypes intermediate to those of maize and teosinte. Since maize and teosinte differ in several respects, including vigor, tillering, and growth habit, it is challenging to understand how individual components of their phenotypes contribute to stress adaptation. Simulation modeling can be a useful approach to understanding maize and teosinte root architectures because it allows traits to be experimentally modified in isolation while other components of the phenotype remain constant. The functional-structural plant model *OpenSimRoot* includes a detailed root architectural model that accounts for root construction costs, respiration, and nutrient uptake at the level of individual root segments (Postma *et al*., 2017). It also allows for the simulation of low-phosphorus soils and for soil nitrate leaching and depletion to be simulated in three dimensions.

This study aims to understand the impact of maize seminal roots on seedling nitrogen acquisition and examine how maize and teosinte phenotypes might have caused domesticated maize to have a greater number of seminal roots than its wild ancestor. We propose that increased seminal root number improves nitrogen acquisition by maize seedlings, and the large increase in seed size associated with domestication might have facilitated an increase in seminal root number by providing additional resources for growth during the early development of the plant.

## Materials and Methods

The functional-structural plant model *OpenSimRoot* (Postma *et al*., 2017) was used to evaluate the impacts of seminal root number and other components of maize and teosinte phenotypes on nitrogen and phosphorus acquisition. Model version 2e4779cf was used, which is publicly available (https://gitlab.com/rootmodels/OpenSimRoot). *OpenSimRoot* uses the SWMS_3D model of water and solute movement (Šimunek *et al*., 1995) to simulate nitrate leaching with a finite element approach. Organic matter mineralization follows the Yang and Janssen model (Yang and Janssen, 2000). Ammoniacal nitrogen is not modeled, as nitrification is generally rapid in the field conditions being simulated (Barber, 1995). Phosphorus uptake is simulated using the Barber-Cushman model (Barber and Cushman, 1981). Roots are composed of connected cylinders or truncated cones that are generally less than one centimeter in length. Root segment construction requires carbon and nutrients, and respiration and nutrient uptake are calculated at the root segment level. Root cortical aerenchyma formation occurs after root construction, and it results in decreased respiration rates and the remobilization of nutrients (Postma and Lynch, 2011 *a*, *b*). Distinct root classes are simulated that have individual branching angles, diameters, and growth rates, among other things. Shoots are simulated non-geometrically, but a leaf area parameter is included from which light interception and photosynthesis are calculated. Nutrient stresses have defined impacts on the rate of leaf area expansion, photosynthesis, and root growth rates. Carbon is partitioned among roots, shoots, and leaves according to allocation coefficients.

The existing *OpenSimRoot* maize model uses plant parameters that are based on empirical measurements, and some of the parameters include stochasticity (Postma and Lynch, 2011 *a*). The existing ‘Rocksprings’ and ‘WageningseBovenBuurt’ low-nitrogen environments were used in this study as the silt-loam and sand-textured soils, respectively (described in Dathe *et al*., 2016). The ‘Rocksprings’ environment is parameterized to resemble the Russell E. Larson Agricultural Research Center in Rock Springs, Pennsylvania, USA, and the ‘WageningeseBovenBuurt’ environment is based on the ‘De Bovenbuurt’ research field at Wageningen University, the Netherlands. Bulk density and parameters related to soil water retention are different for the two soils, while the same precipitation data and initial concentrations of nitrate by depth, which were measured in Rock Springs in 2009, were used for both environments. Models begin with nitrate being concentrated close to the soil surface, and nitrate leaching and depletion take place over time.

### Model Parameterization

Seeds of PI 213706 (*Zea mays* ssp. *mays*) and Ames 21803 (*Zea mays* ssp. *parviglumis*) were acquired from the United States Department of Agriculture North Central Regional Plant Introduction Station, Ames, Iowa, USA. Ames 21803 was collected in the Mexican state of Guerrero, and its root architecture, anatomy, and vigor are fairly representative of other *parviglumis* accessions (Burton *et al*., 2013 *a*). PI 213706 is an open-pollinated Midwestern dent landrace sold commercially in the 1920s as Osterland Reid’s Yellow Dent. It is a strain of Reid’s Yellow Dent, which was the most popular cultivar in the corn belt region of the United States around the beginning of the 20th century (Troyer, 1999). Osterland Reid’s Yellow Dent is also the parent of many notable maize inbred lines and contributes about 15% of the background of US maize hybrids (Troyer, 2004). Average seed mass was determined for both genotypes, and the fruitcase was removed from 10 representative teosinte seeds to determine the average kernel weight.

Plants were grown in a growth chamber (Environmental Growth Chambers, Chargin Falls, OH, USA) for 14 h days with 27°C day and 17°C night temperatures. Approximately 600 µmol m^-2^ s^-1^ photosynthetically active radiation (PAR) were provided by metal halide bulbs with a photoperiod of 14 h. Plants were grown in 1.2 x 0.15 m circular polyvinyl chloride mesocosms lined with plastic. The growing media consisted of 50% commercial sand (US Silica, Berkley Springs, WV, USA), 30% vermiculite, 5% perlite, 15% field soil (Hagertown silt loam, mesic typic Hapludalf) collected from the Russell E. Larson Agricultural Research Center, and 60 g Osmocote Plus Fertilizer (Scotts Miracle-Gro Company, Marysville, OH, USA). Seeds were planted at a depth of 3 cm and irrigated with tap water as needed.

Four plants of each genotype were destructively harvested 5, 8, 10, 15, 20, and 25 days after planting. Leaf area was measured using a LI-3100 Area Meter (LI-COR Biosciences, Lincoln, NE, USA). Basal segments of primary, seminal, and nodal roots were excised for respiration measurements, and lateral roots were removed from those segments using a razor blade. Root respiration was measured using a LI-6400 Portable Photosynthesis System (LI-COR Biosciences) and a custom root chamber by recording carbon dioxide evolution every 5 s for 180 s. Respiration measurements were taken in a climate-controlled room at 22°C. Apical and basal root segments preserved in 75% ethanol were sectioned using laser ablation tomography (Hall and Lanba, 2019; Strock *et al*., 2019), and root diameter was determined from the resulting images using the ObjectJ plugin for ImageJ (Schneider *et al*., 2012). Root angle for each whorl was measured on separate plants grown in 10 L pots with three replications. Mesocotyl diameter and growth rate were also measured in separate plants that were grown in tree pots 25.4 cm in height and 6.35 cm in diameter (Stuewe & Sons, Tangent, OR, USA) by destructively harvesting four seedlings every day for six days. Existing *OpenSimRoot* maize parameters were modified to include the measured root angle for each whorl, branching frequency for each root class, seminal root number, nodal root number per whorl, axial root growth rate for each root class, leaf area expansion rate, mesocotyl growth rate and diameter, root respiration rate, root diameter for each root class, time of root emergence for each whorl of nodal roots, and kernel size for both subspecies. Branching frequency values are drawn randomly from a uniform distribution in the existing *OpenSimRoot* maize model, so a distribution of the same range was centered around the average measured branching frequency value for each subspecies. Growth rates were calculated by fitting second order polynomial regressions to the measured data and taking the first derivative. At 25 days after planting, teosinte seedlings had two small tillers at most and no tiller roots, so tiller leaf area was considered to be part of the main plant, and tiller formation was not simulated explicitly.

Parameters representing the amounts of seed nitrogen and phosphorus available for seedling growth are included in *OpenSimRoot*. Rather than measuring total seed N and P, which might include nutrients contained in structural compounds that cannot be remobilized, tissue nutrient content was measured on seedlings that were grown for three weeks in germination paper (Anchor Paper, St. Paul, MN, USA) partially submerged in 5 mM calcium sulfate solution. At the time of sampling, seedlings exhibited nutrient deficiency symptoms and it was assumed that seed nutrient reserves had been exhausted. For nitrogen content, root and shoot tissue was collected from six seedlings per subspecies, and the tissue was dried, ground, and homogenized. N was quantified in approximately 2 mg subsamples using a CHN elemental analyzer (EA2400, PerkinElmer, Waltham, MA, USA). For phosphorus content, four samples per subspecies of approximately 60 mg were digested with nitric acid, and P was measured using inductively coupled plasma optical emission spectrometry (Avio 200, PerkinElmer). Because of the larger tissue mass required for P analysis and limited seed availability, *parviglumis* seeds from multiple sources were used for that measurement.

*OpenSimRoot* includes multiple functions that can be used to simulate the process by which seed carbohydrate reserves become available to the growing seedling. The function commonly used by the maize model calculates seed carbohydrate reserves as a proportion of seed mass and provides those reserves to the developing seedling as demanded by carbon sinks. To account for the changes associated with maize domestication, the carbon module was modified to add a seed carbohydrate content parameter (code available at https://gitlab.com/AP2003/opensimroot-seed-carbohydrate-content). Values for the average kernel carbohydrate content of *parviglumis* accessions and maize landraces reported in Flint-Garcia *et al*. (2009) were used.

Other model parameters were thought to be fairly conserved through domestication or of less importance in 25-day-old seedlings than older plants. Since seminal roots were not observed frequently enough in teosinte for parameterization, the maize seminal root parameters were used in the teosinte model for simulations involving the potential utility of seminal roots to teosinte. Some differences between maize and teosinte are shown in Table S1, and full parameterization files are provided [**Supplementary Information**].

### Simulated Environments

Except when stated otherwise, models simulated a single maize plant grown in a monoculture at the ‘Rocksprings’ environment by simulating a rectangular prism of soil 60 cm in width, 26 cm in length, and 150 cm in depth, and roots that touched the vertical edges of the simulated area were reflected back in order to account for roots of neighboring plants. Different rates of available soil nitrogen were obtained by multiplying the initial measured concentrations of nitrate and organic matter at each soil depth by a constant as described by Dathe *et al*. (2016). Phosphorus rates were adjusted by changing phosphorus concentrations by depth in the same way. Planting density treatments were created by maintaining commercial row spacing of 76.2 cm and changing the in-row spacing.

### Data Analysis

Data were analyzed using R version 3.6.2 (R Core Team, 2019) using the packages ‘Biobase’ (Huber *et al*., 2015) and ‘tidyverse’ (Wickham *et al*., 2019). Spatial data were processed with Paraview 5.5.2 (Ahrens *et al*., 2005).

### Statistics

Several *OpenSimRoot* model parameters are sampled from distributions using a random number generator, which causes stochasticity in model runs. At least six replications with different random number generator seeds were performed for all treatments, and raw data are shown as appropriate to demonstrate when treatment effects are larger than those due to stochasticity. Where the figure format precludes the display of raw data, averages of at least six replications are shown. Performing statistical hypothesis testing on model results of this type is not appropriate because statistical power is somewhat arbitrary and knowledge of the model makes the null hypothesis false *a priori* (White *et al*., 2014; Postma *et al*., 2014).

### Recombinant Inbred Line (RIL) Experiment

To examine the relationship between seed mass and seminal root number in maize inbred lines, seeds of 60 members of a B73 x Mo17 recombinant inbred line (RIL) population (Kaeppler *et al*. 2000) and 95 members of a Ny821 x H99 RIL population (Johnson *et al*., 2016) were acquired from Dr. Shawn Kaeppler (University of Wisconsin, Madison, WI, USA). Average kernel mass was determined based on ten seeds, and three seeds per genotype were germinated and transferred to pouches consisting of germination paper (Anchor Paper) covered with black polyethylene and suspended in nutrient solution as described by Hund *et al*. (2009). The nutrient solution consisted 32 of (in µM) 1000 NH_4_NO_3_, 125 MgSO_4_, 3000 KH_2_PO_4_, 500 CaCl_2_, 12.5 H_3_BO_3_, 1 MnSO_4_·2H_2_O, 1 ZnSO_4_·7H_2_O, 0.25 CuSO_4_·5H_2_O, 1.5 Na_2_MoO_4_·2H_2_O, and 100 Fe-DTPA. The pH was adjusted to 6.0 daily using 3N KOH. Seedlings were grown in fluorescent light racks with a photoperiod of 14 h and at a temperature of 22°C. Seminal root number was recorded 14 d after planting.

## Results

### Spatiotemporal nature of nutrient acquisition by seminal roots in domesticated maize

Maize landrace models had larger and deeper root systems than those of teosinte at 25 days after planting (Fig. 1). During that time, seminal roots and their laterals were responsible for approximately 35 percent of the nitrogen and phosphorus acquired by maize in field conditions with 50 kg ha^-1^ available nitrate and 2 kg ha^-1^ available phosphorus (Fig. 2). Lateral roots acquired more phosphorus than axial roots, while axial roots acquired more nitrogen than their laterals during the first 25 days of growth. N and P capture by nodal roots is likely to overtake that of seminal roots after that time. Because seminal roots emerged before nodal roots, they were able to acquire nitrogen from deeper in the soil profile than nodal roots at 15 days after planting (Fig. 3). By 25 days after planting, the contributions of seminal and nodal roots to nutrient acquisition by depth were more similar. In teosinte, nodal roots acquired a larger percentage of the total nitrogen and phosphorus than they did in maize (Fig. S1), and, like maize, the primary root was able to acquire deeper nitrogen than the nodal root system at 15 days after planting (Fig. S2).

**Fig 1.**
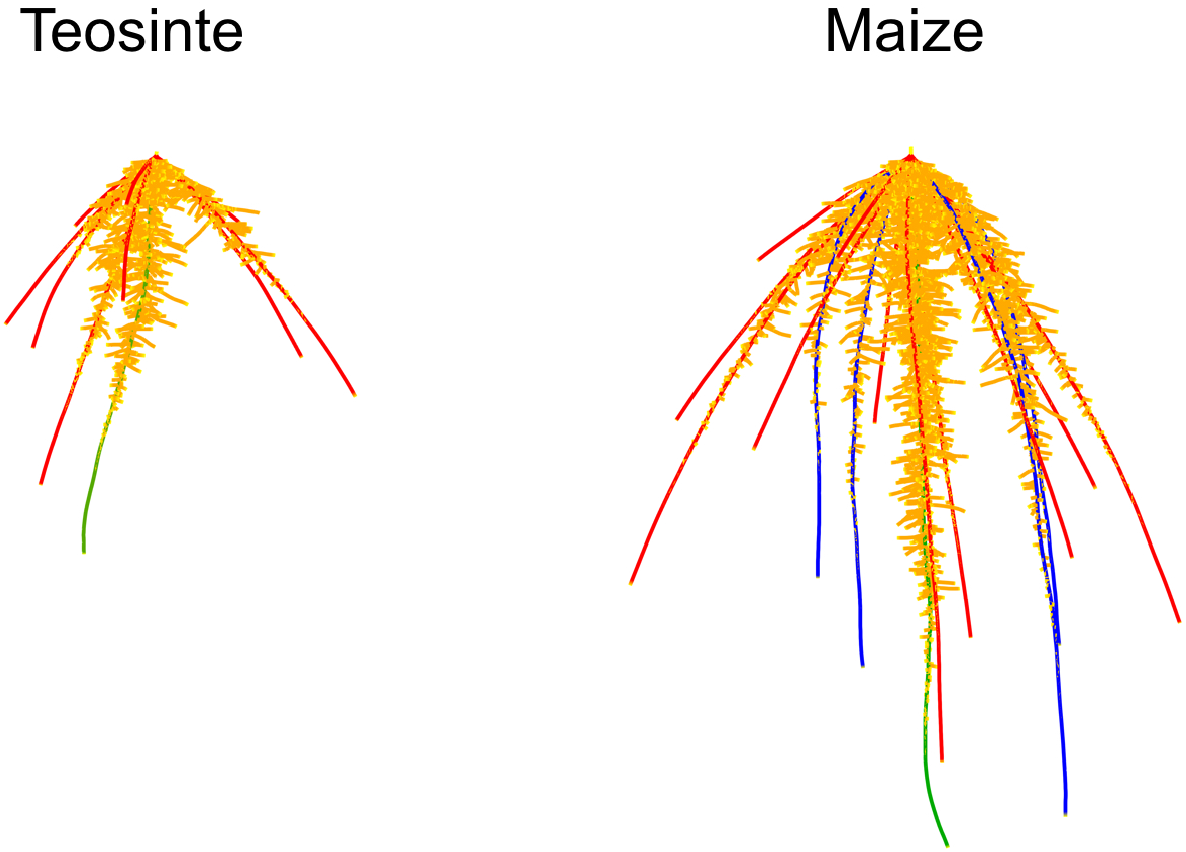
Graphical renderings of the teosinte and maize root system models at 25 days after planting when grown without nutrient stress. Roots are dilated for clarity. Root classes are colored as follows: blue, primary root; green, seminal roots; red, nodal roots; orange, lateral roots.

**Fig 2.**
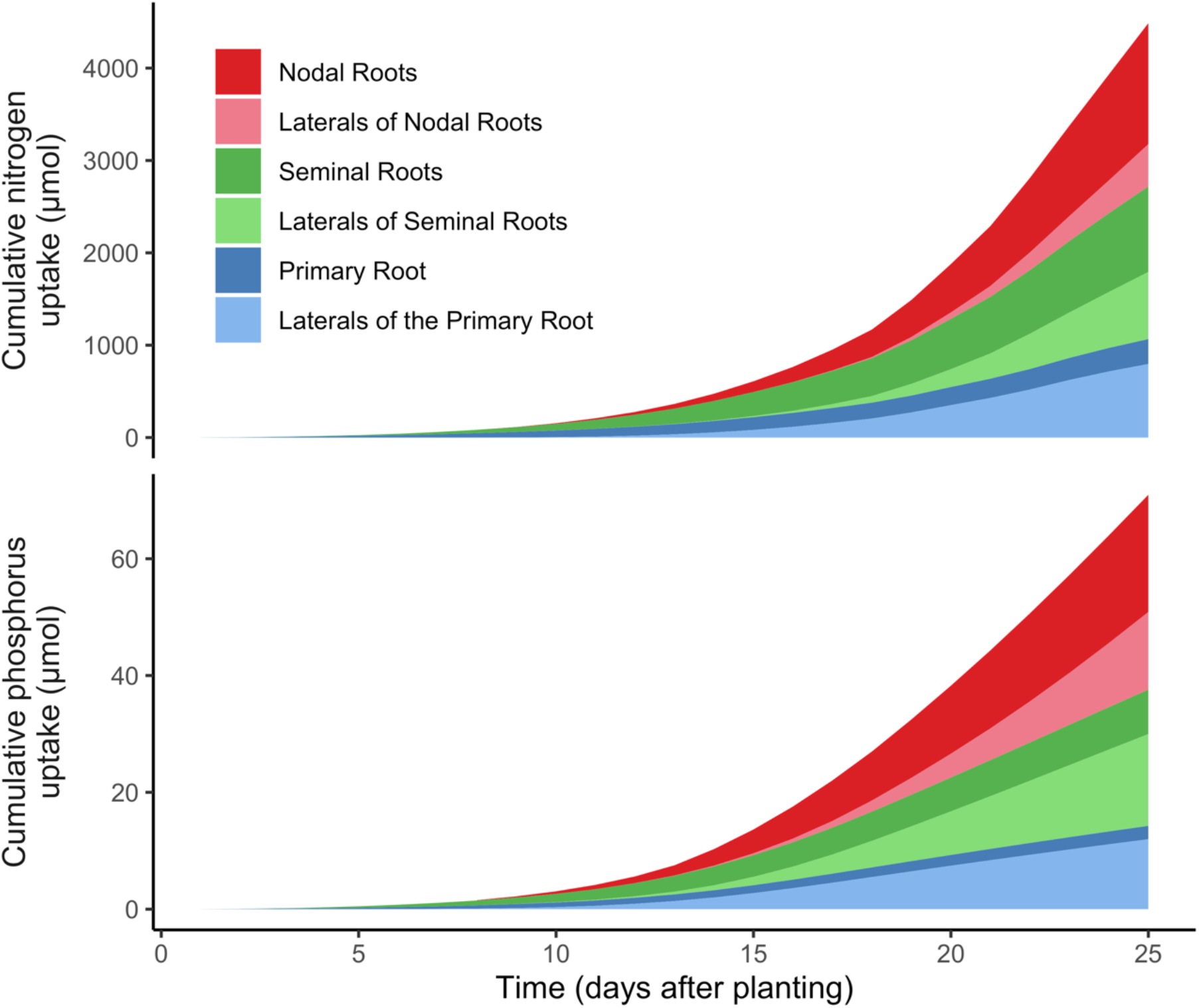
The cumulative contribution of each root class of a simulated maize landrace plant to nutrient acquisition in field conditions with 50 kg ha^-1^ available nitrate (top) and 2 kg ha^-1^ available phosphorus (bottom). These levels of nutrient availability lead to an approximately 50% reduction in shoot dry weight at 25 days after planting. The values presented are averages from six model replications that include stochasticity. Nodal roots emerged seven days after planting.

**Fig 3.**
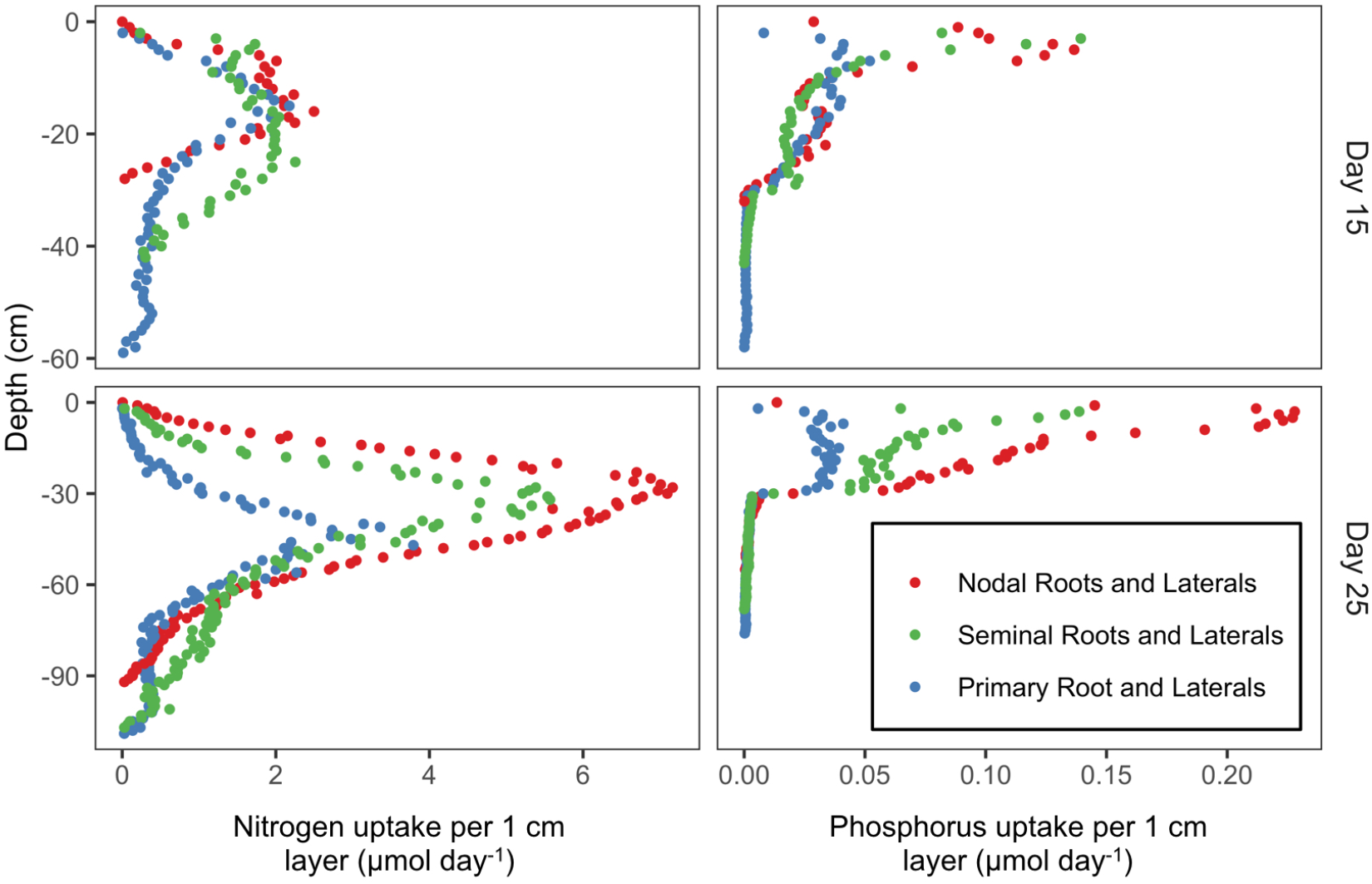
The contributions of each root class of a maize landrace to the acquisition of nitrogen and phosphorus by depth on the 15th and 25th days following planting in field conditions with 50 kg ha^-1^ available nitrate (left) 2 kg ha^-1^ available phosphorus (right). Root segments are binned by 1 cm soil layers, and the values presented are averages from six model runs that include stochasticity.

### Utility of seminal roots to domesticated maize in different environments

Since nitrate is highly mobile in the soil, its location over the course of a growing season depends on precipitation and soil texture, among other factors (Gaines and Gaines, 1994; Dodd *et al*., 2000). Therefore, the utility of root phenotypes for nitrate capture may depend on the environment (Dathe *et al*. 2016). However, seminal roots were beneficial for nitrogen acquisition with two different soil textures in all the precipitation regimes and nitrate fertilization rates that were simulated (Fig. 4). Precipitation was inversely related to plant nitrogen acquisition. Increased seminal root number also improved phosphorus acquisition and plant growth in a simulated low-P environment (Fig. 5), which has been previously reported by a field study (Zhu *et al*., 2006).

**Fig 4.**
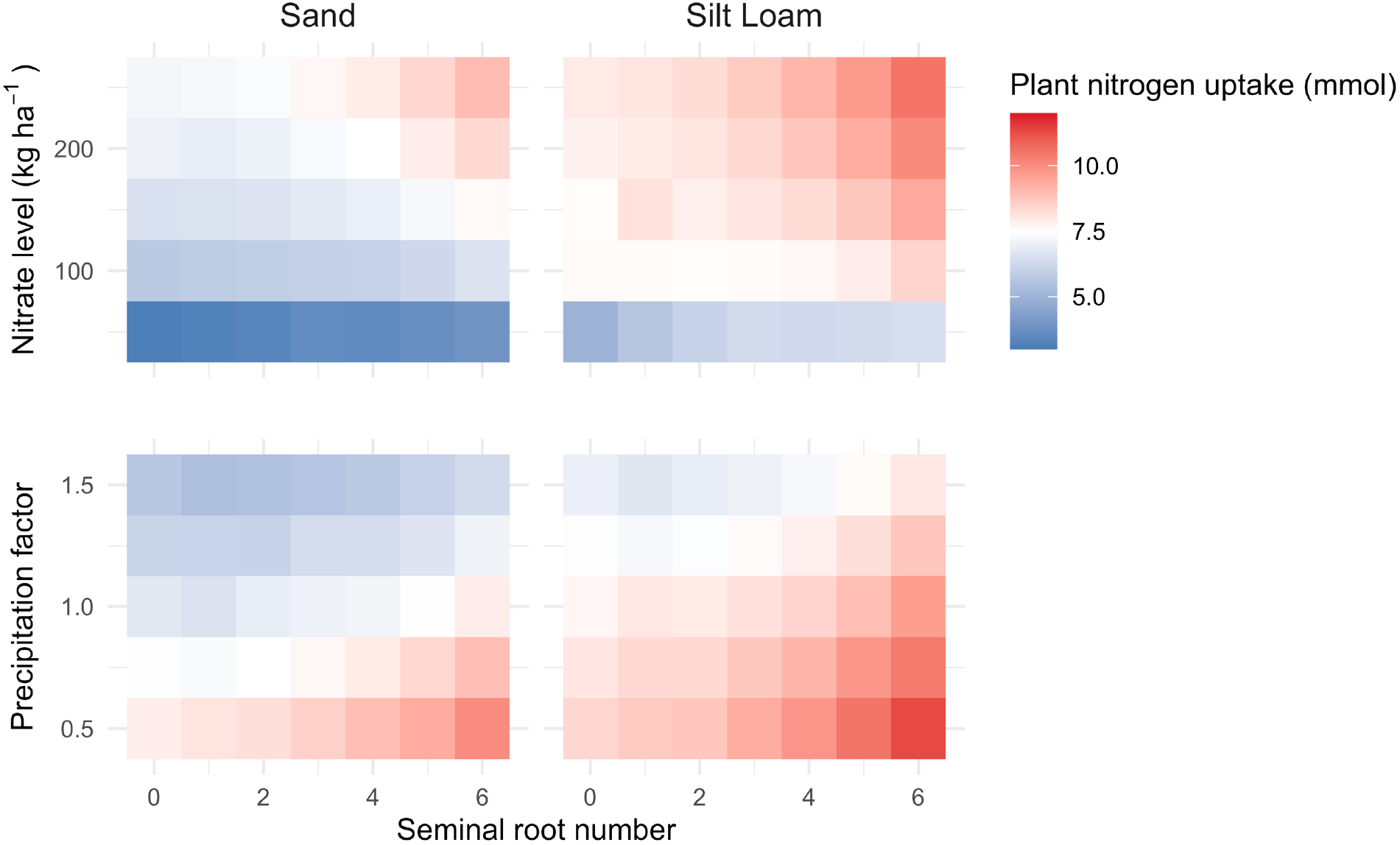
The nitrogen acquisition of a maize landrace with varied seminal root number at 25 days after planting in environments with different initial nitrate concentrations, precipitation regimes, and soil textures. Precipitation treatments were created by multiplying the measured daily rainfall by a constant. The values presented are averages of replicated model runs that include stochasticity.

**Fig 5.**
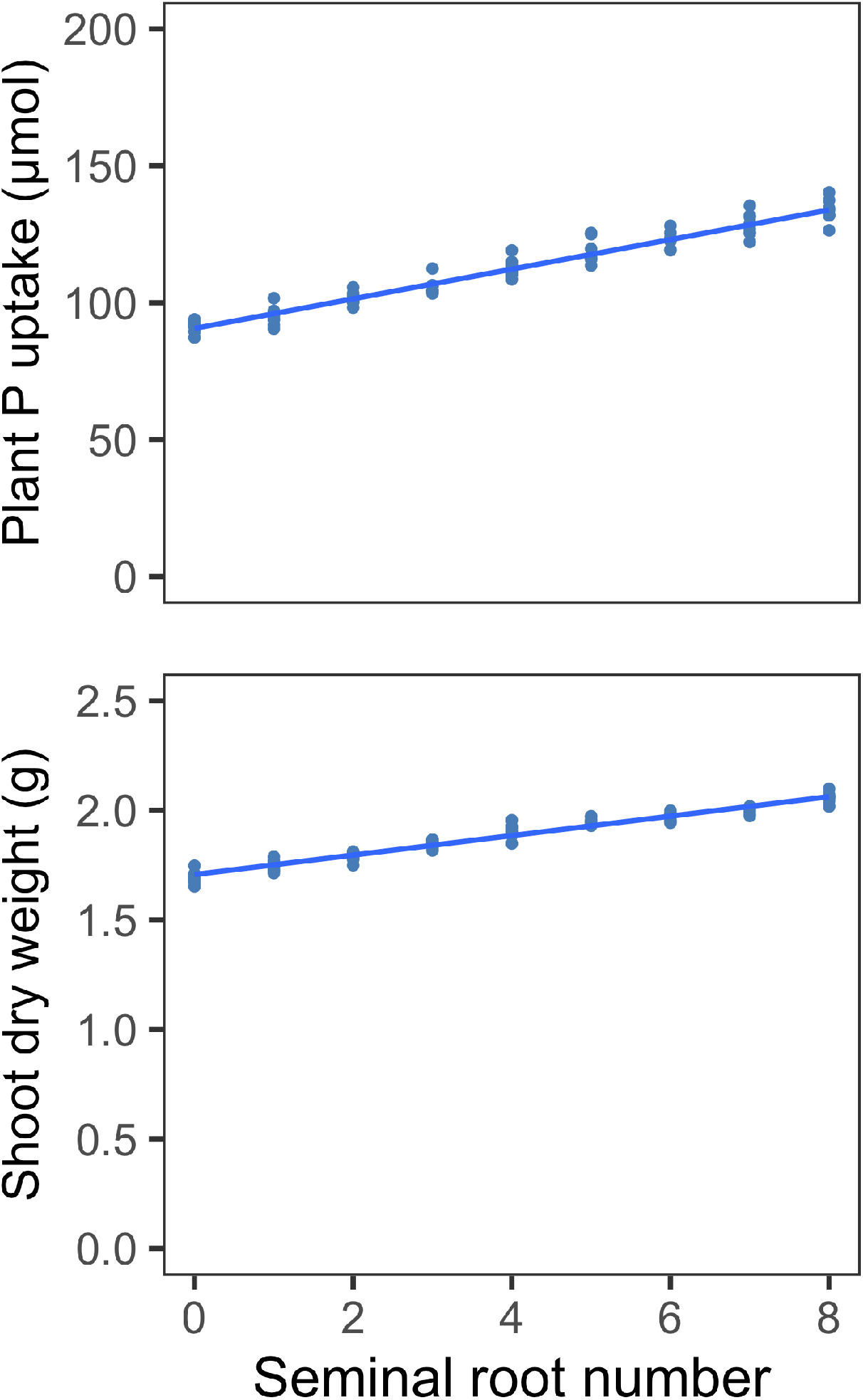
The impact of seminal root number on phosphorus acquisition (top) and shoot biomass (bottom) of a maize landrace in a soil with 2 kg ha^-1^ available phosphorus at 25 days after planting. Points represent values from individual model runs.

The presence of seminal roots decreases the average distance to the nearest root of a neighboring plant (Fig. 6), which might have implications for interplant competition. However, increasing planting density from 37,050 plants per hectare to 111,150 plants per hectare reduced plant nitrogen acquisition by less than 1 mmol in the first 25 days after planting. Seminal roots improved nitrogen acquisition and seedling growth for all planting density treatments.

**Fig 6.**
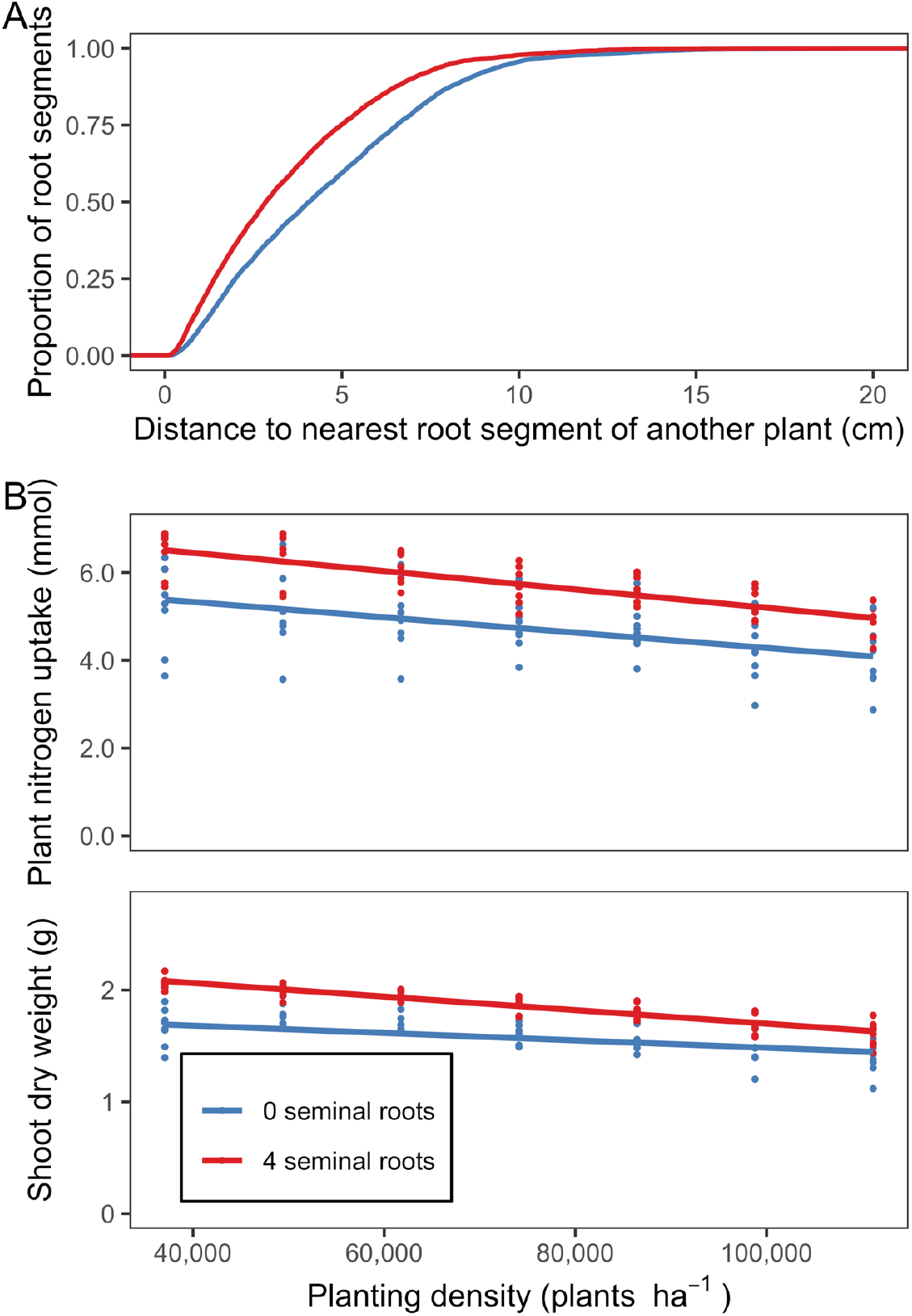
The impact of planting density on the utility of seminal roots. (A) Cumulative distribution functions of the distance from a randomly selected root segment of a maize landrace to the nearest root segment of a neighboring plant at a planting density of 74,100 plants per hectare (close to commercial planting density) at 25 days after planting. Root position data from six model runs per treatment were subsampled. Nutrient stress is not included in order to avoid confounding allometric effects. On average, the nearest root of a neighboring plant is closer at 25 days after planting if seminal roots are formed. (B) Nitrogen acquisition and shoot dry weight of a maize landrace at 25 days after planting in a soil with 50 kg ha^-1^ available nitrate. Points represent values from individual model runs.

### Utility of seminal roots to teosinte with low-N and low-P

While increased seminal root number improved plant growth and nutrient acquisition in the maize landrace model in low-N and low-P environments, forming two or more seminal roots left teosinte with insufficient resources to germinate (Fig. 7). The utility of seminal roots to maize decreased when the landrace was simulated as having the growth rates of teosinte due to the decreased nutrient demands of the smaller plant. When teosinte was simulated as having the amount of seed nitrogen and phosphorus available from maize seeds, it was able to grow larger, but the optimal number of seminal roots was not impacted. When it was simulated to include the carbohydrate reserves that are present in maize seeds, however, the optimal number of seminal roots was increased.

**Fig 7.**
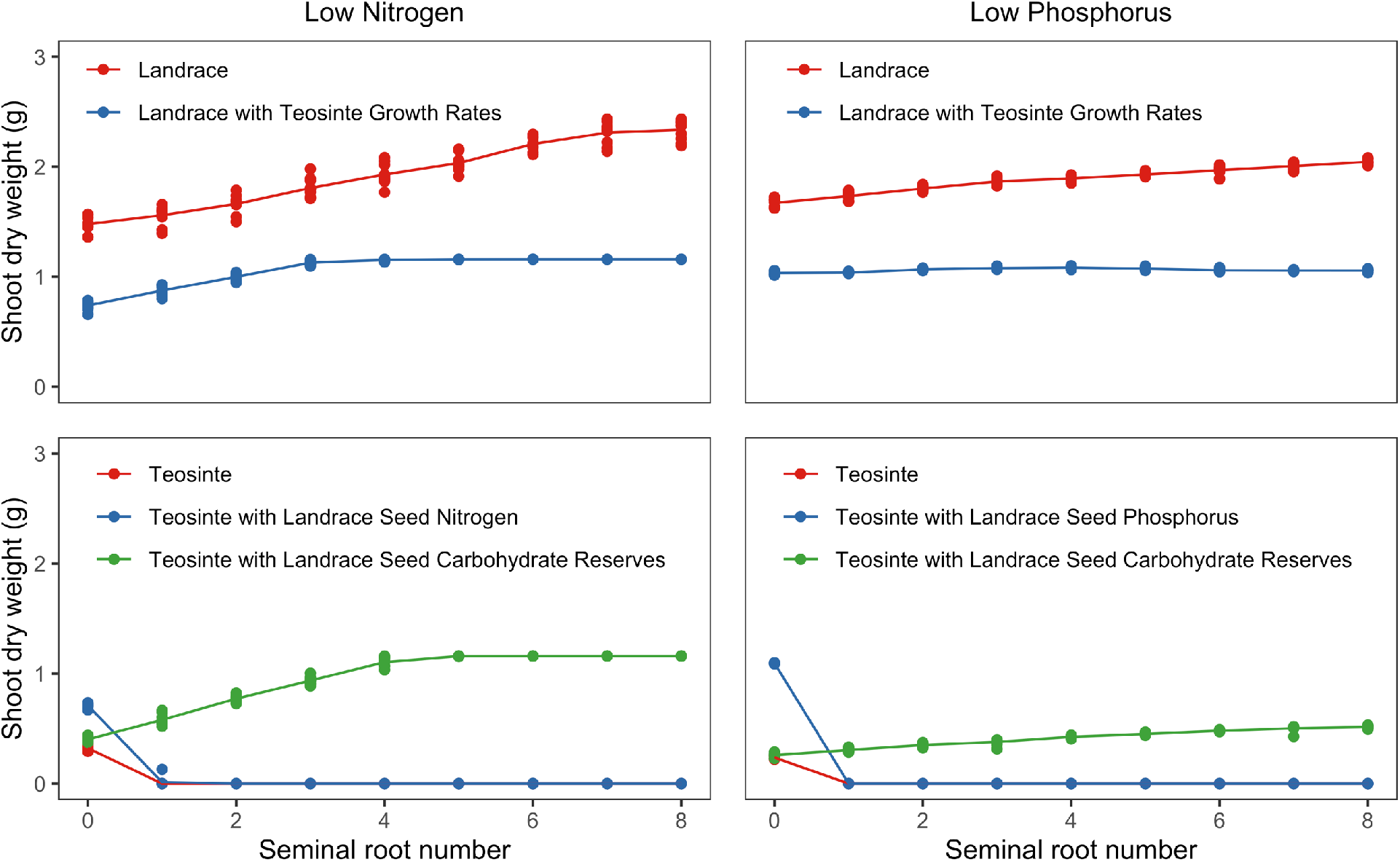
The utility of seminal roots to a maize landrace, teosinte, and three composite phenotypes in a low-N soil with 50 kg ha^-1^ available nitrate (left) and a low-P soil with 2 kg ha^-1^ available P (right). Points represent individual model runs, and trendlines pass through the average value for each treatment. Teosinte treatments with shoot dry weights of approximately zero did not have the seed resources necessary for germination.

Assuming that seed carbohydrate content as a proportion of seed mass is constant, simulations of maize seeds intermediate in size to those of teosinte and the maize landrace suggest that the optimal seminal root number for plant growth under low-N stress may be dependent on seed carbohydrate reserves when seeds are smaller than approximately 125 mg (Fig. 8). Seminal root number did not appear to be correlated with seed size in two populations of maize recombinant inbred lines in which all lines had seeds of mass greater than 125 mg on average (Fig. S3). The average kernel mass of teosinte is about 30 mg while the average kernel mass of maize landraces is approximately 280 mg, and carbohydrate content as a proportion of seed mass is lower in teosinte than landraces (Flint-Garcia *et al*., 2009). Some Poales that have seeds intermediate in mass to those of maize and teosinte, such as barley, oat (*Avena sativa*), rye (*Secale cereale*), and wheat, form a variable number of seminal roots, while other species with seeds of less than 30 mg on average, such as pearl millet (*Cenchrus americanus*), rice, and sorghum, do not form seminal roots (Fig. 9). Variation in seed composition and carbon partitioning, among other things, might change the relationship between seed mass and the optimal number of seminal roots in other species, however.

**Fig 8.**
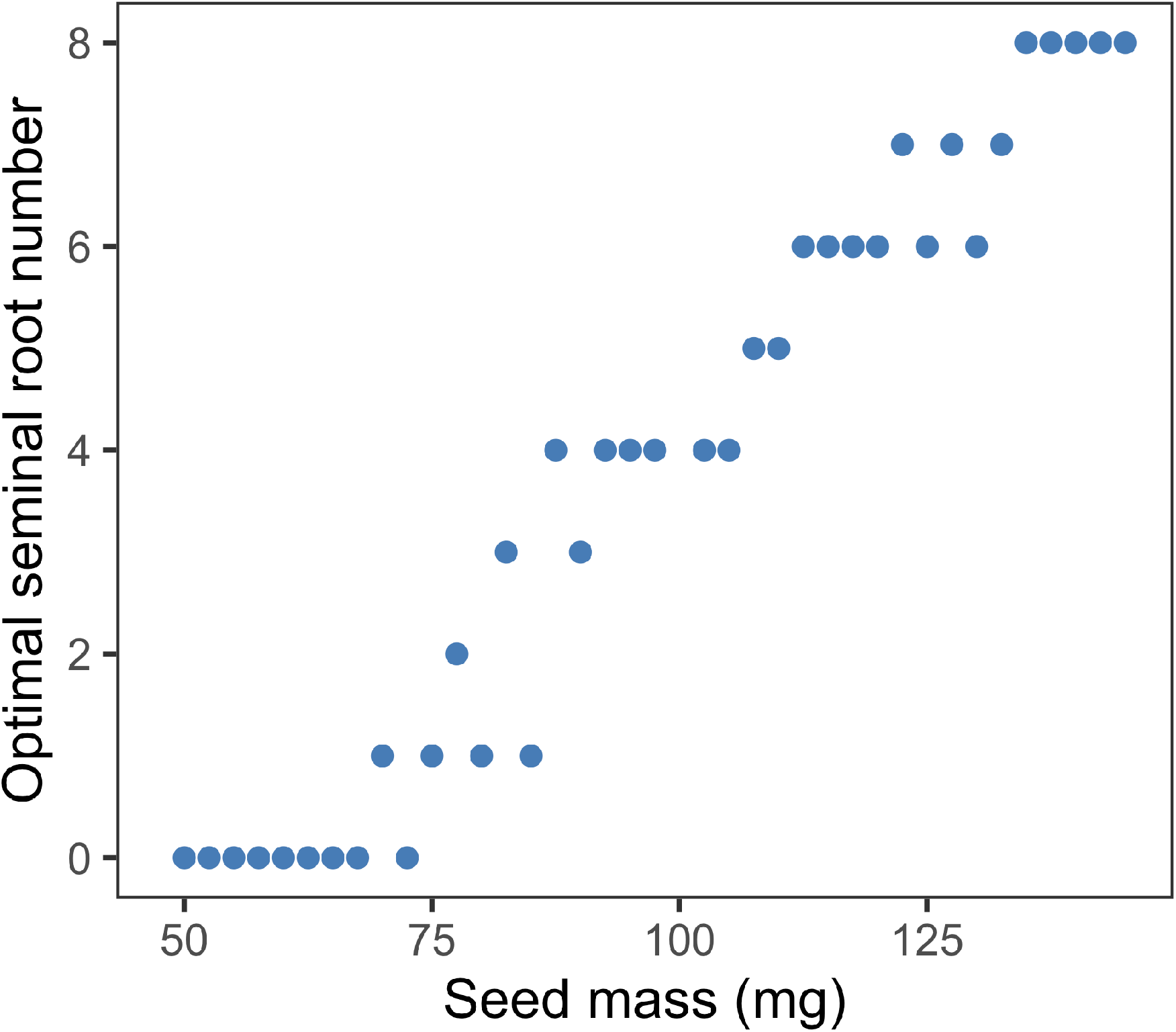
The relationship between seed mass and optimal seminal root number in a low-N environment with 50 kg ha^-1^ available nitrogen if seed carbohydrate content as a proportion of seed mass and seed N reserves are constant (the maize landrace parameters were used). Optimum values were determined based on replicated model runs that include stochasticity.

**Fig 9.**
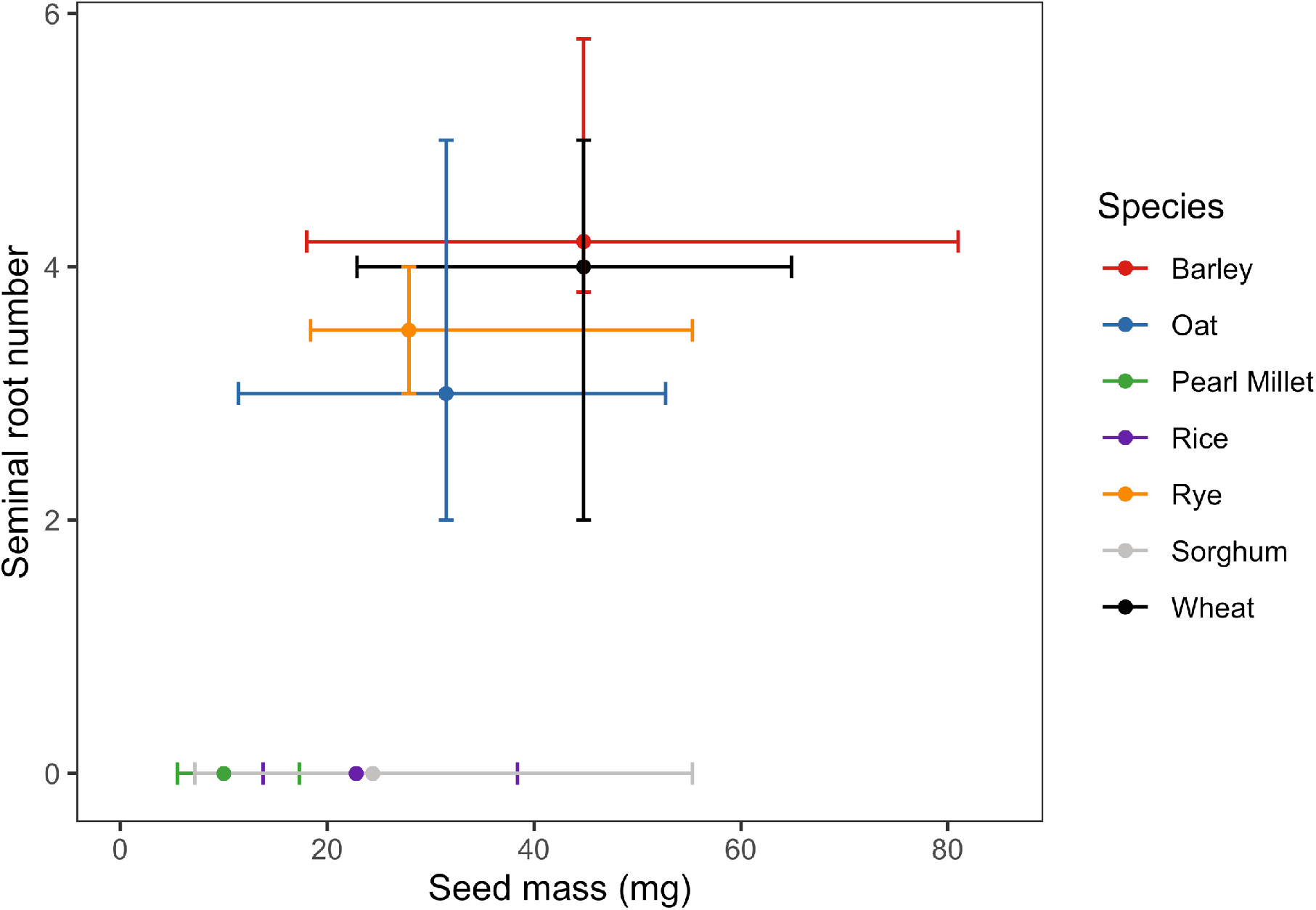
Variation in seminal root number and seed mass in domesticated barley, oat, pearl millet, rice, rye, sorghum, and wheat based on the literature and germplasm repository records. Following the definition of seminal roots commonly applied to maize (Hochholdinger *et al*., 2004), the radicle is not considered a seminal root. The mean, minimum, and maximum seed mass values are calculated using all landraces for which data are available from the United States Department of Agriculture National Plant Germplasm System. Sources for the seminal root number values of domesticates are as follows: Barley, Grando and Ceccarelli, 1995; Oat, Schuurman and Boer, 1970; Pearl Millet, Passot *et al*., 2016; Rice, Hochholdinger *et al*., 2004; Rye, Pavlychenko and Harrington, 1934; Sorghum, Singh *et al*., 2010; Wheat, Golan *et al*., 2018.

As a result of the differences in seed mass and carbohydrate content between maize and teosinte, a much greater proportion of the starting seed carbohydrate reserves were depleted by seven days after planting in teosinte than in maize (Fig. S4). Low-nitrogen and low-phosphorus stress, however, were not experienced until at least 10 days after planting (Fig. S4). Due to the carbohydrate limitation, tradeoffs may also exist between increased seminal root number and the growth of the primary root. In teosinte, the presence of additional seminal roots decreased the average length of the primary root and the length of the longest seminal root at seven days after planting (Fig. 10).

**Fig 10.**
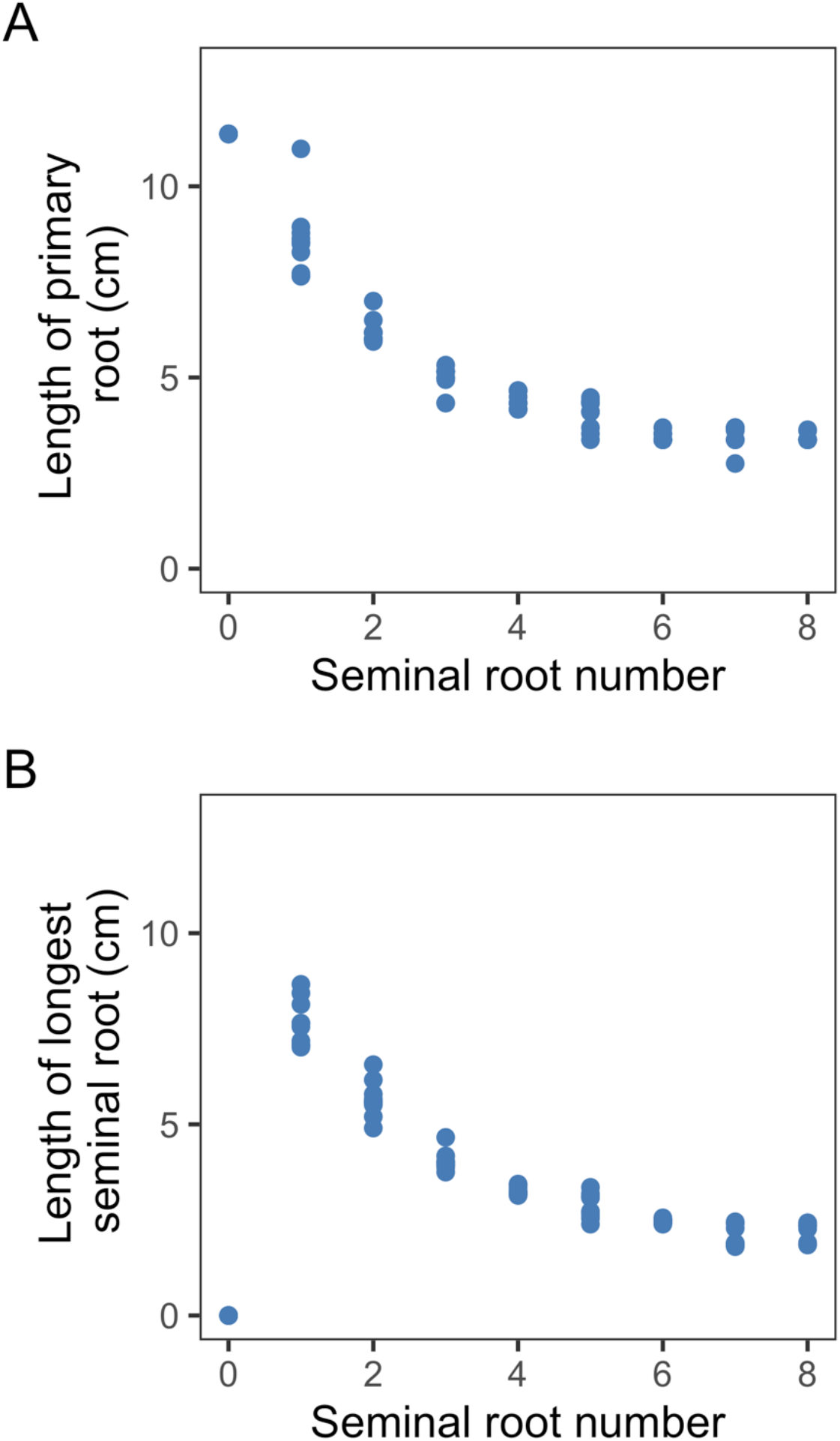
The impact of seminal root number on the length of the primary root and longest seminal root in teosinte at seven days after planting. This tradeoff exists due to the limited seed carbohydrate reserves of teosinte. Points are values from individual model runs.

## Discussion

*OpenSimRoot* was used to understand the spatiotemporal dynamics of nutrient acquisition by the seminal roots of maize, the utility of seminal roots to maize in different low-nutrient environments, and the degree to which differences in the growth and morphology of maize and teosinte impact the optimal seedling root architecture of each subspecies. The results suggest that seminal roots are beneficial for both nitrogen and phosphorus acquisition during the development of maize seedlings, and seminal roots can improve nitrogen acquisition in environments with several different precipitation regimes, fertilization rates, and soil textural classes. High numbers of seminal roots may not be beneficial to teosinte because its small seeds have smaller carbohydrate reserves to support seedling growth, and the lower growth rates of teosinte mean that it has lower nutrient requirements as a seedling. The relationship between increased seminal root number and improved phosphorus acquisition has previously been reported (Zhu *et al*., 2006), but the benefit of seminal roots in low-nitrogen soils is a new finding. Tradeoffs sometimes exist between nitrogen and phosphorus acquisition because phosphorus is generally concentrated in the topsoil and is fairly immobile (Lynch and Brown, 2001), while nitrate, the dominant form of nitrogen in most agricultural soils, is mobile (Wiersum, 1962). Growers often apply nitrogen fertilizer just prior to planting in order to reduce losses to leaching (Vetsch and Randall, 2004). Therefore, acquisition of both resources may be improved by high seminal root number because both resources are abundant in the topsoil during seedling establishment, while by flowering, shallow nitrogen is more likely to have been leached or depleted. Soil organic matter also tends to be concentrated in the topsoil (Rethemeyer *et al*., 2005), so mineralization may contribute to the availability of shallow nitrogen. Ammonium nitrogen is also likely to be concentrated in shallow soil because of its interactions with soil cation exchange sites.

While teosinte probably experiences interspecific competition for resources, modern maize production takes place in high-density, genetically uniform stands. Therefore, interplant root competition might contribute to fitness in the wild, while it may be undesirable in high-input agroecosystems (Denison *et al*., 2003). Root competition for phosphorus can be quantified by measuring the overlap of depletion zones because phosphorus primarily moves in the soil through diffusion (Rubio *et al*., 2001), while competition for mobile resources like nitrate and water is more dynamic. Nitrate is primarily acquired by plants through mass flow, and mass flow rates are related to the nutrient concentration in the soil and the rate of water uptake by the plant (Barber, 1995). Increasing planting density had only small impacts on plant nitrogen acquisition at 25 days after planting (Fig. 6), so seminal root number is not likely to have implications for interplant competition in agroecosystems. Some wild environments might have greater plant densities than those simulated here, however.

Seeds of maize landraces are approximately ten times larger than those of teosinte (Flint-Garcia *et al*., 2009). Small seed size may be adaptive for dispersal and population growth (Moles *et al*., 2005; Turnbull *et al*., 2012), while large seeds may be better for seedling establishment and stress tolerance (Wang *et al*., 2020). The presence of the cupulate fruitcase surrounding teosinte kernels 10 may have also constrained the seed size of the wild ancestors of maize, so selection for naked kernels may have preceded selection for increased seed size (Flint-Garcia, 2017). Seminal roots can emerge from maize seeds before photosynthesis has started and when the primary root is still small, so it may be advantageous for teosinte to apply all of its limited seed carbohydrate reserves to the growth of the radicle and coleoptile, while domesticated maize has more seed carbon, which can support optimal growth of the radicle and coleoptile in addition to seminal root development. It seems unlikely that the formation of seminal root primordia itself is constrained by the physical size of the seed because the primordia are quite small (Taramino *et al*., 2007).

Growth rates limit the utility of seminal roots to teosinte because it has a smaller nitrogen requirement as a seedling than maize due to its smaller size. The increase in seedling vigor associated with domestication may have occurred for several reasons. Genotypes that produce more secondary compounds to reduce herbivory often have lower growth rates, and herbivory may be a greater challenge in the wild than in managed agroecosystems (Paul-Victor *et al*., 2010; Züst *et al*., 2011). At the genome level, genome size appears to be inversely related to leaf elongation rate (Bilinski *et al*., 2018). Genome size is smaller in maize landraces than teosinte, but there is considerable variation (Díez *et al*., 2013). While growth rates and seed resources are parameters that can be separated in *OpenSimRoot*, maternal investment may also be biologically related to seedling vigor. Seed mass and seedling size are correlated in several species (Wulff, 1986; Zhang and Maun, 1990; Moegenburg, 1996), so the increased growth rates of maize relative to teosinte 29 may also have been a result of that change. Studies on the impacts of maize seed size on crop growth suggest that large seeds are associated with increased plant height in seedlings, but seed size is not related to yield or the size of the mature plant in non-stressed conditions (Kiesselbach, 1937; Hunter and Kannenberg, 1972).

Domesticated wheat and barley have more seminal roots than their wild progenitors, although the increase is smaller than the difference between maize and teosinte. Grando and Ceccarelli (1995) screened seminal root number in a small collection of modern barley cultivars, landraces, and wild *Hordeum spontaneum* accessions. Modern cultivars had seeds with an average mass of 49.4 mg and about 4.5 seminal roots on average, while wild *H. spontaneum* seeds were 28.5 mg on average and had an average of approximately 2.5 seminal roots. Average shoot dry weight was also greater on average in modern cultivars than *H. spontaneum*, but there was large variation within each germplasm group. While the same number of seminal root primordia are found in the seeds of diploid and tetraploid wild and domesticated wheat species, more primordia develop in the domesticates (Robertson *et al*., 1979; Golan *et al*., 2018). Grain weight is significantly greater in cultivated durum wheat (*T. turgidum* ssp. *durum*) than in wild emmer (*T. turgidum* ssp. *dioccoides*), but the difference is smaller than that observed in maize (Golan, 2015). While teosinte seeds overlap in mass with those of some species that frequently form seminal roots, such as wild barley, species differences in seed composition, growth rates, and other root traits might also influence the optimal seminal root number. For example, wheat seminal roots may be of smaller diameter than those of domesticated maize (Atwell, 1990; Hund *et al*., 2004), which would influence the construction and maintenance costs of each root. Sorghum offers an interesting comparison to maize because of the genetic and phenotypic similarity between the two species. Wild sorghum seeds are approximately 18 mg, while seeds of the domesticate may be as large as 58 mg (Wang *et al*., 2020). This is considerably smaller than most domesticated maize seeds, and seminal roots have not evolved in sorghum (Singh *et al*., 2010).

Considerable variation in seminal root number is present in maize landraces and inbred lines, and between zero and eight seminal roots are commonly reported (Zhu *et al*., 2006; Burton *et al*., 2013 *a*). Seminal root number is not correlated with seed size within the B73 x Mo17 and Ny821 x H99 populations of recombinant inbred lines, which might suggest that the two traits are under separate genetic control (Fig. S3). There is some evidence that inbreeding reduces seminal root number (Mangelsdorf and Goodsell, 1929), and inbred lines may have fewer seminal roots than hybrids (Salungyu *et al*., 2018). Northern flint corn, sweet corn, and popcorn generally have fewer seminal roots than dent corn lines (Wiggans, 1916; Siemens, 1929). This might be due to their relatedness (Revilla and Tracy, 1995), or it might be due to the unusual endosperm characteristics of those groups. Flint corn seeds have very vitreous endosperms (Gayral *et al*., 2016), and sweet corn seeds with the *sugary 1* mutation accumulate phytoglycogen, a branched form of starch, at the expense of amylose and amylopectin (Doehlert *et al*., 1993). Popcorn seeds are generally small, and flake formation is related to the ratio of hard to soft endosperm (Ziegler and Ashman, 1994). Our results suggest that seed carbohydrate content should not limit the optimal number of seminal roots for nutrient acquisition in dent corn due to its large kernel size, however.

While the importance of seminal roots for nutrient acquisition in maize probably decreases as the plant matures (Fig. 2), seminal root number may still have agronomic implications. Matching the rate and timing of nitrogen fertilizer applications to plant demand has the potential to reduce pollution and input costs (Ladha *et al*., 2005; Rutan and Steinke, 2018). For this approach to work best, nitrogen acquisition efficiency at all stages of development needs to be optimized. In addition, low soil temperatures at the time of spring planting may limit soil microbial processes and nutrient availability. Knowledge of how domestication has influenced root systems may also be useful for *de novo* domestication efforts.

The impact of seminal root number on seedling performance under drought was not simulated as drought responses are not yet fully implemented in *OpenSimRoot*. In the corn belt region of the United States, water is often available in the topsoil at the time of planting because of snowmelt and spring rains (Asbjornsen *et al*., 2008), so seedling water acquisition efficiency may be of less importance. In certain arid regions, such as the Central Rift Valley of Ethiopia, the Mexican highlands, and the southwestern United States, some indigenous growers have planted corn at depths of 10 cm and greater in order to ensure that sufficient moisture is available for germination (Collins, 1914; Eagles and Lothrup, 1994; Liben *et al*., 2015). Planting depth may influence the ideal seedling root system architecture in these systems, and it is possible that increased seminal root number has implications for seedling water acquisition in some arid environments.

While *OpenSimRoot* includes models of nitrogen mineralization, leaching, and depletion, it does not currently simulate soil temperature effects on mineralization or root respiration explicitly. In some parts of the world, maize may be planted into cold soils that have decreasing soil temperatures with depth (Gupta *et al*., 1982). This gradient might impact root maintenance costs or nutrient availability, among other things. Similarly, the model does not include soil hardness effects, which might be important because maize seminal roots are smaller in diameter than primary and nodal roots (Tai *et al*., 2015). Increased root diameter may be associated with improved root penetration in compacted soils (Clark *et al*., 2003), while thin roots may be more able to grow through small pores (Potocka and Szymanowska-Pulka, 2018).

Some of the comparisons made in this study would be challenging to validate empirically, such as the models that combine parts of the maize or teosinte phenotype with the phenotype of the other subspecies. Other components, such as the utility of seminal roots under low-nitrogen stress, could be validated if appropriate genetic resources are available. This study uses models that were parameterized to resemble one maize landrace and one teosinte accession, but modeling more of the diversity of both subspecies could also be informative. Teosinte is highly plastic, and teosinte plants that are more mature respond to suboptimal nitrogen availability by reducing their tiller number (Gaudin *et al*., 2011). This study considers only the first 25 days of growth because that is the period during which seminal roots make the largest contribution to nutrient acquisition, and tiller roots did not form during this time period in the plants used for parameterization. Experiments covering a longer period of growth would need to consider tiller number plasticity, however. The study of teosinte *ex situ* is also challenged by its photoperiod requirement (Minow *et al*., 2018) and sensitivity to inbreeding depression. Finally, seed nutrient content is somewhat variable, and maize grown in low-nitrogen environments forms kernels with decreased protein content (Mayer *et al*., 2012). Therefore, it is possible that the environment in which seed production occurs could influence some of the model parameters used here.

Increased seed size is often described as a component of domestication syndrome in grain crops. In maize, selection for seed size, coupled with increased seedling vigor, may have increased the nutrient demand of seedlings and facilitated the development of seminal roots. Understanding domestication effects on maize seedling root architecture may help to improve the stress tolerance of maize and other crops and inform domestication efforts in other species.

## Acknowledgements

We thank Michael Williams for technical assistance.

## Supplementary Information: OpenSimRoot Maize Landrace (PI 213706) Parametrization

OpenSimRoot uses a hierarchical input file which is summarized below. The hierarchy gives the parameters context. For example, the parameter ‘specific leaf area’ belongs to the shoot of a specific plant. In OpenSimRoot, parameters can be a single value, a value drawn from a distribution (random), or the result of an interpolation table. For constants we give the value, for distributions the distribution parameters and for the tables a list of space separated values e.g. x1 y1 x2 y2 …. xn yn.

**Table S1.** Means, standard errors, and sample size per treatment for selected model parameters and other measurements from the maize landrace (PI 213706) and the teosinte accession (Ames 21803). Significant differences were evaluated using a Mann-Whitney test. Sample sizes vary because plants sampled close to germination had not yet developed all root classes. Seminal root diameter is not reported for teosinte because it typically did not form seminal roots.

| **Trait** | **Maize Landrace** | **Teosinte** | **n** | ***p*** |
| --- | --- | --- | --- | --- |
| Seed N Available to Seedling (μmol) | 383.9 ± 46.0 | 20.1 ± 2.9 | 4 | < 0.01 |
| Seed P Available to Seedling (μmol) | 40.8 ± 4.0 | 3.0 ± 0.1 | 3 | 0.1 |
| Seminal Root Number | 3.90 ± 0.28 | 0.25 ± 0.14 | 20 | < 0.01 |
| First Whorl Nodal Root Number | 4.42 ± 0.23 | 2.73 ± 0.14 | 12 | < 0.01 |
| Second Whorl Nodal Root Number | 4.38 ± 0.26 | 3.25 ± 0.31 | 8 | 0.02 |
| Primary Root Basal Diameter (mm) | 1.09 ± 0.03 | 0.61 ± 0.05 | 20 | < 0.01 |
| Seminal Root Basal Diameter (mm) | 0.74 ± 0.06 |  | 20 |  |
| First Whorl Nodal Root Basal Diameter (mm) | 0.84 ± 0.03 | 0.69 ± 0.02 | 12 | < 0.01 |
| Second Whorl Nodal Root Basal Diameter (mm) | 1.26 ± 0.05 | 1.00 ± 0.03 | 8 | < 0.01 |
| First Whorl Nodal Root Angle (°) | 132 ± 10 | 127 ± 14 | 3 | 1 |
| Second Whorl Nodal Root Angle (°) | 133 ± 3 | 120 ± 4 | 3 | 0.2 |
| Biomass at 25 days after planting (g) | 5.85 ± 0.36 | 1.25 ± 0.17 | 4 | 0.03 |
| Leaf area at 25 days after planting (cm<sup>2</sup>) | 799 ± 60 | 226 ± 21 | 4 | 0.03 |

**Fig S1.**
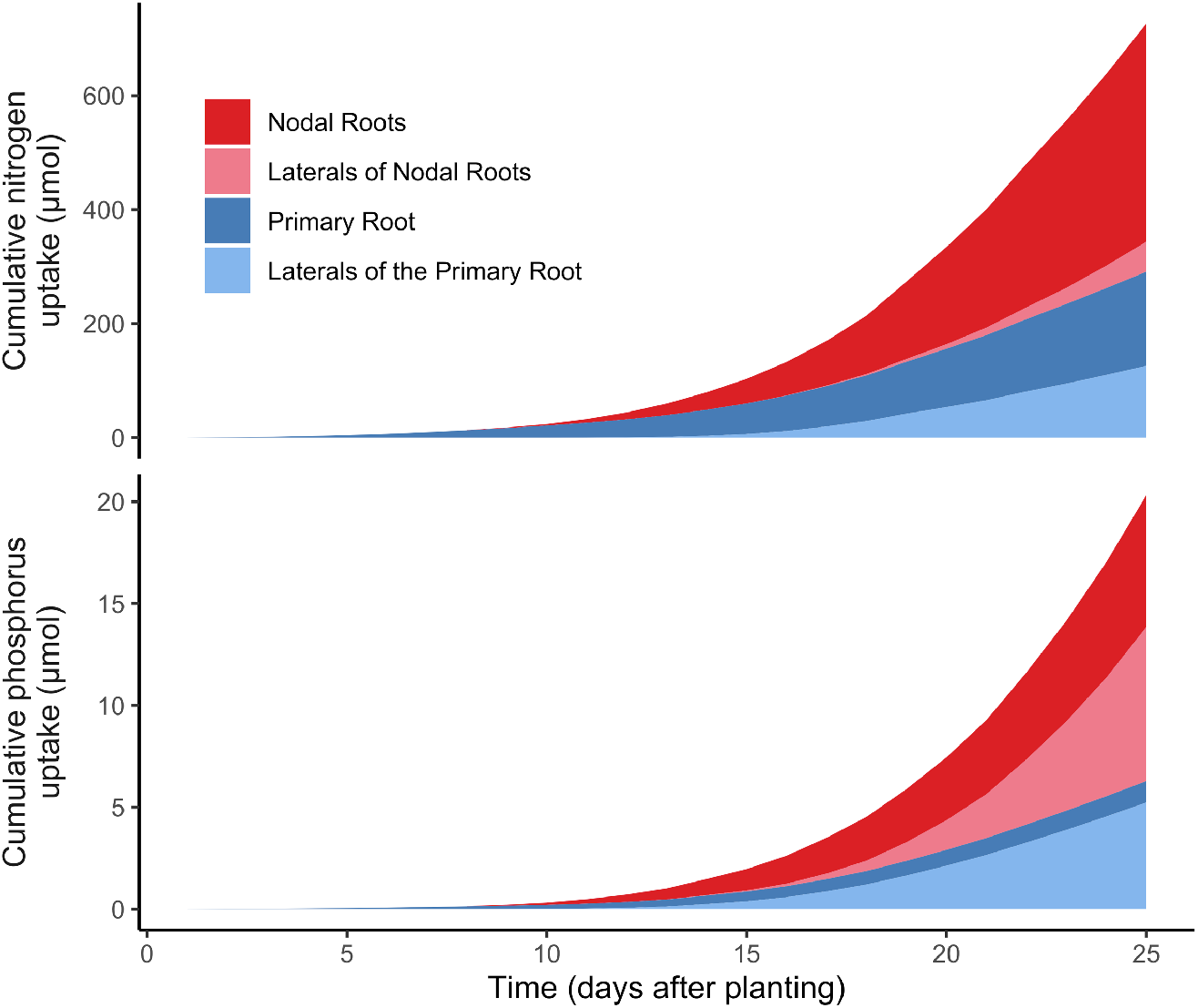
The contribution of the primary and nodal roots to nutrient acquisition in teosinte grown in field conditions with 50 kg ha^-1^ available nitrate (top) and 2 kg ha^-1^ available phosphorus (bottom). The values presented are an average of six model replications that include stochasticity. Nodal roots emerged seven days after planting.

**Fig S2.**
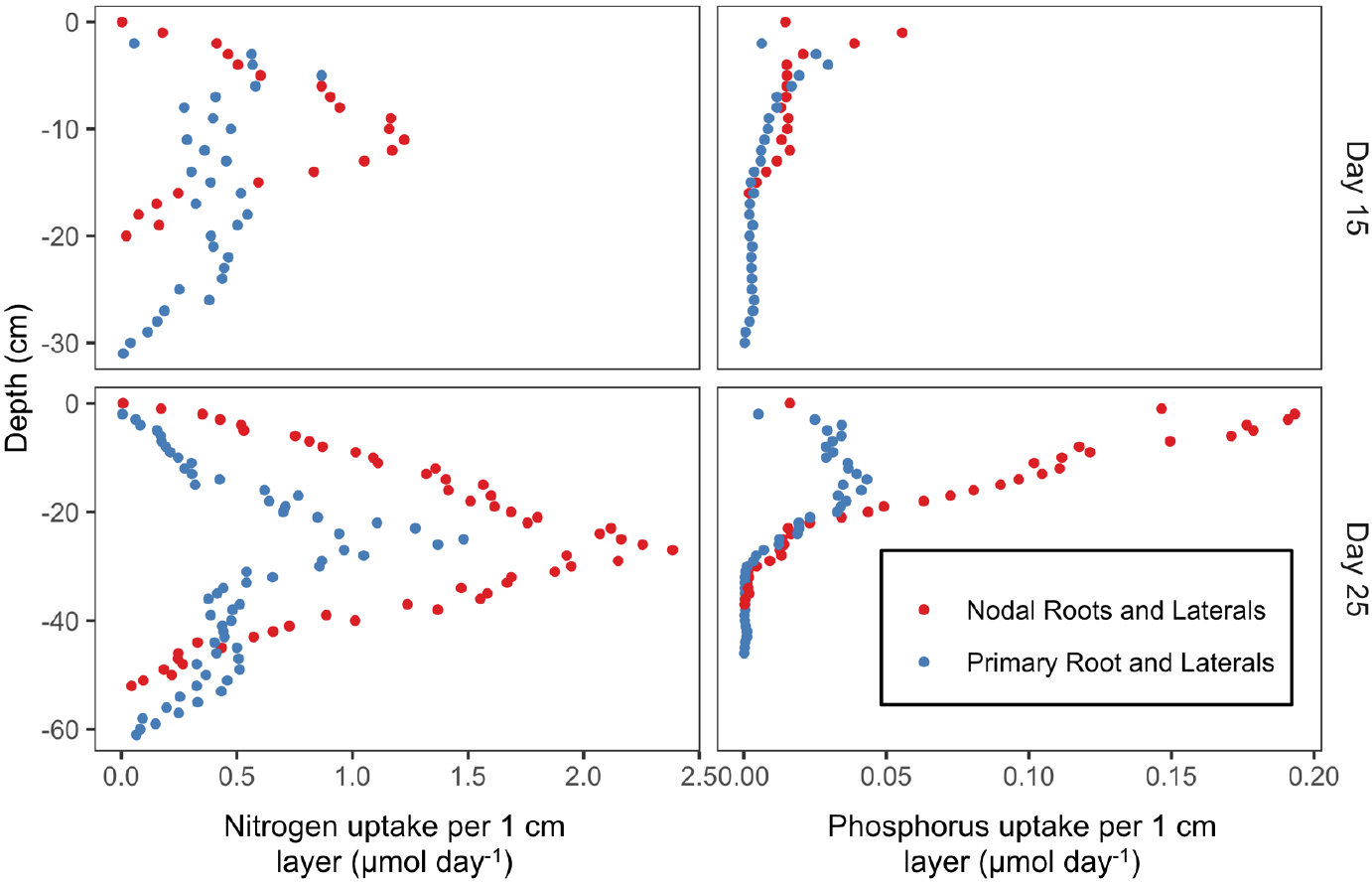
The acquisition of nitrogen and phosphorus by teosinte at 15 and 25 days after planting. Soils with 50 kg ha^-1^ available nitrate (left) and 2 kg ha^-1^ available phosphorus (right) were used. Points represent an average of six replications.

**Fig S3.**
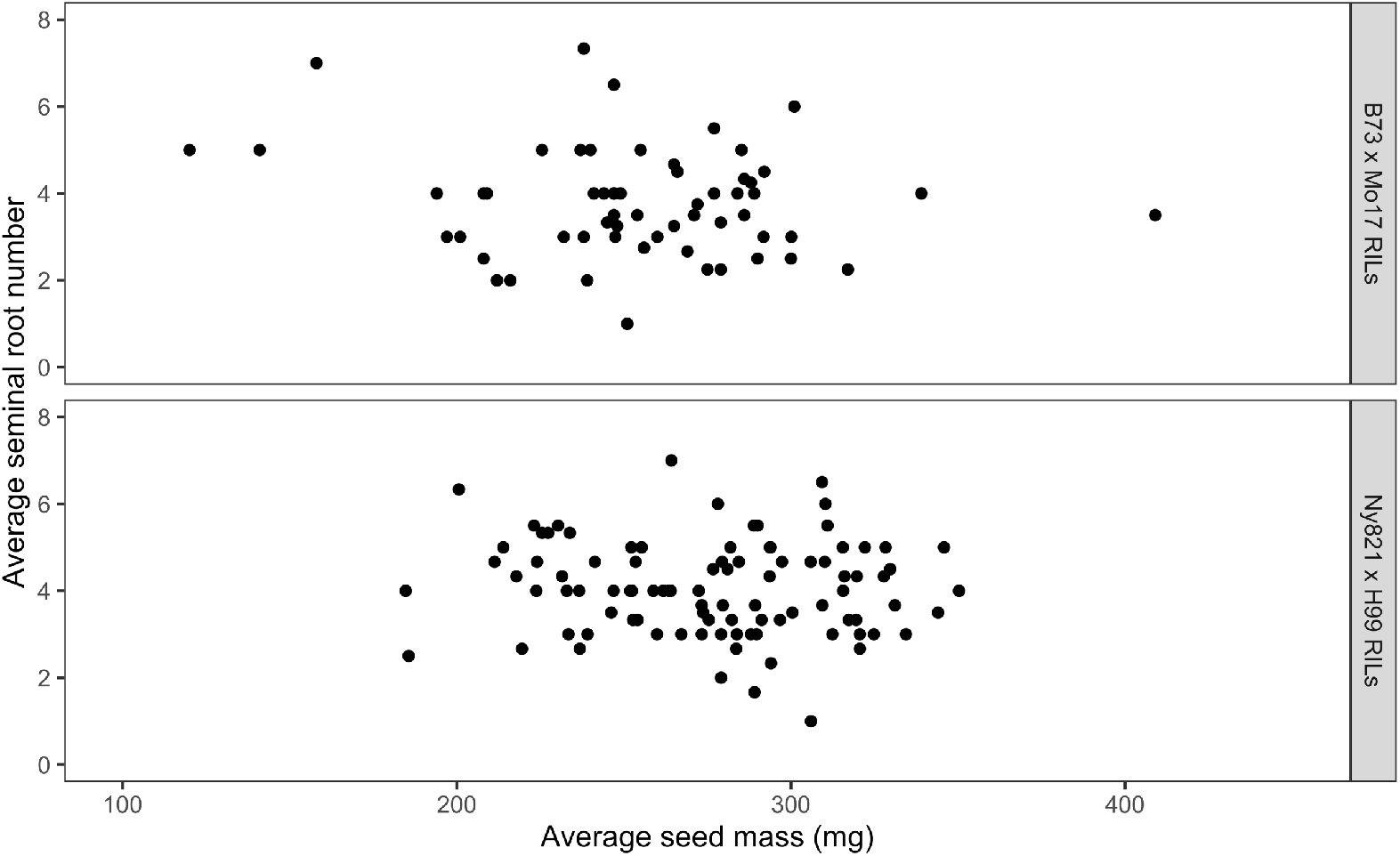
Seminal root number does not appear to be related to seed mass in dent corn recombinant inbred lines resulting from B73 x Mo17 (Spearman’s rank correlation = −0.089) and Ny821 x H99 (Spearman’s rank correlation = −0.086).

**Fig S4.**
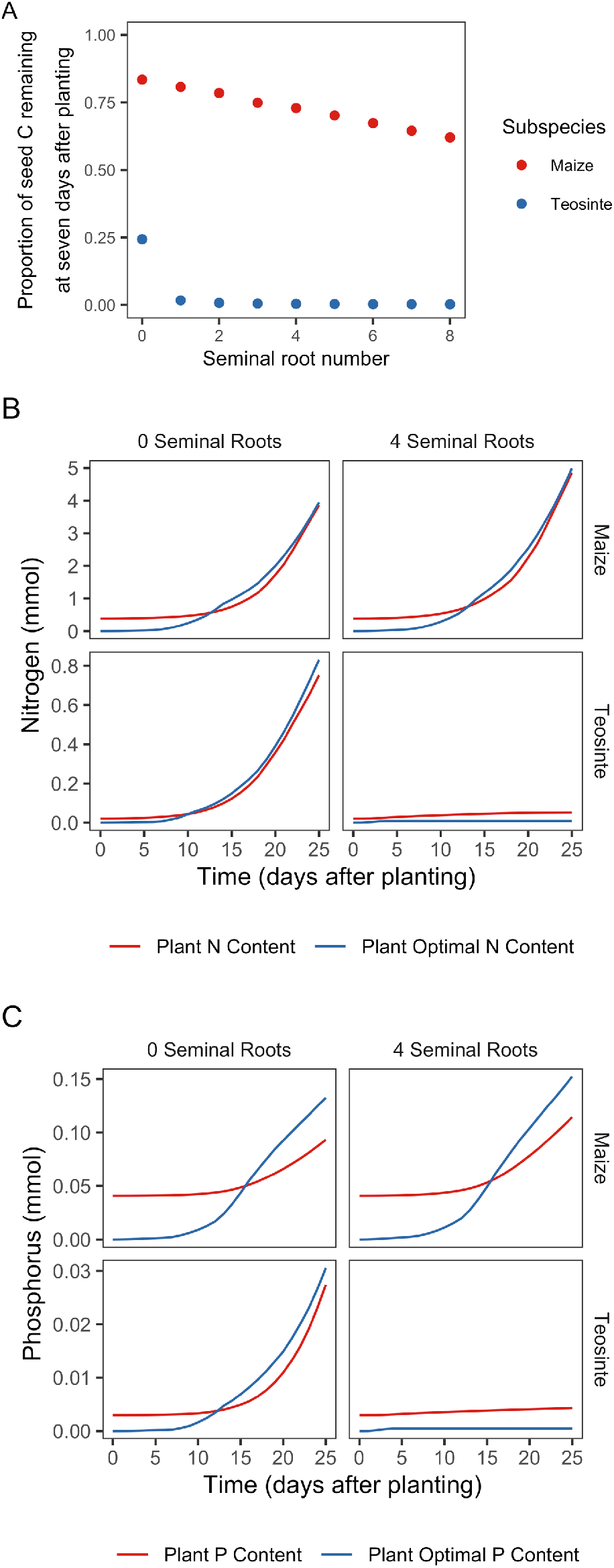
The temporal nature of seedling carbon and nutrient stress. (A) The proportion of the starting seed carbon reserves remaining at seven days after planting for maize and teosinte. The low-N environment with 50 kg ha^-1^ available nitrate was used. (B) The timing of low-nitrogen stress onset in maize and teosinte grown in an environment with 50 kg ha^-1^ available nitrate. (C) The timing of low-phosphorus stress onset in an environment with 2 kg ha^-1^ available phosphorus.

