## Supplementary Information for "Increased seed carbohydrate reserves associated with domestication influence the optimal seminal root number of *Zea mays*"

### *OpenSimRoot* Maize Landrace (PI 213706) Parametrization

*OpenSimRoot* uses a hierarchical input file which is summarized below. The hierarchy gives the parameters context. For example, the parameter 'specific leaf area' belongs to the shoot of a specific plant. In *OpenSimRoot*, parameters can be a single value, a value drawn from a distribution (random), or the result of an interpolation table. For constants we give the value, for distributions the distribution parameters and for the tables a list of space separated values e.g. x1 y1 x2 y2 .... xn yn.

1. 'data point template'(noUnit)
    1. 'aerenchyma formation'
    2. 'branching frequency multiplier'(noUnit)
    3. 'combined root class ID'(noUnit)
    4. 'parent root class ID'(noUnit)
    5. 'root circumference'(cm)
    6. 'root class ID'(noUnit)
    7. 'root diameter'(cm) initial value = 0 0 0
    8. 'root hair density'
    9. 'root hair diameter'
    10. 'root hair length'
    11. 'root hair surface area'(cm2/cm)
    12. 'root length to base'(cm)
    13. 'root potential secondary growth'(cm) initial value = 0 0 0
    14. 'root segment age'(day)
    15. 'root segment carbon cost of exudates'(g) initial value = 0 0 0
    16. 'root segment dry weight'(g)
    17. 'root segment length'(cm)
    18. 'root segment length duration'(cm.day) initial value = 0 0 0
    19. 'root segment respiration'(g/day)
    20. 'root segment secondary potential carbon sink for growth'(g) initial value = 0 0 0
    21. 'root segment surface area'(cm2)
    22. 'root segment volume'(cm3)
    23. 'spatial root density'(cm/cm3)
2. 'environment'(noUnit)
    1. 'atmosphere'(noUnit)
        1. 'PAR/ RDD' = 1 (100%)
        2. 'actual durationof sunshine' x,y pairs :{ 0 0 60 0 }
        3. 'albedo crop' = 0.23 (noUnit)
        4. 'albedo soil' = 0.17 (noUnit)
        5. 'altitude' = 91 (m)
        6. 'average daily temperature' x,y pairs :{ 0 16.9 100 16.9 }
        7. 'evaporation' x,y pairs :{ 0 0 1 0.05 2 0.1 3 0.1 4 0.05 5 0.05 6 0.1 7 0.05 8 0.05 9 0.1 10 0.1 11 0.05 12 0.1 13 0.1 14 0.05 15 0.04 16 0.03 17 0.02 18 0.09 19 0.09 20 0.04 21 0.09 22 0.09 23 0.04 24 0.03 25 0.02 26 0.02 27 0.08 28 0.03 29 0.08 30 0.03 31 0.08 32 0.07 33 0.07 34 0.07 35 0.03 36 0.02 37 0.01 38 0 39 0 40 0 41 0 42 0.06 }
        8. 'irradiation' = 4000 (umol/cm2/day)
        9. 'latitude' = 50.8 (noUnit)
        10. 'net radiation' = 50 (W/m2)
        11. 'net radiation soil' = 0 (W/m2)
        12. 'precipitation' x,y pairs :{ 0 0 1 0 2 1 3 0.29 4 0 5 0 6 0.61 7 0 8 0 9 0.25 10 0.03 11 0 12 0.64 13 0.33 14 0 15 0 16 0 17 0 18 1.8 19 0.2 20 0 21 2.84 22 0.38 23 0 24 0 25 0 26 0 27 0.18 28 0 29 0.46 30 0 31 1.35 32 0.13 33 0.23 34 0.25 35 0 36 0 37 0 38 0 39 0 40 0 41 0 42 1.42 }
        13. 'relative humidity' = 60 (m/s)
        14. 'start day' = 22 (noUnit)
        15. 'start month' = 6 (noUnit)
        16. 'wind speed' = 2 (m/s)
    2. 'dimensions'(noUnit)
        1. 'max corner' = 13 0 30 (cm)
        2. 'min corner' = -13 -150 -30 (cm)
        3. 'resolution' = 1 1 1 (cm)
    3. 'soil'(noUnit)
        1. 'bulk density' x,y pairs :{ -200 1.51 -65 1.51 -47 1.4 -30 1.42 -16 1.29 -5 1.24 0 1.24 }
        2. 'nitrate'(noUnit)
            1. 'adsorption coefficient' = 0 (umol/cm)
            2. 'buffer power' x,y pairs :{ -1000 0.4 1000 0.4 }

            3. 'concentration' x,y pairs :{ -1000 1.59 -55 1.59 -45 1.67 -35 2.17 -25 3.15 -15 4.02 -5 2.36 0 2.8 0.01 0 100 0 }
            4. 'diffusion coefficient' x,y pairs :{ -1000 0.07 -0 0.07 1e-05 1e-08 1000 1e-08 }
            5. 'increase time step' = 1 (noUnit)
            6. 'longitudinal dispersivity' = 1 (cm)
            7. 'r1-r0' = 4 (cm)
            8. 'saturated diffusion coefficient' = 1.6416 (cm2/day)
            9. 'transverse dispersivity' = 0.5 (cm)
        3. 'organic'(noUnit)
            1. 'CNRatio microbes' = 10 (g/g)
            2. 'CNratio' x,y pairs :{ -10000 13 0 13 }
            3. 'assimilation efficiency microbes' = 1 (noUnit)
            4. 'carbon content' x,y pairs :{ -200 0.005 -40 0.005 -30 0.01 -10 0.02 0 0.02 }
            5. 'initial relative mineralisation rate' x,y pairs :{ -1000 0 -25 0 -10 0.037 0 0.037 }
            6. 'speed of aging' = 0.46 (noUnit)
            7. 'time offset' = 30 (day)
        4. 'phosphorus'(noUnit)
            1. 'adsorption coefficient' = 1333.3 (umol/cm)
            2. 'buffer power' x,y pairs :{ -1000 400 1000 400 }
            3. 'concentration' x,y pairs :{ -1000 0.00024 -30 0.00025 -29 0.00175 0 0.00175 0.0001 0 1000 0 }
            4. 'diffusion coefficient' x,y pairs :{ -1000 0.00019872 1000 0.00019872 }
            5. 'increase time step' = 1.1 (noUnit)
            6. 'longitudinal dispersivity' = 0 (cm)
            7. 'r1-r0' = 0.3 (cm)
            8. 'saturated diffusion coefficient' = 0.00495 (cm2/day)
            9. 'transverse dispersivity' = 0 (cm)
        5. 'potassium'(noUnit)
            1. 'adsorption coefficient' = 33.3 (umol/cm)
            2. 'buffer power' x,y pairs :{ -1000 10 1000 10 }
            3. 'concentration' x,y pairs :{ -1000 0.05 -30 0.05 -29 0.15 0 0.15 1e-05 0 1000 0 }
            4. 'diffusion coefficient' x,y pairs :{ -1000 0.067 1000 0.067 }
            5. 'increase time step' = 1.01 (noUnit)
            6. 'longitudinal dispersivity' = 1 (cm)
            7. 'r1-r0' = 1.5 (cm)
            8. 'saturated diffusion coefficient' = 1.56 (cm2/day)
            9. 'transverse dispersivity' = 0.5 (cm)
        6. 'water'(noUnit)
            1. 'initial hydraulic head' x,y pairs :{ -200 -10 -150 -52 -60 -143 0 -215 }
            2. 'residual water content' x,y pairs :{ -300 0.067 0 0.067 }
            3. 'saturated conductivity' x,y pairs :{ -300 1 -150 1 -140 10.8 0 10.8 }
            4. 'saturated water content' x,y pairs :{ -300 0.39 -65 0.39 -35 0.39 -25 0.43 -15 0.45 0 0.46 }
            5. 'van genuchten:alpha' x,y pairs :{ -300 0.02 0 0.02 }
            6. 'van genuchten:n' x,y pairs :{ -300 1.41 0 1.41 }
            7. 'volumetric water content in barber cushman' = 0.3 (cm3/cm3)
3. 'hypocotyl template'(noUnit)
    1. 'branches'(noUnit)
    2. 'data points'(noUnit)
    3. 'growthpoint'(cm) initial position = 0 0 0 0 0 0 0
        1. 'branching frequency multiplier'(noUnit)
        2. 'root circumference'(cm)
        3. 'root diameter'(cm)
        4. 'root longitudinal growth'(cm) initial value = 0 0 0
        5. 'root potential longitudinal growth'(cm) initial value = 0 0 0
        6. 'root potential secondary growth'(cm)
        7. 'root segment age' = 0 (day)
        8. 'root segment carbon cost of exudates' = 0 (g)
        9. 'root segment dry weight' = 0 (g)
        10. 'root segment length' = 0 (cm)
        11. 'root segment length duration' = 0 (cm.day)
        12. 'root segment potential carbon sink for growth'(g) initial value = 0 0 0
        13. 'root segment respiration' = 0 (g)
        14. 'root segment secondary potential carbon sink for growth' = 0 (g)
        15. 'root segment surface area' = 0 (cm2)

16. 'root segment volume' = 0 (cm3)
4. 'root carbon cost of exudates'(g) initial value = 0 0 0
5. 'root dry weight'(g)
6. 'root length'(cm)
7. 'root respiration'(g) initial value = 0 0 0
8. 'root secondary potential carbon sink for growth'(g) initial value = 0 0 0
9. 'root surface area'(cm2)
10. 'root system carbon cost of exudates'(g) initial value = 0 0 0
11. 'root system dry weight'(g)
12. 'root system length'(cm)
13. 'root system longitudinal growth'(cm) initial value = 0 0 0
14. 'root system potential carbon sink for growth'(g) initial value = 0 0 0
15. 'root system potential carbon sink for growth;major axis'(g) initial value = 0 0 0
    1. 'included root classes' = hypocotyl, primaryRoot, seminal, nodalroots, nodalroots1, nodalroots2, nodalroots3, nodalroots4, nodalroots5, braceroots, braceroots1, braceroots2, braceroots3, basalWhorl1, basalWhorl2, basalWhorl3, basalWhorl4 (noUnit)
16. 'root system respiration'(g) initial value = 0 0 0
17. 'root system secondary potential carbon sink for growth'(g) initial value = 0 0 0
18. 'root system surface area'(cm2)
19. 'root system volume'(cm3)
20. 'root volume'(cm3)

4. 'plant template'(noUnit)
    1. 'carbon allocation to roots'(g) initial value = 0 0 0
    2. 'carbon allocation to shoot'(g) initial value = 0 0 0
    3. 'carbon available for growth'(g) initial value = 0 0 0
    4. 'carbon reserves'(g) initial value = 0 0 0
    5. 'carbon to dry weight ratio'(100%)
    6. 'plant carbon balance'(g)
    7. 'plant carbon income'(g) initial value = 0 0 0
    8. 'plant dry weight'(g)
    9. 'plant potential carbon sink for growth'(g) initial value = 0 0 0
    10. 'plant respiration'(g) initial value = 0 0 0
    11. 'relative carbon allocation to roots'(100%)
    12. 'relative carbon allocation to shoot'(100%)
    13. 'reserves'(g) initial value = 0 0 0
    14. 'root carbon cost of biological nitrogen fixation'(g) initial value = 0 0 0
    15. 'root carbon cost of exudates'(g) initial value = 0 0 0
    16. 'root carbon cost of nutrient uptake'(g) initial value = 0 0 0
    17. 'root carbon costs'(g) initial value = 0 0 0
        1. 'paths' = rootCarbonCostOfExudates;rootCarbonCostOfNutrientUptake;rootCarbonCostOfBiologicalNitrogenFixation (noUnit)
    18. 'root dry weight'(g)
    19. 'root growth scaling factor'(100%) initial value = 0 0 0
    20. 'root growth scaling factor;major axis'(100%) initial value = 0 0 0
    21. 'root length'(cm)
    22. 'root longitudinal growth'(cm) initial value = 0 0 0
    23. 'root potential carbon sink for growth'(g) initial value = 0 0 0
    24. 'root potential carbon sink for growth;major axis'(g) initial value = 0 0 0
    25. 'root respiration'(g) initial value = 0 0 0
    26. 'root secondary potential carbon sink for growth'(g) initial value = 0 0 0
    27. 'root surface area'(cm2)
    28. 'root volume'(cm3)
    29. 'secondary root growth scaling factor'(100%) initial value = 0 0 0
    30. 'seed carbohydrate content'(100%)
    31. 'seed carbohydrate to CFactor'(100%)
    32. 'shoot dry weight'(g)
    33. 'shoot potential carbon sink for growth'(g) initial value = 0 0 0
    34. 'shoot respiration'(g) initial value = 0 0 0
    35. 'stress factor'(noUnit)
    36. 'stress factor:impact on:leaf area expantion rate'(noUnit)
    37. 'stress factor:impact on:leaf respiration'(noUnit)
    38. 'stress factor:impact on:photosynthesis'(noUnit)

    39. 'stress factor:impact on:root potential longitudinal growth'(noUnit)
    40. 'stress factor:impact on:root segment carbon cost of exudates'(noUnit)
    41. 'stress factor:impact on:root segment respiration'(noUnit)
    42. 'stress factor:impact on:root segment secondary growth'(noUnit)
    43. 'stress factor:impact on:stem respiration'(noUnit)
5. 'plants'(noUnit)
    1. 'maize'(noUnit)
        1. 'carbon allocation to roots'(g) initial value = 0 0 0.000841285
        2. 'carbon allocation to shoot'(g) initial value = 0 0 0
        3. 'carbon available for growth'(g) initial value = 0 0 0.000841285
        4. 'carbon reserves'(g) initial value = 0 0 0
        5. 'carbon to dry weight ratio'(100%)
        6. 'plant carbon balance'(g)
        7. 'plant carbon income'(g) initial value = 0 0 0.000841285
        8. 'plant dry weight'(g)
        9. 'plant position' = 0 -2 0 (cm)
            1. 'hypocotyl' = 0 0 0 (cm)
                1. 'branches'(noUnit)
                    1. 'braceroots'(noUnit)
                    2. 'braceroots2'(noUnit)
                    3. 'nodalroots'(noUnit)
                    4. 'nodalroots2'(noUnit)
                    5. 'nodalroots3'(noUnit)
                    6. 'nodalroots4'(noUnit)
                2. 'data points'(noUnit)
                    1. 'data point00000' = 0 0 0 (cm)
                        1. 'aerenchyma formation'
                        2. 'branching frequency multiplier'(noUnit)
                        3. 'combined root class ID'(noUnit)
                        4. 'parent root class ID'(noUnit)
                        5. 'root circumference'(cm)
                        6. 'root class ID'(noUnit)
                        7. 'root diameter'(cm) initial value = 0 0.2 0
                        8. 'root hair density'
                        9. 'root hair diameter'
                        10. 'root hair length'
                        11. 'root hair surface area'(cm2/cm)
                        12. 'root length to base'(cm)
                        13. 'root potential secondary growth'(cm) initial value = 0 0.2 0
                        14. 'root segment age'(day)
                        15. 'root segment carbon cost of exudates'(g) initial value = 0 0 0
                        16. 'root segment dry weight'(g)
                        17. 'root segment length'(cm)
                        18. 'root segment length duration'(cm.day) initial value = 0 0 0
                        19. 'root segment respiration'(g/day)
                        20. 'root segment secondary potential carbon sink for growth'(g) initial value = 0 0 0
                        21. 'root segment surface area'(cm2)
                        22. 'root segment volume'(cm3)
                        23. 'spatial root density'(cm/cm3)
                3. 'growth rate multiplier' = 1 (noUnit)
                4. 'growthpoint'(cm) initial position = 0 0 0 0 0 0 0
                    1. 'branching frequency multiplier'(noUnit)
                    2. 'root circumference'(cm)
                    3. 'root diameter'(cm)
                    4. 'root longitudinal growth'(cm) initial value = 0 0 0
                    5. 'root potential longitudinal growth'(cm) initial value = 0 0 0
                    6. 'root potential secondary growth'(cm)
                    7. 'root segment age' = 0 (day)
                    8. 'root segment carbon cost of exudates' = 0 (g)
                    9. 'root segment dry weight' = 0 (g)
                    10. 'root segment length' = 0 (cm)
                    11. 'root segment length duration' = 0 (cm.day)
                    12. 'root segment potential carbon sink for growth'(g) initial value = 0 0 0

            13. 'root segment respiration' = 0 (g)
            14. 'root segment secondary potential carbon sink for growth' = 0 (g)
            15. 'root segment surface area' = 0 (cm2)
            16. 'root segment volume' = 0 (cm3)
        5. 'root carbon cost of exudates'(g) initial value = 0 0 0
        6. 'root dry weight'(g)
        7. 'root length'(cm)
        8. 'root respiration'(g) initial value = 0 0 0
        9. 'root secondary potential carbon sink for growth'(g) initial value = 0 0 0
        10. 'root surface area'(cm2)
        11. 'root system carbon cost of exudates'(g) initial value = 0 0 0
        12. 'root system dry weight'(g)
        13. 'root system length'(cm)
        14. 'root system longitudinal growth'(cm) initial value = 0 0 0
        15. 'root system potential carbon sink for growth'(g) initial value = 0 0 0
        16. 'root system potential carbon sink for growth;major axis'(g) initial value = 0 0 0
            1. 'included root classes' = hypocotyl, primaryRoot, seminal, nodalroots, nodalroots1, nodalroots2, nodalroots3, nodalroots4, nodalroots5, braceroots, braceroots1, braceroots2, braceroots3, basalWhorl1, basalWhorl2, basalWhorl3, basalWhorl4 (noUnit)
        17. 'root system respiration'(g) initial value = 0 0 0
        18. 'root system secondary potential carbon sink for growth'(g) initial value = 0 0 0
        19. 'root system surface area'(cm2)
        20. 'root system volume'(cm3)
        21. 'root type' = hypocotyl (noUnit)
        22. 'root volume'(cm3)
2. 'primary root' = 0 0 0 (cm)
    1. 'branches'(noUnit)
        1. 'lateral'(noUnit)
        2. 'seminal'(noUnit)
    2. 'data points'(noUnit)
        1. 'data point00000' = 0 0 0 (cm)
            1. 'aerenchyma formation'
            2. 'branching frequency multiplier'(noUnit)
            3. 'combined root class ID'(noUnit)
            4. 'parent root class ID'(noUnit)
            5. 'root circumference'(cm)
            6. 'root class ID'(noUnit)
            7. 'root diameter'(cm) initial value = 0 0.088 0
            8. 'root hair density'
            9. 'root hair diameter'
            10. 'root hair length'
            11. 'root hair surface area'(cm2/cm)
            12. 'root length to base'(cm)
            13. 'root potential secondary growth'(cm) initial value = 0 0.088 0
            14. 'root segment age'(day)
            15. 'root segment carbon cost of exudates'(g) initial value = 0 0 0
            16. 'root segment dry weight'(g)
            17. 'root segment length'(cm)
            18. 'root segment length duration'(cm.day) initial value = 0 0 0
            19. 'root segment respiration'(g/day)
            20. 'root segment secondary potential carbon sink for growth'(g) initial value = 0 0 0
            21. 'root segment surface area'(cm2)
            22. 'root segment volume'(cm3)
            23. 'spatial root density'(cm/cm3)
    3. 'growth rate multiplier' = 1 (noUnit)
    4. 'growthpoint'(cm) initial position = 0 0 0 0 0 0 0
        1. 'branching frequency multiplier'(noUnit)
        2. 'gravitropism'
            1. 'multiplier'(noUnit)
        3. 'root circumference'(cm)
        4. 'root diameter'(cm)
        5. 'root longitudinal growth'(cm) initial value = 0 0 3.27
        6. 'root potential longitudinal growth'(cm) initial value = 0 0 3.27

                1. 'rate multiplier'(noUnit)
            7. 'root potential secondary growth'(cm)
            8. 'root segment age' = 0 (day)
            9. 'root segment carbon cost of exudates' = 0 (g)
            10. 'root segment dry weight' = 0 (g)
            11. 'root segment length' = 0 (cm)
            12. 'root segment length duration' = 0 (cm.day)
            13. 'root segment potential carbon sink for growth'(g) initial value = 0 0 0.000841285
            14. 'root segment respiration' = 0 (g)
            15. 'root segment secondary potential carbon sink for growth' = 0 (g)
            16. 'root segment specific weight' = 0 (g/cm3)
            17. 'root segment surface area' = 0 (cm2)
            18. 'root segment volume' = 0 (cm3)
        5. 'root carbon cost of exudates'(g) initial value = 0 0 0
        6. 'root dry weight'(g)
        7. 'root length'(cm)
        8. 'root respiration'(g) initial value = 0 0 0
        9. 'root secondary potential carbon sink for growth'(g) initial value = 0 0 0
        10. 'root surface area'(cm2)
        11. 'root system carbon cost of exudates'(g) initial value = 0 0 0
        12. 'root system dry weight'(g)
        13. 'root system length'(cm)
        14. 'root system longitudinal growth'(cm) initial value = 0 0 3.27
        15. 'root system potential carbon sink for growth'(g) initial value = 0 0 0.000841285
        16. 'root system potential carbon sink for growth;major axis'(g) initial value = 0 0 0.000841285
            1. 'included root classes' = hypocotyl, primaryRoot, seminal, nodalroots, nodalroots1, nodalroots2, nodalroots3, nodalroots4, nodalroots5, braceroots, braceroots1, braceroots2, braceroots3, basalWhorl1, basalWhorl2, basalWhorl3, basalWhorl4 (noUnit)
        17. 'root system respiration'(g) initial value = 0 0 0
        18. 'root system secondary potential carbon sink for growth'(g) initial value = 0 0 0
        19. 'root system surface area'(cm2)
        20. 'root system volume'(cm3)
        21. 'root type' = primaryRoot (noUnit)
        22. 'root volume'(cm3)
    3. 'shoot' = 0 0 0 (cm)
        1. 'area per plant'
        2. 'carbon allocation to leafs'(g) initial value = 0 0 0
        3. 'carbon allocation to stems'(g) initial value = 0 0 0
        4. 'extinction coefficient'
        5. 'leaf area'(cm2) initial value = 0 0 0
        6. 'leaf area index'(cm2/cm2)
        7. 'leaf area reduction coefficient'(cm2/cm2)
        8. 'leaf dry weight'(g) initial value = 0 0 0
        9. 'leaf potential carbon sink for growth'(g) initial value = 0 0 0
        10. 'leaf respiration'(g) initial value = 0 0 0
        11. 'light interception'(umol/cm2/day)
        12. 'photosynthesis'(g) initial value = 0 0 0
        13. 'potential leaf area'(cm2) initial value = 0 0 0
        14. 'relative carbon allocation to leafs'(100%)
        15. 'relative carbon allocation to stems'(100%)
        16. 'stem dry weight'(g) initial value = 0 0 0
        17. 'stem potential carbon sink for growth'(g) initial value = 0 0 0
        18. 'stem respiration'(g) initial value = 0 0 0
        19. 'stress adjusted potential leaf area'(cm2) initial value = 0 0 0
10. 'plant potential carbon sink for growth'(g) initial value = 0 0 0.000841285
11. 'plant respiration'(g) initial value = 0 0 0
12. 'plant type' = PI213706 (noUnit)
13. 'planting time' = 0 (day)
14. 'relative carbon allocation to roots'(100%)
15. 'relative carbon allocation to shoot'(100%)
16. 'reserves'(g) initial value = 0 0.35 -0.00266271
17. 'root carbon cost of biological nitrogen fixation'(g) initial value = 0 0 0
18. 'root carbon cost of exudates'(g) initial value = 0 0 0

                    19. 'root carbon cost of nutrient uptake'(g) initial value = 0 0 0
                    20. 'root carbon costs'(g) initial value = 0 0 0
                        1. 'paths' = rootCarbonCostOfExudates;rootCarbonCostOfNutrientUptake;rootCarbonCostOfBiologicalNitrogenFixation (noUnit)
                    21. 'root dry weight'(g)
                    22. 'root growth scaling factor'(100%) initial value = 0 1 0
                    23. 'root growth scaling factor;major axis'(100%) initial value = 0 1 0
                    24. 'root length'(cm)
                    25. 'root longitudinal growth'(cm) initial value = 0 0 3.27
                    26. 'root potential carbon sink for growth'(g) initial value = 0 0 0.000841285
                    27. 'root potential carbon sink for growth;major axis'(g) initial value = 0 0 0.000841285
                    28. 'root respiration'(g) initial value = 0 0 0
                    29. 'root secondary potential carbon sink for growth'(g) initial value = 0 0 0
                    30. 'root surface area'(cm2)
                    31. 'root volume'(cm3)
                    32. 'secondary root growth scaling factor'(100%) initial value = 0 1 0
                    33. 'seed carbohydrate content'(100%)
                    34. 'seed carbohydrate to CFactor'(100%)
                    35. 'shoot dry weight'(g)
                    36. 'shoot potential carbon sink for growth'(g) initial value = 0 0 0
                    37. 'shoot respiration'(g) initial value = 0 0 0
                    38. 'stress factor'(noUnit)
                    39. 'stress factor:impact on:leaf area expantion rate'(noUnit)
                    40. 'stress factor:impact on:leaf respiration'(noUnit)
                    41. 'stress factor:impact on:photosynthesis'(noUnit)
                    42. 'stress factor:impact on:root potential longitudinal growth'(noUnit)
                    43. 'stress factor:impact on:root segment carbon cost of exudates'(noUnit)
                    44. 'stress factor:impact on:root segment respiration'(noUnit)
                    45. 'stress factor:impact on:root segment secondary growth'(noUnit)
                    46. 'stress factor:impact on:stem respiration'(noUnit)
6. 'root type parameters'(noUnit)
    1. 'PI to 13706'(noUnit)
        1. 'braceroots'(noUnit)
            1. 'aerenchyma formation' x,y pairs :{ 0 0 3 0 5 0.1 10 0.25 20 0.393 1000 0.393 }
            2. 'bottom boundary' = 0 (noUnit)
            3. 'bounce of the side' = 0 (noUnit)
            4. 'branch list'(noUnit)
                1. 'lateral of crown roots'(noUnit)
                    1. 'allow branches to form above ground' = 0 (noUnit)
                    2. 'branching frequency'(cm)=f{'uniform distribution'} minimum=0.100000 maximum=0.300000
                    3. 'branching spatial offset' = 12 (cm)
                    4. 'length root tip' = 10.93 (cm)
                    5. 'number of branches/whorl' = 1 (#)
            5. 'branching angle' = 140 (degrees)
            6. 'cannotgrowup' = 1 (noUnit)
            7. 'copy defaults from' = ../defaultsMajorAxis (noUnit)
            8. 'density' = 0.094 (g/cm3)
            9. 'diameter' x,y pairs :{ 0 0.4 8 0.4 15 0.15 24 0.1 100 0.1 }
            10. 'gravitropism.v2'(cm)=f{'uniform distribution'} minimum=-0.010000 maximum=-0.005000
            11. 'growth rate' x,y pairs :{ 0 0.01 5 1 10 4.5 17 4.5 22 0 1000 0 }
            12. 'length multiplier to diameter multiplier' x,y pairs :{ 0 0.25 1 1 2 1.5 3 1.8 4 2 100 2 1000 2 }
            13. 'length root tip without xylem vessels' = 2 (cm)
            14. 'local resource responses'(noUnit)
                1. 'impact on:branching frequency'(noUnit)
                    1. 'aggregation function' = maxRelativeDeviationFromOne (noUnit)
                    2. 'impact by:nitrate' x,y pairs :{ 0 1 2000 1 }
                    3. 'impact by:phosphorus' x,y pairs :{ 0 0.2 0.015 1 1000 1 }
                    4. 'impact by:potassium' x,y pairs :{ 0 1 1000 1 }
                2. 'impact on:gravitropism'(noUnit)
                    1. 'aggregation function' = maxRelativeDeviationFromOne (noUnit)
                    2. 'impact by:nitrate' x,y pairs :{ 0 1.5 100 1 2000 1 }
                    3. 'impact by:phosphorus' x,y pairs :{ 0 0.5 0.015 2 1000 0.5 }

            4. 'impact by:potassium' x,y pairs :{ 0 1 1000 1 }
        3. 'impact on:root potential longitudinal growth'(noUnit)
            1. 'aggregation function' = maxRelativeDeviationFromOne (noUnit)
            2. 'impact by:nitrate' x,y pairs :{ 0 1 2000 1 }
            3. 'impact by:phosphorus' x,y pairs :{ 0 0.2 0.015 1 1000 1 }
            4. 'impact by:potassium' x,y pairs :{ 0 1 1000 1 }
15. 'longitudinal growth rate multiplier'(cm)=f{'uniform distribution'} minimum=0.700000 maximum=1.000000
16. 'nitrate'(noUnit)
    1. 'Cmin' = 0.001 (umol/ml)
    2. 'Imax' x,y pairs :{ 0 1.21 2 2.1 40 2.1 }
    3. 'Km' x,y pairs :{ 0 0.0157 2 0.0522 40 0.0522 }
    4. 'minimal nutrient concentration' = 600 (umol/g)
    5. 'optimal nutrient concentration' = 1200 (umol/g)
17. 'number of xylem poles' = 40 (noUnit)
18. 'phosphorus'(noUnit)
    1. 'Cmin' = 0.0002 (umol/ml)
    2. 'Efflux' = 1e-06 (umol/cm/day)
    3. 'Imax' = 0.0555 (umol/cm2/day)
    4. 'Km' = 0.00545 (umol/ml)
    5. 'minimal nutrient concentration' = 30 (umol/g)
    6. 'optimal nutrient concentration' = 60 (umol/g)
19. 'potassium'(noUnit)
    1. 'Cmin' = 0.002 (umol/ml)
    2. 'Efflux' = 1e-06 (umol/cm/day)
    3. 'Imax' = 0.467 (umol/cm2/day)
    4. 'Km' = 0.014 (umol/ml)
    5. 'minimal nutrient concentration' = 117 (umol/g)
    6. 'optimal nutrient concentration' = 234 (umol/g)
20. 'radial hydraulic conductivity' x,y pairs :{ 0 0 1 0.000216 10 0.000216 20 0.00025 30 0.000216 40 0.0001 60 0 }
21. 'reduction in respiration due to aerenchyma' x,y pairs :{ 0 0 0.3 0.7 0.6 1 }
22. 'regular topology' = 4 (noUnit)
23. 'relative carbon cost of exudation' x,y pairs :{ 0 5e-06 100 5e-06 }
24. 'relative respiration' x,y pairs :{ 0 0.09 2 0.035 6 0.035 1000 0.035 }
25. 'root class ID' = 102 (noUnit)
26. 'root hair density' x,y pairs :{ 0 2000 1 2000 2 2000 10 2000 30 0 2000 0 }
27. 'root hair diameter' = 0.0005 (cm)
28. 'root hair length' x,y pairs :{ 0 0 1 0 2 0.028 2000 0.028 }
29. 'soil impedence.v2'(cm)=f{'uniform distribution'} minimum=-0.030000 maximum=0.030000
30. 'top boundary' = 1 (noUnit)

2. 'braceroots2'(noUnit)
    1. 'aerenchyma formation' x,y pairs :{ 0 0 3 0 5 0.1 10 0.25 20 0.393 1000 0.393 }
    2. 'bottom boundary' = 0 (noUnit)
    3. 'bounce of the side' = 0 (noUnit)
    4. 'branch list'(noUnit)
        1. 'lateral of crown roots'(noUnit)
            1. 'allow branches to form above ground' = 0 (noUnit)
            2. 'branching frequency'(cm)=f{'uniform distribution'} minimum=0.100000 maximum=0.400000
            3. 'branching spatial offset' = 15 (cm)
            4. 'length root tip' = 10.93 (cm)
            5. 'number of branches/whorl' = 1 (#)
    5. 'branching angle' = 130 (degrees)
    6. 'cannotgrowup' = 1 (noUnit)
    7. 'copy defaults from' = ../defaultsMajorAxis (noUnit)
    8. 'density' = 0.094 (g/cm3)
    9. 'diameter' x,y pairs :{ 0 0.5 9 0.5 16 0.2 24 0.1 100 0.1 }
    10. 'gravitropism.v2'(cm)=f{'uniform distribution'} minimum=-0.010000 maximum=-0.005000
    11. 'growth rate' x,y pairs :{ 0 0.01 5 1 10 4.5 17 4.5 22 0 1000 0 }
    12. 'length multiplier to diameter multiplier' x,y pairs :{ 0 0.25 1 1 2 1.5 3 1.8 4 2 100 2 1000 2 }
    13. 'length root tip without xylem vessels' = 2 (cm)
    14. 'local resource responses'(noUnit)
        1. 'impact on:branching frequency'(noUnit)

            1. 'aggregation function' = maxRelativeDeviationFromOne (noUnit)
            2. 'impact by:nitrate' x,y pairs :{ 0 1 2000 1 }
            3. 'impact by:phosphorus' x,y pairs :{ 0 0.2 0.015 1 1000 1 }
            4. 'impact by:potassium' x,y pairs :{ 0 1 1000 1 }
        2. 'impact on:gravitropism'(noUnit)
            1. 'aggregation function' = maxRelativeDeviationFromOne (noUnit)
            2. 'impact by:nitrate' x,y pairs :{ 0 1.5 100 1 2000 1 }
            3. 'impact by:phosphorus' x,y pairs :{ 0 0.5 0.015 2 1000 0.5 }
            4. 'impact by:potassium' x,y pairs :{ 0 1 1000 1 }
        3. 'impact on:root potential longitudinal growth'(noUnit)
            1. 'aggregation function' = maxRelativeDeviationFromOne (noUnit)
            2. 'impact by:nitrate' x,y pairs :{ 0 1 2000 1 }
            3. 'impact by:phosphorus' x,y pairs :{ 0 0.2 0.015 1 1000 1 }
            4. 'impact by:potassium' x,y pairs :{ 0 1 1000 1 }
    15. 'longitudinal growth rate multiplier'(cm)=f{'uniform distribution'} minimum=0.700000 maximum=1.000000
    16. 'nitrate'(noUnit)
        1. 'Cmin' = 0.001 (umol/ml)
        2. 'Imax' x,y pairs :{ 0 1.21 2 2.1 40 2.1 }
        3. 'Km' x,y pairs :{ 0 0.0157 2 0.0522 40 0.0522 }
        4. 'minimal nutrient concentration' = 600 (umol/g)
        5. 'optimal nutrient concentration' = 1200 (umol/g)
    17. 'number of xylem poles' = 48 (noUnit)
    18. 'phosphorus'(noUnit)
        1. 'Cmin' = 0.0002 (umol/ml)
        2. 'Efflux' = 1e-06 (umol/cm/day)
        3. 'Imax' = 0.0555 (umol/cm2/day)
        4. 'Km' = 0.00545 (umol/ml)
        5. 'minimal nutrient concentration' = 30 (umol/g)
        6. 'optimal nutrient concentration' = 60 (umol/g)
    19. 'potassium'(noUnit)
        1. 'Cmin' = 0.002 (umol/ml)
        2. 'Efflux' = 1e-06 (umol/cm/day)
        3. 'Imax' = 0.467 (umol/cm2/day)
        4. 'Km' = 0.014 (umol/ml)
        5. 'minimal nutrient concentration' = 117 (umol/g)
        6. 'optimal nutrient concentration' = 234 (umol/g)
    20. 'radial hydraulic conductivity' x,y pairs :{ 0 0 1 0.000216 10 0.000216 20 0.00025 30 0.000216 40 0.0001 60 0 }
    21. 'reduction in respiration due to aerenchyma' x,y pairs :{ 0 0 0.3 0.7 0.6 1 }
    22. 'regular topology' = 3 (noUnit)
    23. 'relative carbon cost of exudation' x,y pairs :{ 0 5e-06 100 5e-06 }
    24. 'relative respiration' x,y pairs :{ 0 0.09 2 0.035 6 0.035 1000 0.035 }
    25. 'root class ID' = 102 (noUnit)
    26. 'root hair density' x,y pairs :{ 0 2000 1 2000 2 2000 10 2000 30 0 2000 0 }
    27. 'root hair diameter' = 0.0005 (cm)
    28. 'root hair length' x,y pairs :{ 0 0 1 0 2 0.028 2000 0.028 }
    29. 'soil impedence.v2'(cm)=f{'uniform distribution'} minimum=-0.030000 maximum=0.030000
    30. 'top boundary' = 1 (noUnit)
3. 'defaults'(noUnit)
    1. 'bottom boundary' = 0 (noUnit)
    2. 'bounce of the side' = 0 (noUnit)
    3. 'cannotgrowup' = 1 (noUnit)
    4. 'local resource responses'(noUnit)
        1. 'impact on:branching frequency'(noUnit)
            1. 'aggregation function' = maxRelativeDeviationFromOne (noUnit)
            2. 'impact by:nitrate' x,y pairs :{ 0 1 2000 1 }
            3. 'impact by:phosphorus' x,y pairs :{ 0 0.5 0.015 1 1000 1 }
            4. 'impact by:potassium' x,y pairs :{ 0 1 1000 1 }
        2. 'impact on:gravitropism'(noUnit)
            1. 'aggregation function' = maxRelativeDeviationFromOne (noUnit)
            2. 'impact by:nitrate' x,y pairs :{ 0 1 2000 1 }
            3. 'impact by:phosphorus' x,y pairs :{ 0 1 1000 1 }

            4. 'impact by:potassium' x,y pairs :{ 0 1 1000 1 }
        3. 'impact on:root potential longitudinal growth'(noUnit)
            1. 'aggregation function' = maxRelativeDeviationFromOne (noUnit)
            2. 'impact by:nitrate' x,y pairs :{ -10 1.5 50 1.5 100 1 2000 1 }
            3. 'impact by:phosphorus' x,y pairs :{ 0 0.2 0.015 1 1000 1 }
            4. 'impact by:potassium' x,y pairs :{ 0 1 1000 1 }
    5. 'phosphorus'(noUnit)
        1. 'Cmin' = 0.0002 (umol/ml)
        2. 'Efflux' = 1e-06 (umol/cm/day)
        3. 'Imax' = 0.0555 (umol/cm2/day)
        4. 'Km' = 0.00545 (umol/ml)
        5. 'minimal nutrient concentration' = 30 (umol/g)
        6. 'optimal nutrient concentration' = 60 (umol/g)
    6. 'potassium'(noUnit)
        1. 'Cmin' = 0.002 (umol/ml)
        2. 'Efflux' = 1e-06 (umol/cm/day)
        3. 'Imax' = 0.467 (umol/cm2/day)
        4. 'Km' = 0.014 (umol/ml)
        5. 'minimal nutrient concentration' = 117 (umol/g)
        6. 'optimal nutrient concentration' = 234 (umol/g)
    7. 'relative carbon cost of exudation' x,y pairs :{ 0 5e-06 100 5e-06 }
    8. 'relative respiration' x,y pairs :{ 0 0.09 2 0.035 6 0.035 1000 0.035 }
    9. 'top boundary' = 1 (noUnit)
4. 'defaults major axis'(noUnit)
    1. 'bottom boundary' = 0 (noUnit)
    2. 'bounce of the side' = 0 (noUnit)
    3. 'cannotgrowup' = 1 (noUnit)
    4. 'copy defaults from' = ../defaults (noUnit)
    5. 'local resource responses'(noUnit)
        1. 'impact on:branching frequency'(noUnit)
            1. 'aggregation function' = maxRelativeDeviationFromOne (noUnit)
            2. 'impact by:nitrate' x,y pairs :{ 0 1 2000 1 }
            3. 'impact by:phosphorus' x,y pairs :{ 0 0.2 0.015 1 1000 1 }
            4. 'impact by:potassium' x,y pairs :{ 0 1 1000 1 }
        2. 'impact on:gravitropism'(noUnit)
            1. 'aggregation function' = maxRelativeDeviationFromOne (noUnit)
            2. 'impact by:nitrate' x,y pairs :{ 0 1.5 100 1 2000 1 }
            3. 'impact by:phosphorus' x,y pairs :{ 0 0.5 0.015 2 1000 0.5 }
            4. 'impact by:potassium' x,y pairs :{ 0 1 1000 1 }
        3. 'impact on:root potential longitudinal growth'(noUnit)
            1. 'aggregation function' = maxRelativeDeviationFromOne (noUnit)
            2. 'impact by:nitrate' x,y pairs :{ 0 1 2000 1 }
            3. 'impact by:phosphorus' x,y pairs :{ 0 0.2 0.015 1 1000 1 }
            4. 'impact by:potassium' x,y pairs :{ 0 1 1000 1 }
    6. 'phosphorus'(noUnit)
        1. 'Cmin' = 0.0002 (umol/ml)
        2. 'Efflux' = 1e-06 (umol/cm/day)
        3. 'Imax' = 0.0555 (umol/cm2/day)
        4. 'Km' = 0.00545 (umol/ml)
        5. 'minimal nutrient concentration' = 30 (umol/g)
        6. 'optimal nutrient concentration' = 60 (umol/g)
    7. 'potassium'(noUnit)
        1. 'Cmin' = 0.002 (umol/ml)
        2. 'Efflux' = 1e-06 (umol/cm/day)
        3. 'Imax' = 0.467 (umol/cm2/day)
        4. 'Km' = 0.014 (umol/ml)
        5. 'minimal nutrient concentration' = 117 (umol/g)
        6. 'optimal nutrient concentration' = 234 (umol/g)
    8. 'relative carbon cost of exudation' x,y pairs :{ 0 5e-06 100 5e-06 }
    9. 'relative respiration' x,y pairs :{ 0 0.09 2 0.035 6 0.035 1000 0.035 }
    10. 'top boundary' = 1 (noUnit)
5. 'finelateral'(noUnit)
    1. 'aerenchyma formation' x,y pairs :{ 0 0 3 0 5 0.1 10 0.25 20 0.393 1000 0.393 }

    2. 'branch list'(noUnit)
        1. 'finelateral2'(noUnit)
            1. 'allow branches to form above ground' = 0 (noUnit)
            2. 'branching frequency'(cm)=f{'uniform distribution'} minimum=0.400000 maximum=0.600000
            3. 'length root tip' = 1.5 (cm)
    3. 'branching angle' = 62.83 (degrees)
    4. 'density' = 0.094 (g/cm3)
    5. 'diameter' = 0.025 (cm)
    6. 'gravitropism.v2' = 0 0 0 (cm)
    7. 'growth rate' x,y pairs :{ 0 0.01 1 0.35 6 0 1000 0 }
    8. 'length multiplier to diameter multiplier' x,y pairs :{ 0 0.25 1 1 2 1.5 3 1.8 4 2 100 2 1000 2 }
    9. 'length root tip without xylem vessels' = 2 (cm)
    10. 'longitudinal growth rate multiplier'(cm)=f{'normal distribution'} minimum=0.500000 maximum=1.500000 mean=1.000000 stdev=0.100000
    11. 'nitrate'(noUnit)
        1. 'Cmin' = 0.0017 (umol/ml)
        2. 'Imax' = 1.27 (umol/cm2/day)
        3. 'Km' = 0.0027 (umol/ml)
        4. 'minimal nutrient concentration' = 600 (umol/g)
        5. 'optimal nutrient concentration' = 1200 (umol/g)
    12. 'number of xylem poles' = 4 (noUnit)
    13. 'radial hydraulic conductivity' x,y pairs :{ 0 0 1 0.000416 60 0.000416 }
    14. 'reduction in respiration due to aerenchyma' x,y pairs :{ 0 0 0.3 0.7 0.6 1 }
    15. 'relative carbon cost of exudation' x,y pairs :{ 0 5e-06 100 1e-06 }
    16. 'root class ID' = 98 (noUnit)
    17. 'root hair density' x,y pairs :{ 0 2000 1 2000 2 2000 10 2000 30 0 2000 0 }
    18. 'root hair diameter' = 0.0005 (cm)
    19. 'root hair length' x,y pairs :{ 0 0 1 0 2 0.028 2000 0.028 }
    20. 'soil impedence.v2'(cm)=f{'uniform distribution'} minimum=-0.050000 maximum=0.050000
6. 'finelateral2'(noUnit)
    1. 'aerenchyma formation' x,y pairs :{ 0 0 3 0 5 0.1 10 0.25 20 0.393 1000 0.393 }
    2. 'branch list'(noUnit)
    3. 'branching angle' = 62.83 (degrees)
    4. 'density' = 0.094 (g/cm3)
    5. 'diameter' = 0.015 (cm)
    6. 'gravitropism.v2' = 0 0 0 (cm)
    7. 'growth rate' x,y pairs :{ 0 0.001 1 0.28 4 0 1000 0 }
    8. 'length multiplier to diameter multiplier' x,y pairs :{ 0 0.25 1 1 2 1.5 3 1.8 4 2 100 2 1000 2 }
    9. 'length root tip without xylem vessels' = 2 (cm)
    10. 'longitudinal growth rate multiplier'(cm)=f{'normal distribution'} minimum=0.500000 maximum=1.500000 mean=1.000000 stdev=0.100000
    11. 'nitrate'(noUnit)
        1. 'Cmin' = 0.0017 (umol/ml)
        2. 'Imax' = 1.27 (umol/cm2/day)
        3. 'Km' = 0.0027 (umol/ml)
        4. 'minimal nutrient concentration' = 600 (umol/g)
        5. 'optimal nutrient concentration' = 1200 (umol/g)
    12. 'number of xylem poles' = 4 (noUnit)
    13. 'radial hydraulic conductivity' x,y pairs :{ 0 0 1 0.000416 60 0.000416 }
    14. 'reduction in respiration due to aerenchyma' x,y pairs :{ 0 0 0.3 0.7 0.6 1 }
    15. 'relative carbon cost of exudation' x,y pairs :{ 0 5e-06 100 1e-06 }
    16. 'root class ID' = 98 (noUnit)
    17. 'root hair density' x,y pairs :{ 0 2000 1 2000 2 2000 10 2000 30 0 2000 0 }
    18. 'root hair diameter' = 0.0005 (cm)
    19. 'root hair length' x,y pairs :{ 0 0 1 0 2 0.028 2000 0.028 }
    20. 'soil impedence.v2'(cm)=f{'uniform distribution'} minimum=-0.050000 maximum=0.050000
7. 'hypocotyl'(noUnit)
    1. 'aerenchyma formation' x,y pairs :{ 0 0 100 0 }
    2. 'bottom boundary' = 0 (noUnit)
    3. 'bounce of the side' = 0 (noUnit)
    4. 'branch list'(noUnit)
        1. 'braceroots'(noUnit)
            1. 'allometric scaling' = 1 (noUnit)

        2. 'branching spatial offset' = 4 (cm)
        3. 'branching time offset' = 25 (day)
        4. 'max number of branches' = 14 (#)
        5. 'number of branches/whorl' = 14 (#)
    2. 'braceroots2'(noUnit)
        1. 'allometric scaling' = 1 (noUnit)
        2. 'branching delay' = 14 (day)
        3. 'branching frequency' = 5 (cm)
        4. 'branching spatial offset' = 7 (cm)
        5. 'branching time offset' = 36 (day)
        6. 'number of branches/whorl' = 20 (#)
    3. 'nodalroots'(noUnit)
        1. 'branching spatial offset' = 1.5 (cm)
        2. 'branching time offset' = 7 (day)
        3. 'max number of branches' = 4 (#)
        4. 'number of branches/whorl' = 4 (#)
    4. 'nodalroots2'(noUnit)
        1. 'allometric scaling' = 1 (noUnit)
        2. 'branching spatial offset' = 1.9 (cm)
        3. 'branching time offset' = 12 (day)
        4. 'max number of branches' = 5 (#)
        5. 'number of branches/whorl' = 5 (#)
    5. 'nodalroots3'(noUnit)
        1. 'allometric scaling' = 1 (noUnit)
        2. 'branching spatial offset' = 2.1 (cm)
        3. 'branching time offset' = 17 (day)
        4. 'max number of branches' = 5 (#)
        5. 'number of branches/whorl' = 5 (#)
    6. 'nodalroots4'(noUnit)
        1. 'allometric scaling' = 1 (noUnit)
        2. 'branching spatial offset' = 2.3 (cm)
        3. 'branching time offset' = 23 (day)
        4. 'max number of branches' = 6 (#)
        5. 'number of branches/whorl' = 6 (#)
5. 'cannotgrowup' = 0 (noUnit)
6. 'copy defaults from' = ../defaultsMajorAxis (noUnit)
7. 'density' = 0.094 (g/cm3)
8. 'diameter' = 0.2 (cm)
9. 'gravitropism.v2' = 0 1 0 (cm)
10. 'growth rate' x,y pairs :{ 0 0 1 0 2 0.87 3 0.91 4 1.53 5 0.35 6 0 1000 0 }
11. 'length root tip without xylem vessels' = 2 (cm)
12. 'local resource responses'(noUnit)
    1. 'impact on:branching frequency'(noUnit)
        1. 'aggregation function' = maxRelativeDeviationFromOne (noUnit)
        2. 'impact by:nitrate' x,y pairs :{ 0 1 10000 1 }
        3. 'impact by:phosphorus' x,y pairs :{ 0 1 1000 1 }
        4. 'impact by:potassium' x,y pairs :{ 0 1 1000 1 }
    2. 'impact on:gravitropism'(noUnit)
        1. 'aggregation function' = maxRelativeDeviationFromOne (noUnit)
        2. 'impact by:nitrate' x,y pairs :{ 0 1 10000 1 }
        3. 'impact by:phosphorus' x,y pairs :{ 0 1 1000 1 }
        4. 'impact by:potassium' x,y pairs :{ 0 1 1000 1 }
    3. 'impact on:root potential longitudinal growth'(noUnit)
        1. 'aggregation function' = maxRelativeDeviationFromOne (noUnit)
        2. 'impact by:nitrate' x,y pairs :{ 0 1 10000 1 }
        3. 'impact by:phosphorus' x,y pairs :{ 0 1 1000 1 }
        4. 'impact by:potassium' x,y pairs :{ 0 1 1000 1 }
13. 'nitrate'(noUnit)
    1. 'Cmin' = 0 (umol/ml)
    2. 'Imax' = 0 (umol/cm2/day)
    3. 'Km' = 1 (umol/ml)
    4. 'minimal nutrient concentration' = 600 (umol/g)
    5. 'optimal nutrient concentration' = 1200 (umol/g)

14. 'number of xylem poles' = 61 (noUnit)
15. 'phosphorus'(noUnit)
    1. 'Cmin' = 0.0002 (umol/ml)
    2. 'Efflux' = 1e-06 (umol/cm/day)
    3. 'Imax' = 0.0555 (umol/cm2/day)
    4. 'Km' = 0.00545 (umol/ml)
    5. 'minimal nutrient concentration' = 30 (umol/g)
    6. 'optimal nutrient concentration' = 60 (umol/g)
16. 'potassium'(noUnit)
    1. 'Cmin' = 0.002 (umol/ml)
    2. 'Efflux' = 1e-06 (umol/cm/day)
    3. 'Imax' = 0.467 (umol/cm2/day)
    4. 'Km' = 0.014 (umol/ml)
    5. 'minimal nutrient concentration' = 117 (umol/g)
    6. 'optimal nutrient concentration' = 234 (umol/g)
17. 'radial hydraulic conductivity' x,y pairs :{ 0 0 60 0 }
18. 'reduction in respiration due to aerenchyma' x,y pairs :{ 0 0 0.3 0.7 0.6 1 }
19. 'relative carbon cost of exudation' x,y pairs :{ 0 0 100 0 }
20. 'relative respiration' x,y pairs :{ 0 0.09 2 0.035 6 0.035 1000 0.035 }
21. 'root class ID' = 97 (noUnit)
22. 'root hair density' x,y pairs :{ 0 0 2000 0 }
23. 'root hair diameter' = 0.0005 (cm)
24. 'root hair length' x,y pairs :{ 0 0 1 0 2 0.028 2000 0.028 }
25. 'soil impedence.v2'(cm)=f{'uniform distribution'} minimum=-0.300000 maximum=0.300000
26. 'top boundary' = 0 (noUnit)

8. 'lateral'(noUnit)
    1. 'aerenchyma formation' x,y pairs :{ 0 0 3 0 5 0.1 10 0.25 20 0.393 1000 0.393 }
    2. 'bottom boundary' = 0 (noUnit)
    3. 'bounce of the side' = 0 (noUnit)
    4. 'branch list'(noUnit)
        1. 'finelateral'(noUnit)
            1. 'allow branches to form above ground' = 0 (noUnit)
            2. 'branching frequency'(cm)=f{'uniform distribution'} minimum=0.150000 maximum=0.350000
            3. 'length root tip' = 4 (cm)
    5. 'branching angle' = 90 (degrees)
    6. 'cannotgrowup' = 1 (noUnit)
    7. 'density' = 0.094 (g/cm3)
    8. 'diameter' = 0.04 (cm)
    9. 'gravitropism.v2' = 0 0 0 (cm)
    10. 'growth rate' x,y pairs :{ 0 0.01 1 0.2 3 0.4 7 1 11 0 1000 0 }
    11. 'length multiplier to diameter multiplier' x,y pairs :{ 0 0.25 1 1 2 1.5 3 1.8 4 2 100 2 1000 2 }
    12. 'length root tip without xylem vessels' = 2 (cm)
    13. 'local resource responses'(noUnit)
        1. 'impact on:branching frequency'(noUnit)
            1. 'aggregation function' = maxRelativeDeviationFromOne (noUnit)
            2. 'impact by:nitrate' x,y pairs :{ 0 1 2000 1 }
            3. 'impact by:phosphorus' x,y pairs :{ 0 0.5 0.015 1 1000 1 }
            4. 'impact by:potassium' x,y pairs :{ 0 1 1000 1 }
        2. 'impact on:gravitropism'(noUnit)
            1. 'aggregation function' = maxRelativeDeviationFromOne (noUnit)
            2. 'impact by:nitrate' x,y pairs :{ 0 1 2000 1 }
            3. 'impact by:phosphorus' x,y pairs :{ 0 1 1000 1 }
            4. 'impact by:potassium' x,y pairs :{ 0 1 1000 1 }
        3. 'impact on:root potential longitudinal growth'(noUnit)
            1. 'aggregation function' = maxRelativeDeviationFromOne (noUnit)
            2. 'impact by:nitrate' x,y pairs :{ -10 1.5 50 1.5 100 1 2000 1 }
            3. 'impact by:phosphorus' x,y pairs :{ 0 0.2 0.015 1 1000 1 }
            4. 'impact by:potassium' x,y pairs :{ 0 1 1000 1 }
    14. 'longitudinal growth rate multiplier'(cm)=f{'lognormal distribution'} minimum=0.100000 maximum=2.000000
    15. 'nitrate'(noUnit)
        1. 'Cmin' = 0.0017 (umol/ml)
        2. 'Imax' = 1.27 (umol/cm2/day)

        3. 'Km' = 0.0027 (umol/ml)
        4. 'minimal nutrient concentration' = 600 (umol/g)
        5. 'optimal nutrient concentration' = 1200 (umol/g)
    16. 'number of xylem poles' = 4 (noUnit)
    17. 'phosphorus'(noUnit)
        1. 'Cmin' = 0.0002 (umol/ml)
        2. 'Efflux' = 1e-06 (umol/cm/day)
        3. 'Imax' = 0.0555 (umol/cm2/day)
        4. 'Km' = 0.00545 (umol/ml)
        5. 'minimal nutrient concentration' = 30 (umol/g)
        6. 'optimal nutrient concentration' = 60 (umol/g)
    18. 'potassium'(noUnit)
        1. 'Cmin' = 0.002 (umol/ml)
        2. 'Efflux' = 1e-06 (umol/cm/day)
        3. 'Imax' = 0.467 (umol/cm2/day)
        4. 'Km' = 0.014 (umol/ml)
        5. 'minimal nutrient concentration' = 117 (umol/g)
        6. 'optimal nutrient concentration' = 234 (umol/g)
    19. 'radial hydraulic conductivity' x,y pairs :{ 0 0 1 0.000416 60 0.000416 }
    20. 'reduction in respiration due to aerenchyma' x,y pairs :{ 0 0 0.3 0.7 0.6 1 }
    21. 'relative carbon cost of exudation' x,y pairs :{ 0 5e-06 100 3e-06 }
    22. 'relative respiration' x,y pairs :{ 0 0.09 2 0.035 6 0.035 1000 0.035 }
    23. 'root class ID' = 98 (noUnit)
    24. 'root hair density' x,y pairs :{ 0 2000 1 2000 2 2000 10 2000 30 0 2000 0 }
    25. 'root hair diameter' = 0.0005 (cm)
    26. 'root hair length' x,y pairs :{ 0 0 1 0 2 0.028 2000 0.028 }
    27. 'soil impedence.v2'(cm)=f{'uniform distribution'} minimum=-0.100000 maximum=0.100000
    28. 'top boundary' = 1 (noUnit)
9. 'lateral of crown roots'(noUnit)
    1. 'aerenchyma formation' x,y pairs :{ 0 0 3 0 5 0.1 10 0.25 20 0.393 1000 0.393 }
    2. 'branch list'(noUnit)
        1. 'lateral'(noUnit)
            1. 'allow branches to form above ground' = 0 (noUnit)
            2. 'branching frequency'(cm)=f{'uniform distribution'} minimum=0.250000 maximum=0.350000
            3. 'length root tip' = 5 (cm)
    3. 'branching angle' = 90 (degrees)
    4. 'density' = 0.094 (g/cm3)
    5. 'diameter' = 0.07 (cm)
    6. 'gravitropism.v2' = 0 0 0 (cm)
    7. 'growth rate' x,y pairs :{ 0 0.1 1 0.5 3 1.2 12 1.2 18 0 1000 0 }
    8. 'length multiplier to diameter multiplier' x,y pairs :{ 0 0.25 1 1 2 1.5 3 1.8 4 2 100 2 1000 2 }
    9. 'length root tip without xylem vessels' = 2 (cm)
    10. 'longitudinal growth rate multiplier'(cm)=f{'normal distribution'} minimum=0.100000 maximum=1.000000 mean=0.400000 stdev=0.300000
    11. 'nitrate'(noUnit)
        1. 'Cmin' = 0.0017 (umol/ml)
        2. 'Imax' = 1.27 (umol/cm2/day)
        3. 'Km' = 0.0027 (umol/ml)
        4. 'minimal nutrient concentration' = 600 (umol/g)
        5. 'optimal nutrient concentration' = 1200 (umol/g)
    12. 'number of xylem poles' = 4 (noUnit)
    13. 'radial hydraulic conductivity' x,y pairs :{ 0 0 1 0.000216 60 0.000216 }
    14. 'reduction in respiration due to aerenchyma' x,y pairs :{ 0 0 0.3 0.7 0.6 1 }
    15. 'relative carbon cost of exudation' x,y pairs :{ 0 5e-06 100 4e-06 }
    16. 'root class ID' = 98 (noUnit)
    17. 'root hair density' x,y pairs :{ 0 2000 1 2000 2 2000 10 2000 30 0 2000 0 }
    18. 'root hair diameter' = 0.0005 (cm)
    19. 'root hair length' x,y pairs :{ 0 0 1 0 2 0.028 2000 0.028 }
    20. 'soil impedence.v2'(cm)=f{'uniform distribution'} minimum=-0.050000 maximum=0.050000
10. 'nodalroots'(noUnit)
    1. 'aerenchyma formation' x,y pairs :{ 0 0 3 0 5 0.1 10 0.25 20 0.393 1000 0.393 }
    2. 'bottom boundary' = 0 (noUnit)
    3. 'bounce of the side' = 0 (noUnit)

4. 'branch list'(noUnit)
    1. 'lateral'(noUnit)
        1. 'allow branches to form above ground' = 0 (noUnit)
        2. 'branching frequency'(cm)=f{'uniform distribution'} minimum=0.237000 maximum=0.437000
        3. 'length root tip' = 10.93 (cm)
5. 'branching angle' = 132.01 (degrees)
6. 'cannotgrowup' = 1 (noUnit)
7. 'copy defaults from' = ../defaultsMajorAxis (noUnit)
8. 'density' = 0.094 (g/cm3)
9. 'diameter' x,y pairs :{ 0 0.084 8 0.083 13 0.096 18 0.113 }
10. 'gravitropism.v2'(cm)=f{'uniform distribution'} minimum=-0.010000 maximum=-0.005000
11. 'growth rate' x,y pairs :{ 0 3.26 1 3.44 2 3.61 3 3.8 4 4 5 4.1 7 4.5 10 5.03 12 5.38 15 5.92 17 6.3 18 6.45 40 6.45 }
12. 'length multiplier to diameter multiplier' x,y pairs :{ 0 0.25 1 1 2 1.5 3 1.8 4 2 100 2 1000 2 }
13. 'length root tip without xylem vessels' = 2 (cm)
14. 'local resource responses'(noUnit)
    1. 'impact on:branching frequency'(noUnit)
        1. 'aggregation function' = maxRelativeDeviationFromOne (noUnit)
        2. 'impact by:nitrate' x,y pairs :{ 0 1 2000 1 }
        3. 'impact by:phosphorus' x,y pairs :{ 0 0.2 0.015 1 1000 1 }
        4. 'impact by:potassium' x,y pairs :{ 0 1 1000 1 }
    2. 'impact on:gravitropism'(noUnit)
        1. 'aggregation function' = maxRelativeDeviationFromOne (noUnit)
        2. 'impact by:nitrate' x,y pairs :{ 0 1.5 100 1 2000 1 }
        3. 'impact by:phosphorus' x,y pairs :{ 0 0.5 0.015 2 1000 0.5 }
        4. 'impact by:potassium' x,y pairs :{ 0 1 1000 1 }
    3. 'impact on:root potential longitudinal growth'(noUnit)
        1. 'aggregation function' = maxRelativeDeviationFromOne (noUnit)
        2. 'impact by:nitrate' x,y pairs :{ 0 1 2000 1 }
        3. 'impact by:phosphorus' x,y pairs :{ 0 0.2 0.015 1 1000 1 }
        4. 'impact by:potassium' x,y pairs :{ 0 1 1000 1 }
15. 'longitudinal growth rate multiplier'(cm)=f{'normal distribution'} minimum=0.600000 maximum=1.200000 mean=1.000000 stdev=0.100000
16. 'nitrate'(noUnit)
    1. 'Cmin' = 0.001 (umol/ml)
    2. 'Imax' x,y pairs :{ 0 1.21 2 2.1 40 2.1 }
    3. 'Km' x,y pairs :{ 0 0.0157 2 0.0522 40 0.0522 }
    4. 'minimal nutrient concentration' = 600 (umol/g)
    5. 'optimal nutrient concentration' = 1200 (umol/g)
17. 'number of xylem poles' = 10 (noUnit)
18. 'phosphorus'(noUnit)
    1. 'Cmin' = 0.0002 (umol/ml)
    2. 'Efflux' = 1e-06 (umol/cm/day)
    3. 'Imax' = 0.0555 (umol/cm2/day)
    4. 'Km' = 0.00545 (umol/ml)
    5. 'minimal nutrient concentration' = 30 (umol/g)
    6. 'optimal nutrient concentration' = 60 (umol/g)
19. 'potassium'(noUnit)
    1. 'Cmin' = 0.002 (umol/ml)
    2. 'Efflux' = 1e-06 (umol/cm/day)
    3. 'Imax' = 0.467 (umol/cm2/day)
    4. 'Km' = 0.014 (umol/ml)
    5. 'minimal nutrient concentration' = 117 (umol/g)
    6. 'optimal nutrient concentration' = 234 (umol/g)
20. 'radial hydraulic conductivity' x,y pairs :{ 0 0 1 0.000216 10 0.000216 20 0.00025 30 0.000216 40 0.0001 60 0 }
21. 'reduction in respiration due to aerenchyma' x,y pairs :{ 0 0 0.3 0.7 0.6 1 }
22. 'regular topology' = 3 (noUnit)
23. 'relative carbon cost of exudation' x,y pairs :{ 0 5e-06 100 5e-06 }
24. 'relative respiration' x,y pairs :{ 0 0.09 2 0.035 6 0.035 1000 0.035 }
25. 'root class ID' = 101 (noUnit)
26. 'root hair density' x,y pairs :{ 0 2000 1 2000 2 2000 10 2000 30 0 2000 0 }
27. 'root hair diameter' = 0.0005 (cm)

28. 'root hair length' x,y pairs :{ 0 0 1 0 2 0.028 2000 0.028 }
29. 'soil impedence.v2'(cm)=f{'uniform distribution'} minimum=-0.020000 maximum=0.020000
30. 'top boundary' = 1 (noUnit)
31. 'topology offset' = 0 (noUnit)

11. 'nodalroots2'(noUnit)
    1. 'aerenchyma formation' x,y pairs :{ 0 0 3 0 5 0.1 10 0.25 20 0.393 1000 0.393 }
    2. 'bottom boundary' = 0 (noUnit)
    3. 'bounce of the side' = 0 (noUnit)
    4. 'branch list'(noUnit)
        1. 'lateral'(noUnit)
            1. 'allow branches to form above ground' = 0 (noUnit)
            2. 'branching frequency'(cm)=f{'uniform distribution'} minimum=0.328000 maximum=0.528000
            3. 'length root tip' = 10.93 (cm)
    5. 'branching angle' = 133.24 (degrees)
    6. 'cannotgrowup' = 1 (noUnit)
    7. 'copy defaults from' = ../defaultsMajorAxis (noUnit)
    8. 'density' = 0.094 (g/cm3)
    9. 'diameter' x,y pairs :{ 0 0.126 8 0.105 13 0.115 100 0.115 }
    10. 'gravitropism.v2'(cm)=f{'uniform distribution'} minimum=-0.010000 maximum=-0.005000
    11. 'growth rate' x,y pairs :{ 0 4.09 1 4.24 2 4.39 3 4.54 4 4.69 5 4.84 7 5.14 10 5.59 12 5.89 13 6.04 40 6.04 }
    12. 'length multiplier to diameter multiplier' x,y pairs :{ 0 0.25 1 1 2 1.5 3 1.8 4 2 100 2 1000 2 }
    13. 'length root tip without xylem vessels' = 2 (cm)
    14. 'local resource responses'(noUnit)
        1. 'impact on:branching frequency'(noUnit)
            1. 'aggregation function' = maxRelativeDeviationFromOne (noUnit)
            2. 'impact by:nitrate' x,y pairs :{ 0 1 2000 1 }
            3. 'impact by:phosphorus' x,y pairs :{ 0 0.2 0.015 1 1000 1 }
            4. 'impact by:potassium' x,y pairs :{ 0 1 1000 1 }
        2. 'impact on:gravitropism'(noUnit)
            1. 'aggregation function' = maxRelativeDeviationFromOne (noUnit)
            2. 'impact by:nitrate' x,y pairs :{ 0 1.5 100 1 2000 1 }
            3. 'impact by:phosphorus' x,y pairs :{ 0 0.5 0.015 2 1000 0.5 }
            4. 'impact by:potassium' x,y pairs :{ 0 1 1000 1 }
        3. 'impact on:root potential longitudinal growth'(noUnit)
            1. 'aggregation function' = maxRelativeDeviationFromOne (noUnit)
            2. 'impact by:nitrate' x,y pairs :{ 0 1 2000 1 }
            3. 'impact by:phosphorus' x,y pairs :{ 0 0.2 0.015 1 1000 1 }
            4. 'impact by:potassium' x,y pairs :{ 0 1 1000 1 }
    15. 'longitudinal growth rate multiplier'(cm)=f{'normal distribution'} minimum=0.600000 maximum=1.200000 mean=1.000000 stdev=0.100000
    16. 'nitrate'(noUnit)
        1. 'Cmin' = 0.001 (umol/ml)
        2. 'Imax' x,y pairs :{ 0 1.21 2 2.1 40 2.1 }
        3. 'Km' x,y pairs :{ 0 0.0157 2 0.0522 40 0.0522 }
        4. 'minimal nutrient concentration' = 600 (umol/g)
        5. 'optimal nutrient concentration' = 1200 (umol/g)
    17. 'number of xylem poles' = 18 (noUnit)
    18. 'phosphorus'(noUnit)
        1. 'Cmin' = 0.0002 (umol/ml)
        2. 'Efflux' = 1e-06 (umol/cm/day)
        3. 'Imax' = 0.0555 (umol/cm2/day)
        4. 'Km' = 0.00545 (umol/ml)
        5. 'minimal nutrient concentration' = 30 (umol/g)
        6. 'optimal nutrient concentration' = 60 (umol/g)
    19. 'potassium'(noUnit)
        1. 'Cmin' = 0.002 (umol/ml)
        2. 'Efflux' = 1e-06 (umol/cm/day)
        3. 'Imax' = 0.467 (umol/cm2/day)
        4. 'Km' = 0.014 (umol/ml)
        5. 'minimal nutrient concentration' = 117 (umol/g)
        6. 'optimal nutrient concentration' = 234 (umol/g)
    20. 'radial hydraulic conductivity' x,y pairs :{ 0 0 1 0.000216 10 0.000216 20 0.00025 30 0.000216 40 0.0001 60 0 }

21. 'reduction in respiration due to aerenchyma' x,y pairs :{ 0 0 0.3 0.7 0.6 1 }
22. 'regular topology' = 0 (noUnit)
23. 'relative carbon cost of exudation' x,y pairs :{ 0 5e-06 100 5e-06 }
24. 'relative respiration' x,y pairs :{ 0 0.09 2 0.035 6 0.035 1000 0.035 }
25. 'root class ID' = 101 (noUnit)
26. 'root hair density' x,y pairs :{ 0 2000 1 2000 2 2000 10 2000 30 0 2000 0 }
27. 'root hair diameter' = 0.0005 (cm)
28. 'root hair length' x,y pairs :{ 0 0 1 0 2 0.028 2000 0.028 }
29. 'soil impedence.v2'(cm)=f{'uniform distribution'} minimum=-0.020000 maximum=0.020000
30. 'top boundary' = 1 (noUnit)
31. 'topology offset' = 0 (noUnit)

12. 'nodalroots3'(noUnit)
    1. 'aerenchyma formation' x,y pairs :{ 0 0 3 0 5 0.1 10 0.25 20 0.393 1000 0.393 }
    2. 'bottom boundary' = 0 (noUnit)
    3. 'bounce of the side' = 0 (noUnit)
    4. 'branch list'(noUnit)
        1. 'lateral'(noUnit)
            1. 'allow branches to form above ground' = 0 (noUnit)
            2. 'branching frequency'(cm)=f{'uniform distribution'} minimum=0.100000 maximum=0.300000
            3. 'length root tip' = 10.93 (cm)
    5. 'branching angle' = 124.21 (degrees)
    6. 'cannotgrowup' = 1 (noUnit)
    7. 'copy defaults from' = ../defaultsMajorAxis (noUnit)
    8. 'density' = 0.094 (g/cm3)
    9. 'diameter' x,y pairs :{ 0 0.178 8 0.113 100 0.113 }
    10. 'gravitropism.v2'(cm)=f{'uniform distribution'} minimum=-0.010000 maximum=-0.005000
    11. 'growth rate' x,y pairs :{ 0 2.96 1 2.98 2 3 3 3.01 4 3.03 5 3.05 8 3.1 40 3.1 }
    12. 'length multiplier to diameter multiplier' x,y pairs :{ 0 0.25 1 1 2 1.5 3 1.8 4 2 100 2 1000 2 }
    13. 'length root tip without xylem vessels' = 2 (cm)
    14. 'local resource responses'(noUnit)
        1. 'impact on:branching frequency'(noUnit)
            1. 'aggregation function' = maxRelativeDeviationFromOne (noUnit)
            2. 'impact by:nitrate' x,y pairs :{ 0 1 2000 1 }
            3. 'impact by:phosphorus' x,y pairs :{ 0 0.2 0.015 1 1000 1 }
            4. 'impact by:potassium' x,y pairs :{ 0 1 1000 1 }
        2. 'impact on:gravitropism'(noUnit)
            1. 'aggregation function' = maxRelativeDeviationFromOne (noUnit)
            2. 'impact by:nitrate' x,y pairs :{ 0 1.5 100 1 2000 1 }
            3. 'impact by:phosphorus' x,y pairs :{ 0 0.5 0.015 2 1000 0.5 }
            4. 'impact by:potassium' x,y pairs :{ 0 1 1000 1 }
        3. 'impact on:root potential longitudinal growth'(noUnit)
            1. 'aggregation function' = maxRelativeDeviationFromOne (noUnit)
            2. 'impact by:nitrate' x,y pairs :{ 0 1 2000 1 }
            3. 'impact by:phosphorus' x,y pairs :{ 0 0.2 0.015 1 1000 1 }
            4. 'impact by:potassium' x,y pairs :{ 0 1 1000 1 }
    15. 'longitudinal growth rate multiplier'(cm)=f{'normal distribution'} minimum=0.600000 maximum=1.200000 mean=1.000000 stdev=0.100000
    16. 'nitrate'(noUnit)
        1. 'Cmin' = 0.001 (umol/ml)
        2. 'Imax' x,y pairs :{ 0 1.21 2 2.1 40 2.1 }
        3. 'Km' x,y pairs :{ 0 0.0157 2 0.0522 40 0.0522 }
        4. 'minimal nutrient concentration' = 600 (umol/g)
        5. 'optimal nutrient concentration' = 1200 (umol/g)
    17. 'number of xylem poles' = 24 (noUnit)
    18. 'phosphorus'(noUnit)
        1. 'Cmin' = 0.0002 (umol/ml)
        2. 'Efflux' = 1e-06 (umol/cm/day)
        3. 'Imax' = 0.0555 (umol/cm2/day)
        4. 'Km' = 0.00545 (umol/ml)
        5. 'minimal nutrient concentration' = 30 (umol/g)
        6. 'optimal nutrient concentration' = 60 (umol/g)
    19. 'potassium'(noUnit)
        1. 'Cmin' = 0.002 (umol/ml)

        2. 'Efflux' = 1e-06 (umol/cm/day)
        3. 'Imax' = 0.467 (umol/cm2/day)
        4. 'Km' = 0.014 (umol/ml)
        5. 'minimal nutrient concentration' = 117 (umol/g)
        6. 'optimal nutrient concentration' = 234 (umol/g)
    20. 'radial hydraulic conductivity' x,y pairs :{ 0 0 1 0.000216 10 0.000216 20 0.00025 30 0.000216 40 0.0001 60 0 }
    21. 'reduction in respiration due to aerenchyma' x,y pairs :{ 0 0 0.3 0.7 0.6 1 }
    22. 'regular topology' = 0 (noUnit)
    23. 'relative carbon cost of exudation' x,y pairs :{ 0 5e-06 100 5e-06 }
    24. 'relative respiration' x,y pairs :{ 0 0.09 2 0.035 6 0.035 1000 0.035 }
    25. 'root class ID' = 101 (noUnit)
    26. 'root hair density' x,y pairs :{ 0 2000 1 2000 2 2000 10 2000 30 0 2000 0 }
    27. 'root hair diameter' = 0.0005 (cm)
    28. 'root hair length' x,y pairs :{ 0 0 1 0 2 0.028 2000 0.028 }
    29. 'soil impedence.v2'(cm)=f{'uniform distribution'} minimum=-0.020000 maximum=0.020000
    30. 'top boundary' = 1 (noUnit)
    31. 'topology offset' = 0 (noUnit)
13. 'nodalroots4'(noUnit)
    1. 'aerenchyma formation' x,y pairs :{ 0 0 3 0 5 0.1 10 0.25 20 0.393 1000 0.393 }
    2. 'bottom boundary' = 0 (noUnit)
    3. 'bounce of the side' = 0 (noUnit)
    4. 'branch list'(noUnit)
        1. 'lateral'(noUnit)
            1. 'allow branches to form above ground' = 0 (noUnit)
            2. 'branching frequency'(cm)=f{'uniform distribution'} minimum=0.100000 maximum=0.300000
            3. 'length root tip' = 10.93 (cm)
    5. 'branching angle' = 135.5 (degrees)
    6. 'cannotgrowup' = 1 (noUnit)
    7. 'copy defaults from' = ../defaultsMajorAxis (noUnit)
    8. 'density' = 0.094 (g/cm3)
    9. 'diameter' x,y pairs :{ 0 0.2 10 0.11 100 0.11 }
    10. 'gravitropism.v2'(cm)=f{'uniform distribution'} minimum=-0.010000 maximum=-0.005000
    11. 'growth rate' x,y pairs :{ 0 0.01 1 1 3 4.5 28 4.5 38 0 1000 0 }
    12. 'length multiplier to diameter multiplier' x,y pairs :{ 0 0.25 1 1 2 1.5 3 1.8 4 2 100 2 1000 2 }
    13. 'length root tip without xylem vessels' = 2 (cm)
    14. 'local resource responses'(noUnit)
        1. 'impact on:branching frequency'(noUnit)
            1. 'aggregation function' = maxRelativeDeviationFromOne (noUnit)
            2. 'impact by:nitrate' x,y pairs :{ 0 1 2000 1 }
            3. 'impact by:phosphorus' x,y pairs :{ 0 0.2 0.015 1 1000 1 }
            4. 'impact by:potassium' x,y pairs :{ 0 1 1000 1 }
        2. 'impact on:gravitropism'(noUnit)
            1. 'aggregation function' = maxRelativeDeviationFromOne (noUnit)
            2. 'impact by:nitrate' x,y pairs :{ 0 1.5 100 1 2000 1 }
            3. 'impact by:phosphorus' x,y pairs :{ 0 0.5 0.015 2 1000 0.5 }
            4. 'impact by:potassium' x,y pairs :{ 0 1 1000 1 }
        3. 'impact on:root potential longitudinal growth'(noUnit)
            1. 'aggregation function' = maxRelativeDeviationFromOne (noUnit)
            2. 'impact by:nitrate' x,y pairs :{ 0 1 2000 1 }
            3. 'impact by:phosphorus' x,y pairs :{ 0 0.2 0.015 1 1000 1 }
            4. 'impact by:potassium' x,y pairs :{ 0 1 1000 1 }
    15. 'longitudinal growth rate multiplier'(cm)=f{'normal distribution'} minimum=0.600000 maximum=1.200000 mean=1.000000 stdev=0.100000
    16. 'nitrate'(noUnit)
        1. 'Cmin' = 0.001 (umol/ml)
        2. 'Imax' x,y pairs :{ 0 1.21 2 2.1 40 2.1 }
        3. 'Km' x,y pairs :{ 0 0.0157 2 0.0522 40 0.0522 }
        4. 'minimal nutrient concentration' = 600 (umol/g)
        5. 'optimal nutrient concentration' = 1200 (umol/g)
    17. 'number of xylem poles' = 32 (noUnit)
    18. 'phosphorus'(noUnit)
        1. 'Cmin' = 0.0002 (umol/ml)

        2. 'Efflux' = 1e-06 (umol/cm/day)
        3. 'Imax' = 0.0555 (umol/cm2/day)
        4. 'Km' = 0.00545 (umol/ml)
        5. 'minimal nutrient concentration' = 30 (umol/g)
        6. 'optimal nutrient concentration' = 60 (umol/g)
    19. 'potassium'(noUnit)
        1. 'Cmin' = 0.002 (umol/ml)
        2. 'Efflux' = 1e-06 (umol/cm/day)
        3. 'Imax' = 0.467 (umol/cm2/day)
        4. 'Km' = 0.014 (umol/ml)
        5. 'minimal nutrient concentration' = 117 (umol/g)
        6. 'optimal nutrient concentration' = 234 (umol/g)
    20. 'radial hydraulic conductivity' x,y pairs :{ 0 0 1 0.000216 10 0.000216 20 0.00025 30 0.000216 40 0.0001 60 0 }
    21. 'reduction in respiration due to aerenchyma' x,y pairs :{ 0 0 0.3 0.7 0.6 1 }
    22. 'relative carbon cost of exudation' x,y pairs :{ 0 5e-06 100 5e-06 }
    23. 'relative respiration' x,y pairs :{ 0 0.09 2 0.035 6 0.035 1000 0.035 }
    24. 'root class ID' = 101 (noUnit)
    25. 'root hair density' x,y pairs :{ 0 2000 1 2000 2 2000 10 2000 30 0 2000 0 }
    26. 'root hair diameter' = 0.0005 (cm)
    27. 'root hair length' x,y pairs :{ 0 0 1 0 2 0.028 2000 0.028 }
    28. 'soil impedence.v2'(cm)=f{'uniform distribution'} minimum=-0.020000 maximum=0.020000
    29. 'top boundary' = 1 (noUnit)
14. 'primary root'(noUnit)
    1. 'aerenchyma formation' x,y pairs :{ 0 0 3 0 5 0.1 10 0.25 20 0.393 1000 0.393 }
    2. 'bottom boundary' = 0 (noUnit)
    3. 'bounce of the side' = 0 (noUnit)
    4. 'branch list'(noUnit)
        1. 'lateral'(noUnit)
            1. 'allow branches to form above ground' = 0 (noUnit)
            2. 'branching frequency'(cm)=f{'uniform distribution'} minimum=0.094000 maximum=0.294000
            3. 'length root tip' = 10.93 (cm)
        2. 'seminal'(noUnit)
            1. 'allow branches to form above ground' = 0 (noUnit)
            2. 'branching spatial offset' = 0 (cm)
            3. 'branching time offset' = 1 (day)
            4. 'number of branches/whorl' = 4 (#)
    5. 'branching angle' = 0 (degrees)
    6. 'cannotgrowup' = 1 (noUnit)
    7. 'copy defaults from' = ../defaultsMajorAxis (noUnit)
    8. 'density' = 0.094 (g/cm3)
    9. 'diameter' = 0.088 (cm)
    10. 'gravitropism.v2'(cm)=f{'uniform distribution'} minimum=-0.015000 maximum=-0.005000
    11. 'growth rate' x,y pairs :{ 0 3.27 5 3.63 10 3.99 15 4.35 20 4.71 25 5.07 40 5.07 }
    12. 'length multiplier to diameter multiplier' x,y pairs :{ 0 0.25 1 1 2 1.5 3 1.8 4 2 100 2 1000 2 }
    13. 'length root tip without xylem vessels' = 2 (cm)
    14. 'local resource responses'(noUnit)
        1. 'impact on:branching frequency'(noUnit)
            1. 'aggregation function' = maxRelativeDeviationFromOne (noUnit)
            2. 'impact by:nitrate' x,y pairs :{ 0 1 2000 1 }
            3. 'impact by:phosphorus' x,y pairs :{ 0 0.2 0.015 1 1000 1 }
            4. 'impact by:potassium' x,y pairs :{ 0 1 1000 1 }
        2. 'impact on:gravitropism'(noUnit)
            1. 'aggregation function' = maxRelativeDeviationFromOne (noUnit)
            2. 'impact by:nitrate' x,y pairs :{ 0 1.5 100 1 2000 1 }
            3. 'impact by:phosphorus' x,y pairs :{ 0 0.5 0.015 2 1000 0.5 }
            4. 'impact by:potassium' x,y pairs :{ 0 1 1000 1 }
        3. 'impact on:root potential longitudinal growth'(noUnit)
            1. 'aggregation function' = maxRelativeDeviationFromOne (noUnit)
            2. 'impact by:nitrate' x,y pairs :{ 0 1 2000 1 }
            3. 'impact by:phosphorus' x,y pairs :{ 0 0.2 0.015 1 1000 1 }
            4. 'impact by:potassium' x,y pairs :{ 0 1 1000 1 }
    15. 'nitrate'(noUnit)

        1. 'Cmin' = 0.001 (umol/ml)
        2. 'Imax' x,y pairs :{ 0 2.3 2 1.92 40 1.92 }
        3. 'Km' x,y pairs :{ 0 0.0105 2 0.0161 40 0.0161 }
        4. 'minimal nutrient concentration' = 600 (umol/g)
        5. 'optimal nutrient concentration' = 1200 (umol/g)
    16. 'number of xylem poles' = 8 (noUnit)
    17. 'phosphorus'(noUnit)
        1. 'Cmin' = 0.0002 (umol/ml)
        2. 'Efflux' = 1e-06 (umol/cm/day)
        3. 'Imax' = 0.0555 (umol/cm2/day)
        4. 'Km' = 0.00545 (umol/ml)
        5. 'minimal nutrient concentration' = 30 (umol/g)
        6. 'optimal nutrient concentration' = 60 (umol/g)
    18. 'potassium'(noUnit)
        1. 'Cmin' = 0.002 (umol/ml)
        2. 'Efflux' = 1e-06 (umol/cm/day)
        3. 'Imax' = 0.467 (umol/cm2/day)
        4. 'Km' = 0.014 (umol/ml)
        5. 'minimal nutrient concentration' = 117 (umol/g)
        6. 'optimal nutrient concentration' = 234 (umol/g)
    19. 'radial hydraulic conductivity' x,y pairs :{ 0 0 1 0.000216 10 0.000216 20 0.000216 30 0.000116 40 5e-05 60 0 }
    20. 'reduction in respiration due to aerenchyma' x,y pairs :{ 0 0 0.3 0.7 0.6 1 }
    21. 'relative carbon cost of exudation' x,y pairs :{ 0 5e-06 100 5e-06 }
    22. 'relative respiration' x,y pairs :{ 0 0.09 2 0.035 6 0.035 1000 0.035 }
    23. 'root class ID' = 100 (noUnit)
    24. 'root hair density' x,y pairs :{ 0 2000 1 2000 2 2000 10 2000 30 0 2000 0 }
    25. 'root hair diameter' = 0.0005 (cm)
    26. 'root hair length' x,y pairs :{ 0 0 1 0 2 0.028 2000 0.028 }
    27. 'soil impedence.v2'(cm)=f{'uniform distribution'} minimum=-0.050000 maximum=0.050000
    28. 'top boundary' = 1 (noUnit)
15. 'resources'(noUnit)
    1. 'Cto dry weight ratio' = 0.45 (100%)
    2. 'carbon allocation to leafs factor' x,y pairs :{ 0 1 10 0.7 20 0.45 33 0.42 40 0.4 60 0.4 }
    3. 'carbon allocation to roots factor' x,y pairs :{ 0 1 1 1 6 0.4 20 0.2 40 0.17 80 0.17 }
    4. 'carbon cost of nitrate uptake' = 1.392e-05 (g/umol)
    5. 'max carbon allocation to shoot' = 0.82 (100%)
    6. 'nitrate'(noUnit)
        1. 'initial nutrient uptake' = 383.88 (umol)
    7. 'phosphorus'(noUnit)
        1. 'initial nutrient uptake' = 40.82 (umol)
    8. 'potassium'(noUnit)
        1. 'initial nutrient uptake' = 27 (umol)
    9. 'reserve allocation rate' x,y pairs :{ 0 0.01 1 0.02 2 0.04 3 0.04 10 0.2 11 0.2 1000 0.2 }
    10. 'seed carbohydrate content' = 0.7116 (100%)
    11. 'seed carbohydrate to CFactor' = 0.444 (100%)
    12. 'seed reserve duration' = 100 (day)
    13. 'seed size' = 0.35 (g)
16. 'seminal'(noUnit)
    1. 'aerenchyma formation' x,y pairs :{ 0 0 3 0 5 0.1 10 0.25 20 0.393 1000 0.393 }
    2. 'bottom boundary' = 0 (noUnit)
    3. 'bounce of the side' = 0 (noUnit)
    4. 'branch list'(noUnit)
        1. 'lateral'(noUnit)
            1. 'allow branches to form above ground' = 0 (noUnit)
            2. 'branching frequency'(cm)=f{'uniform distribution'} minimum=0.282000 maximum=0.482000
            3. 'length root tip' = 10.93 (cm)
    5. 'branching angle' = 119.64 (degrees)
    6. 'cannotgrowup' = 1 (noUnit)
    7. 'copy defaults from' = ../defaultsMajorAxis (noUnit)
    8. 'density' = 0.094 (g/cm3)
    9. 'diameter' = 0.074 (cm)
    10. 'gravitropism.v2'(cm)=f{'uniform distribution'} minimum=-0.035000 maximum=-0.025000

11. 'growth rate' x,y pairs :{ 0 1.29 1 1.51 2 1.72 3 1.94 4 2.15 5 2.37 7 2.8 10 3.44 12 3.87 15 4.52 17 4.95 20 5.59 22 6.02 25 6.67 40 6.67 }
12. 'length multiplier to diameter multiplier' x,y pairs :{ 0 0.25 1 1 2 1.5 3 1.8 4 2 100 2 1000 2 }
13. 'length root tip without xylem vessels' = 2 (cm)
14. 'local resource responses'(noUnit)
    1. 'impact on:branching frequency'(noUnit)
        1. 'aggregation function' = maxRelativeDeviationFromOne (noUnit)
        2. 'impact by:nitrate' x,y pairs :{ 0 1 2000 1 }
        3. 'impact by:phosphorus' x,y pairs :{ 0 0.2 0.015 1 1000 1 }
        4. 'impact by:potassium' x,y pairs :{ 0 1 1000 1 }
    2. 'impact on:gravitropism'(noUnit)
        1. 'aggregation function' = maxRelativeDeviationFromOne (noUnit)
        2. 'impact by:nitrate' x,y pairs :{ 0 1.5 100 1 2000 1 }
        3. 'impact by:phosphorus' x,y pairs :{ 0 0.5 0.015 2 1000 0.5 }
        4. 'impact by:potassium' x,y pairs :{ 0 1 1000 1 }
    3. 'impact on:root potential longitudinal growth'(noUnit)
        1. 'aggregation function' = maxRelativeDeviationFromOne (noUnit)
        2. 'impact by:nitrate' x,y pairs :{ 0 1 2000 1 }
        3. 'impact by:phosphorus' x,y pairs :{ 0 0.2 0.015 1 1000 1 }
        4. 'impact by:potassium' x,y pairs :{ 0 1 1000 1 }
15. 'longitudinal growth rate multiplier'(cm)=f{'normal distribution'} minimum=0.600000 maximum=1.200000 mean=1.000000 stdev=0.100000
16. 'nitrate'(noUnit)
    1. 'Cmin' = 0.001 (umol/ml)
    2. 'Imax' x,y pairs :{ 0 2.3 2 1.92 40 1.92 }
    3. 'Km' x,y pairs :{ 0 0.0105 2 0.0161 40 0.0161 }
    4. 'minimal nutrient concentration' = 600 (umol/g)
    5. 'optimal nutrient concentration' = 1200 (umol/g)
17. 'number of xylem poles' = 6 (noUnit)
18. 'phosphorus'(noUnit)
    1. 'Cmin' = 0.0002 (umol/ml)
    2. 'Efflux' = 1e-06 (umol/cm/day)
    3. 'Imax' = 0.0555 (umol/cm2/day)
    4. 'Km' = 0.00545 (umol/ml)
    5. 'minimal nutrient concentration' = 30 (umol/g)
    6. 'optimal nutrient concentration' = 60 (umol/g)
19. 'potassium'(noUnit)
    1. 'Cmin' = 0.002 (umol/ml)
    2. 'Efflux' = 1e-06 (umol/cm/day)
    3. 'Imax' = 0.467 (umol/cm2/day)
    4. 'Km' = 0.014 (umol/ml)
    5. 'minimal nutrient concentration' = 117 (umol/g)
    6. 'optimal nutrient concentration' = 234 (umol/g)
20. 'radial hydraulic conductivity' x,y pairs :{ 0 0 1 0.000216 10 0.000216 20 0.00025 30 0.000216 40 0.0001 60 0 }
21. 'reduction in respiration due to aerenchyma' x,y pairs :{ 0 0 0.3 0.7 0.6 1 }
22. 'regular topology' = 1 (noUnit)
23. 'relative carbon cost of exudation' x,y pairs :{ 0 5e-06 100 5e-06 }
24. 'relative respiration' x,y pairs :{ 0 0.09 2 0.035 6 0.035 1000 0.035 }
25. 'root class ID' = 99 (noUnit)
26. 'root hair density' x,y pairs :{ 0 2000 1 2000 2 2000 10 2000 30 0 2000 0 }
27. 'root hair diameter' = 0.0005 (cm)
28. 'root hair length' x,y pairs :{ 0 0 1 0 2 0.028 2000 0.028 }
29. 'soil impedence.v2'(cm)=f{'uniform distribution'} minimum=-0.040000 maximum=0.040000
30. 'top boundary' = 1 (noUnit)

17. 'shoot'(noUnit)
    1. 'aerenchyma photosynthesis mitigation' = 0.5 (100%)
    2. 'area per plant' = 1600 (cm2)
    3. 'extinction coefficient' = 0.85 (noUnit)
    4. 'leaf area expantion rate' x,y pairs :{ 0 0 2 0 5 0 6 2.58 7 5.46 8 8.34 9 11.22 10 14.1 11 16.98 12 19.86 13 30.87 14 35.65 15 40.43 16 45.21 17 49.99 18 54.77 19 59.55 20 64.33 21 69.11 22 73.89 23 78.67 24 83.45 25 88.23 40 88.23 }
    5. 'light use efficiency' = 3.8e-07 (g/umol)

    6. 'nitrate'(noUnit)
        1. 'leaf minimal nutrient concentration' x,y pairs :{ 0 1200 80 800 }
        2. 'leaf optimal nutrient concentration' x,y pairs :{ 0 2500 80 1500 }
        3. 'stem minimal nutrient concentration' = 400 (umol/g)
        4. 'stem optimal nutrient concentration' = 800 (umol/g)
    7. 'phosphorus'(noUnit)
        1. 'leaf minimal nutrient concentration' = 35 (umol/g)
        2. 'leaf optimal nutrient concentration' = 70 (umol/g)
        3. 'stem minimal nutrient concentration' = 15 (umol/g)
        4. 'stem optimal nutrient concentration' = 30 (umol/g)
    8. 'potassium'(noUnit)
        1. 'leaf minimal nutrient concentration' = 273 (umol/g)
        2. 'leaf optimal nutrient concentration' = 508 (umol/g)
        3. 'stem minimal nutrient concentration' = 117 (umol/g)
        4. 'stem optimal nutrient concentration' = 250 (umol/g)
    9. 'relative potential transpiration' = 100 (cm3/g)
    10. 'relative respiration rate leafs' = 0.04 (g/g/day)
    11. 'relative respiration rate stems' = 0.02 (g/g/day)
    12. 'specific leaf area' x,y pairs :{ 0 0.0015 24 0.0026 50 0.0032 100 0.0032 }
18. 'stress impact factors'(noUnit)
    1. 'impact on:branching frequency multiplier'(noUnit)
        1. 'impact by:nitrate' x,y pairs :{ 0 1 0.5 1 1 0 }
        2. 'impact by:phosphorus' x,y pairs :{ 0 1 0.5 1 1 0 }
        3. 'impact by:potassium' x,y pairs :{ 0 1 0.5 1 1 0 }
    2. 'impact on:gravitropism multiplier'(noUnit)
        1. 'impact by:nitrate' x,y pairs :{ 0 1 0.5 1 1 0 }
        2. 'impact by:phosphorus' x,y pairs :{ 0 1 0.5 1 1 0 }
        3. 'impact by:potassium' x,y pairs :{ 0 1 0.5 1 1 0 }
    3. 'impact on:leaf area expantion rate'(noUnit)
        1. 'impact by:nitrate' x,y pairs :{ 0 0 0.3 0.1 1 1 }
        2. 'impact by:phosphorus' x,y pairs :{ 0 0 1 1 }
        3. 'impact by:potassium' x,y pairs :{ 0 0 0.2 0.5 1 1 }
    4. 'impact on:leaf respiration'(noUnit)
        1. 'impact by:nitrate' x,y pairs :{ 0 1 1 1 }
        2. 'impact by:phosphorus' x,y pairs :{ 0 1 1 1 }
        3. 'impact by:potassium' x,y pairs :{ 0 1 1 1 }
    5. 'impact on:photosynthesis'(noUnit)
        1. 'impact by:nitrate' x,y pairs :{ 0 0 0.4 0.5 1 1 }
        2. 'impact by:phosphorus' x,y pairs :{ 0 0.5 0.5 1 1 1 }
        3. 'impact by:potassium' x,y pairs :{ 0 0 1 1 }
    6. 'impact on:root potential longitudinal growth'(noUnit)
        1. 'impact by:nitrate' x,y pairs :{ 0 0 0.5 1 1 1 }
        2. 'impact by:phosphorus' x,y pairs :{ 0 1 0.1 1 1 1 }
        3. 'impact by:potassium' x,y pairs :{ 0 0 0.5 1 1 1 }
    7. 'impact on:root potential longitudinal growth multiplier'(noUnit)
        1. 'impact by:nitrate' x,y pairs :{ 0 1 0.5 1 1 0 }
        2. 'impact by:phosphorus' x,y pairs :{ 0 1 0.5 1 1 0 }
        3. 'impact by:potassium' x,y pairs :{ 0 1 0.5 1 1 0 }
    8. 'impact on:root segment carbon cost of exudates'(noUnit)
        1. 'impact by:nitrate' x,y pairs :{ 0 1 1 1 }
        2. 'impact by:phosphorus' x,y pairs :{ 0 1 1 1 }
        3. 'impact by:potassium' x,y pairs :{ 0 1 1 1 }
    9. 'impact on:root segment respiration'(noUnit)
        1. 'impact by:nitrate' x,y pairs :{ 0 1 1 1 }
        2. 'impact by:phosphorus' x,y pairs :{ 0 1 1 1 }
        3. 'impact by:potassium' x,y pairs :{ 0 1 1 1 }
    10. 'impact on:root segment secondary growth'(noUnit)
        1. 'impact by:nitrate' x,y pairs :{ 0 0 1 1 }
        2. 'impact by:phosphorus' x,y pairs :{ 0 0 1 1 }
        3. 'impact by:potassium' x,y pairs :{ 0 0 1 1 }
    11. 'impact on:stem respiration'(noUnit)
        1. 'impact by:nitrate' x,y pairs :{ 0 1 1 1 }
        2. 'impact by:phosphorus' x,y pairs :{ 0 1 1 1 }

        3. 'impact by:potassium' x,y pairs :{ 0 1 1 1 }
7. 'shoot template'(noUnit)
    1. 'area per plant'
    2. 'carbon allocation to leafs'(g) initial value = 0 0 0
    3. 'carbon allocation to stems'(g) initial value = 0 0 0
    4. 'extinction coefficient'
    5. 'leaf area'(cm2) initial value = 0 0 0
    6. 'leaf area index'(cm2/cm2)
    7. 'leaf area reduction coefficient'(cm2/cm2)
    8. 'leaf dry weight'(g) initial value = 0 0 0
    9. 'leaf potential carbon sink for growth'(g) initial value = 0 0 0
    10. 'leaf respiration'(g) initial value = 0 0 0
    11. 'light interception'(umol/cm2/day)
    12. 'photosynthesis'(g) initial value = 0 0 0
    13. 'potential leaf area'(cm2) initial value = 0 0 0
    14. 'relative carbon allocation to leafs'(100%)
    15. 'relative carbon allocation to stems'(100%)
    16. 'stem dry weight'(g) initial value = 0 0 0
    17. 'stem potential carbon sink for growth'(g) initial value = 0 0 0
    18. 'stem respiration'(g) initial value = 0 0 0
    19. 'stress adjusted potential leaf area'(cm2) initial value = 0 0 0
8. 'sibling root template'(noUnit)
    1. 'branches'(noUnit)
    2. 'data points'(noUnit)
    3. 'growthpoint'(cm) initial position = 0 0 0 0 0 0 0
        1. 'branching frequency multiplier'(noUnit)
        2. 'gravitropism'
            1. 'multiplier'(noUnit)
        3. 'root circumference'(cm)
        4. 'root diameter'(cm)
        5. 'root longitudinal growth'(cm) initial value = 0 0 0
        6. 'root potential longitudinal growth'(cm) initial value = 0 0 0
            1. 'rate multiplier'(noUnit)
        7. 'root potential secondary growth'(cm)
        8. 'root segment age' = 0 (day)
        9. 'root segment carbon cost of exudates' = 0 (g)
        10. 'root segment dry weight' = 0 (g)
        11. 'root segment length' = 0 (cm)
        12. 'root segment length duration' = 0 (cm.day)
        13. 'root segment potential carbon sink for growth'(g) initial value = 0 0 0
        14. 'root segment respiration' = 0 (g)
        15. 'root segment secondary potential carbon sink for growth' = 0 (g)
        16. 'root segment specific weight' = 0 (g/cm3)
        17. 'root segment surface area' = 0 (cm2)
        18. 'root segment volume' = 0 (cm3)
    4. 'root carbon cost of exudates'(g) initial value = 0 0 0
    5. 'root dry weight'(g)
    6. 'root length'(cm)
    7. 'root respiration'(g) initial value = 0 0 0
    8. 'root secondary potential carbon sink for growth'(g) initial value = 0 0 0
    9. 'root surface area'(cm2)
    10. 'root system carbon cost of exudates'(g) initial value = 0 0 0
    11. 'root system dry weight'(g)
    12. 'root system length'(cm)
    13. 'root system longitudinal growth'(cm) initial value = 0 0 0
    14. 'root system potential carbon sink for growth'(g) initial value = 0 0 0
    15. 'root system potential carbon sink for growth;major axis'(g) initial value = 0 0 0
        1. 'included root classes' = hypocotyl, primaryRoot, seminal, nodalroots, nodalroots1, nodalroots2, nodalroots3, nodalroots4, nodalroots5, braceroots, braceroots1, braceroots2, braceroots3, basalWhorl1, basalWhorl2, basalWhorl3, basalWhorl4 (noUnit)
    16. 'root system respiration'(g) initial value = 0 0 0
    17. 'root system secondary potential carbon sink for growth'(g) initial value = 0 0 0
    18. 'root system surface area'(cm2)

        19. 'root system volume'(cm3)
        20. 'root volume'(cm3)
9. 'simulation controls'(noUnit)
    1. 'Simula stochastic::number of samples' = 1 (noUnit)
    2. 'aerenchyma'(noUnit)
        1. 'include photosynthesis effects' = 0 (noUnit)
        2. 'reduce respiration' = 0 (noUnit)
        3. 'remobilize p' = 0 (noUnit)
    3. 'integration parameters'(noUnit)
        1. 'default spatial integration length' = 1 (cm)
    4. 'output parameters'(noUnit)
        1. 'RSML'(noUnit)
            1. 'requested variables' = rootLength, rootSurfaceArea (noUnit)
            2. 'run' = 1 (noUnit)
            3. 'time interval' = 2 (noUnit)
        2. 'VTU'(noUnit)
            1. 'include point data' = 1 (noUnit)
            2. 'include roots' = 1 (noUnit)
            3. 'include shoots' = 0 (noUnit)
            4. 'include VTUFor depletion zones' = 1 (noUnit)
            5. 'run' = 1 (noUnit)
            6. 'time interval' = 1 (noUnit)
        3. 'defaults'(noUnit)
            1. 'end time' = 25 (noUnit)
            2. 'start time' = 0 (noUnit)
            3. 'time interval' = 1 (noUnit)
        4. 'garbage collection'(noUnit)
            1. 'lack time' = 1 (noUnit)
            2. 'run' = 1 (noUnit)
            3. 'start time' = 5 (noUnit)
            4. 'time interval' = 1 (noUnit)
        5. 'model dump'(noUnit)
            1. 'end time' = 1 (noUnit)
            2. 'run' = 1 (noUnit)
            3. 'start time' = 0 (noUnit)
        6. 'probe all objects'(noUnit)
            1. 'run' = 0 (noUnit)
            2. 'time interval' = 1 (noUnit)
        7. 'raster image'(noUnit)
            1. 'edge smoothing' = 0 (cm)
            2. 'include binary raw image' = 0 (noUnit)
            3. 'include VTKImage' = 0 (noUnit)
            4. 'overlap function' = additive (noUnit)
            5. 'radius comes from' = rootDiameter,phosphorus/radiusDepletionZone (noUnit)
            6. 'root diameter scaling factor' = 1 (100%)
            7. 'run' = 0 (noUnit)
            8. 'start time' = 5 (noUnit)
            9. 'substeps' = 1 (#)
            10. 'time interval' = 5 (noUnit)
            11. 'voxelsize' = 0.1 (cm)
        8. 'root statistics'(noUnit)
            1. 'requested variables' = rootLength, rootSurfaceArea (noUnit)
            2. 'run' = 0 (noUnit)
            3. 'time interval' = 5 (noUnit)
        9. 'table'(noUnit)
            1. 'run' = 1 (noUnit)
            2. 'searching depth' = 5 (noUnit)
            3. 'skip these variables' = primaryRoot, hypocotyl, (noUnit)
        10. 'table of root nodes'(noUnit)
            1. 'run' = 0 (noUnit)
            2. 'time interval' = 8 (noUnit)
        11. 'vtp'(noUnit)
            1. 'run' = 1 (noUnit)

        2. 'time interval' = 1 (noUnit)
10. 'soil'(noUnit)
    1. 'Root length profile_00-10'(cm)
        1. 'y1' = 0 (noUnit)
        2. 'y2' = -10 (noUnit)
    2. 'Root length profile_10-20'(cm)
        1. 'y1' = -10 (noUnit)
        2. 'y2' = -20 (noUnit)
    3. 'Root length profile_ to 0-30'(cm)
        1. 'y1' = -20 (noUnit)
        2. 'y2' = -30 (noUnit)
    4. 'Root length profile_30-40'(cm)
        1. 'y1' = -30 (noUnit)
        2. 'y2' = -40 (noUnit)
    5. 'Root length profile_ for 0-50'(cm)
        1. 'y1' = -40 (noUnit)
        2. 'y2' = -50 (noUnit)
    6. 'Root length profile_50-60'(cm)
        1. 'y1' = -50 (noUnit)
        2. 'y2' = -60 (noUnit)
    7. 'Root length profile_60-70'(cm)
        1. 'y1' = -60 (noUnit)
        2. 'y2' = -70 (noUnit)
    8. 'Root length profile_70-80'(cm)
        1. 'y1' = -70 (noUnit)
        2. 'y2' = -80 (noUnit)
    9. 'Root length profile_80-90'(cm)
        1. 'y1' = -80 (noUnit)
        2. 'y2' = -90 (noUnit)
    10. 'Root length profile_90+'(cm)
        1. 'y1' = -90 (noUnit)
        2. 'y2' = -1000 (noUnit)
    11. 'between the row coring'(noUnit)
        1. '05.500,00.000'(cm)
            1. 'center' = 5.5 0 0 (cm)
            2. 'coring depth' = -200 (cm)
            3. 'radius' = 4.5 (cm)
            4. 'vertical spacing' = 10 (cm)
    12. 'in the row coring'(noUnit)
        1. '00.000,00.000'(cm)
            1. 'center' = 0 0 0 (cm)
            2. 'coring depth' = -200 (cm)
            3. 'radius' = 4.5 (cm)
            4. 'vertical spacing' = 10 (cm)

#### *OpenSimRoot Zea mays* ssp. *parviglumis* (Ames 21803) Parametrization

*OpenSimRoot* uses a hierarchical input file which is summarized below. The hierarchy gives the parameters context. For example, the parameter 'specific leaf area' belongs to the shoot of a specific plant. In *OpenSimRoot*, parameters can be a single value, a value drawn from a distribution (random), or the result of an interpolation table. For constants we give the value, for distributions the distribution parameters and for the tables a list of space separated values e.g. x1 y1 x2 y2 .... xn yn.

1. 'data point template'(noUnit)
   1. 'aerenchyma formation'
   2. 'branching frequency multiplier'(noUnit)
   3. 'combined root class ID'(noUnit)
   4. 'parent root class ID'(noUnit)
   5. 'root circumference'(cm)
   6. 'root class ID'(noUnit)
   7. 'root diameter'(cm) initial value = 0 0 0
   8. 'root hair density'
   9. 'root hair diameter'
   10. 'root hair length'
   11. 'root hair surface area'(cm2/cm)
   12. 'root length to base'(cm)
   13. 'root potential secondary growth'(cm) initial value = 0 0 0
   14. 'root segment age'(day)
   15. 'root segment carbon cost of exudates'(g) initial value = 0 0 0
   16. 'root segment dry weight'(g)
   17. 'root segment length'(cm)
   18. 'root segment length duration'(cm.day) initial value = 0 0 0
   19. 'root segment respiration'(g/day)
   20. 'root segment secondary potential carbon sink for growth'(g) initial value = 0 0 0
   21. 'root segment surface area'(cm2)
   22. 'root segment volume'(cm3)
   23. 'spatial root density'(cm/cm3)
2. 'environment'(noUnit)
   1. 'atmosphere'(noUnit)
      1. 'PAR/ RDD' = 1 (100%)
      2. 'actual durationof sunshine' x,y pairs :{ 0 0 60 0 }
      3. 'albedo crop' = 0.23 (noUnit)
      4. 'albedo soil' = 0.17 (noUnit)
      5. 'altitude' = 91 (m)
      6. 'average daily temperature' x,y pairs :{ 0 16.9 100 16.9 }
      7. 'evaporation' x,y pairs :{ 0 0 1 0.05 2 0.1 3 0.1 4 0.05 5 0.05 6 0.1 7 0.05 8 0.05 9 0.1 10 0.1 11 0.05 12 0.1 13 0.1 14 0.05 15 0.04 16 0.03 17 0.02 18 0.09 19 0.09 20 0.04 21 0.09 22 0.09 23 0.04 24 0.03 25 0.02 26 0.02 27 0.08 28 0.03 29 0.08 30 0.03 31 0.08 32 0.07 33 0.07 34 0.07 35 0.03 36 0.02 37 0.01 38 0 39 0 40 0 41 0 42 0.06 }
      8. 'irradiation' = 4000 (umol/cm2/day)
      9. 'latitude' = 50.8 (noUnit)
      10. 'net radiation' = 50 (W/m2)
      11. 'net radiation soil' = 0 (W/m2)
      12. 'precipitation' x,y pairs :{ 0 0 1 0 2 1 3 0.29 4 0 5 0 6 0.61 7 0 8 0 9 0.25 10 0.03 11 0 12 0.64 13 0.33 14 0 15 0 16 0 17 0 18 1.8 19 0.2 20 0 21 2.84 22 0.38 23 0 24 0 25 0 26 0 27 0.18 28 0 29 0.46 30 0 31 1.35 32 0.13 33 0.23 34 0.25 35 0 36 0 37 0 38 0 39 0 40 0 41 0 42 1.42 }
      13. 'relative humidity' = 60 (m/s)
      14. 'start day' = 22 (noUnit)
      15. 'start month' = 6 (noUnit)
      16. 'wind speed' = 2 (m/s)
   2. 'dimensions'(noUnit)
      1. 'max corner' = 13 0 30 (cm)
      2. 'min corner' = -13 -150 -30 (cm)
      3. 'resolution' = 1 1 1 (cm)
   3. 'soil'(noUnit)
      1. 'bulk density' x,y pairs :{ -200 1.51 -65 1.51 -47 1.4 -30 1.42 -16 1.29 -5 1.24 0 1.24 }
      2. 'nitrate'(noUnit)
         1. 'adsorption coefficient' = 0 (umol/cm)
         2. 'buffer power' x,y pairs :{ -1000 0.4 1000 0.4 }

            3. 'concentration' x,y pairs :{ -1000 1.59 -55 1.59 -45 1.67 -35 2.17 -25 3.15 -15 4.02 -5 2.36 0 2.8 0.01 0 100 0 }
            4. 'diffusion coefficient' x,y pairs :{ -1000 0.07 -0 0.07 1e-05 1e-08 1000 1e-08 }
            5. 'increase time step' = 1 (noUnit)
            6. 'longitudinal dispersivity' = 1 (cm)
            7. 'r1-r0' = 4 (cm)
            8. 'saturated diffusion coefficient' = 1.6416 (cm2/day)
            9. 'transverse dispersivity' = 0.5 (cm)
        3. 'organic'(noUnit)
            1. 'CNRatio microbes' = 10 (g/g)
            2. 'CNratio' x,y pairs :{ -10000 13 0 13 }
            3. 'assimilation efficiency microbes' = 1 (noUnit)
            4. 'carbon content' x,y pairs :{ -200 0.005 -40 0.005 -30 0.01 -10 0.02 0 0.02 }
            5. 'initial relative mineralisation rate' x,y pairs :{ -1000 0 -25 0 -10 0.037 0 0.037 }
            6. 'speed of aging' = 0.46 (noUnit)
            7. 'time offset' = 30 (day)
        4. 'phosphorus'(noUnit)
            1. 'adsorption coefficient' = 1333.3 (umol/cm)
            2. 'buffer power' x,y pairs :{ -1000 400 1000 400 }
            3. 'concentration' x,y pairs :{ -1000 0.00024 -30 0.00025 -29 0.00175 0 0.00175 0.0001 0 1000 0 }
            4. 'diffusion coefficient' x,y pairs :{ -1000 0.00019872 1000 0.00019872 }
            5. 'increase time step' = 1.1 (noUnit)
            6. 'longitudinal dispersivity' = 0 (cm)
            7. 'r1-r0' = 0.3 (cm)
            8. 'saturated diffusion coefficient' = 0.00495 (cm2/day)
            9. 'transverse dispersivity' = 0 (cm)
        5. 'potassium'(noUnit)
            1. 'adsorption coefficient' = 33.3 (umol/cm)
            2. 'buffer power' x,y pairs :{ -1000 10 1000 10 }
            3. 'concentration' x,y pairs :{ -1000 0.05 -30 0.05 -29 0.15 0 0.15 1e-05 0 1000 0 }
            4. 'diffusion coefficient' x,y pairs :{ -1000 0.067 1000 0.067 }
            5. 'increase time step' = 1.01 (noUnit)
            6. 'longitudinal dispersivity' = 1 (cm)
            7. 'r1-r0' = 1.5 (cm)
            8. 'saturated diffusion coefficient' = 1.56 (cm2/day)
            9. 'transverse dispersivity' = 0.5 (cm)
        6. 'water'(noUnit)
            1. 'initial hydraulic head' x,y pairs :{ -200 -10 -150 -52 -60 -143 0 -215 }
            2. 'residual water content' x,y pairs :{ -300 0.067 0 0.067 }
            3. 'saturated conductivity' x,y pairs :{ -300 1 -150 1 -140 10.8 0 10.8 }
            4. 'saturated water content' x,y pairs :{ -300 0.39 -65 0.39 -35 0.39 -25 0.43 -15 0.45 0 0.46 }
            5. 'van genuchten:alpha' x,y pairs :{ -300 0.02 0 0.02 }
            6. 'van genuchten:n' x,y pairs :{ -300 1.41 0 1.41 }
            7. 'volumetric water content in barber cushman' = 0.3 (cm3/cm3)
3. 'hypocotyl template'(noUnit)
    1. 'branches'(noUnit)
    2. 'data points'(noUnit)
    3. 'growthpoint'(cm) initial position = 0 0 0 0 0 0 0
        1. 'branching frequency multiplier'(noUnit)
        2. 'root circumference'(cm)
        3. 'root diameter'(cm)
        4. 'root longitudinal growth'(cm) initial value = 0 0 0
        5. 'root potential longitudinal growth'(cm) initial value = 0 0 0
        6. 'root potential secondary growth'(cm)
        7. 'root segment age' = 0 (day)
        8. 'root segment carbon cost of exudates' = 0 (g)
        9. 'root segment dry weight' = 0 (g)
        10. 'root segment length' = 0 (cm)
        11. 'root segment length duration' = 0 (cm.day)
        12. 'root segment potential carbon sink for growth'(g) initial value = 0 0 0
        13. 'root segment respiration' = 0 (g)
        14. 'root segment secondary potential carbon sink for growth' = 0 (g)
        15. 'root segment surface area' = 0 (cm2)

        16. 'root segment volume' = 0 (cm3)
    4. 'root carbon cost of exudates'(g) initial value = 0 0 0
    5. 'root dry weight'(g)
    6. 'root length'(cm)
    7. 'root respiration'(g) initial value = 0 0 0
    8. 'root secondary potential carbon sink for growth'(g) initial value = 0 0 0
    9. 'root surface area'(cm2)
    10. 'root system carbon cost of exudates'(g) initial value = 0 0 0
    11. 'root system dry weight'(g)
    12. 'root system length'(cm)
    13. 'root system longitudinal growth'(cm) initial value = 0 0 0
    14. 'root system potential carbon sink for growth'(g) initial value = 0 0 0
    15. 'root system potential carbon sink for growth;major axis'(g) initial value = 0 0 0
        1. 'included root classes' = hypocotyl, primaryRoot, seminal, nodalroots, nodalroots1, nodalroots2, nodalroots3, nodalroots4, nodalroots5, braceroots, braceroots1, braceroots2, braceroots3, basalWhorl1, basalWhorl2, basalWhorl3, basalWhorl4 (noUnit)
    16. 'root system respiration'(g) initial value = 0 0 0
    17. 'root system secondary potential carbon sink for growth'(g) initial value = 0 0 0
    18. 'root system surface area'(cm2)
    19. 'root system volume'(cm3)
    20. 'root volume'(cm3)
4. 'plant template'(noUnit)
    1. 'carbon allocation to roots'(g) initial value = 0 0 0
    2. 'carbon allocation to shoot'(g) initial value = 0 0 0
    3. 'carbon available for growth'(g) initial value = 0 0 0
    4. 'carbon reserves'(g) initial value = 0 0 0
    5. 'carbon to dry weight ratio'(100%)
    6. 'plant carbon balance'(g)
    7. 'plant carbon income'(g) initial value = 0 0 0
    8. 'plant dry weight'(g)
    9. 'plant potential carbon sink for growth'(g) initial value = 0 0 0
    10. 'plant respiration'(g) initial value = 0 0 0
    11. 'relative carbon allocation to roots'(100%)
    12. 'relative carbon allocation to shoot'(100%)
    13. 'reserves'(g) initial value = 0 0 0
    14. 'root carbon cost of biological nitrogen fixation'(g) initial value = 0 0 0
    15. 'root carbon cost of exudates'(g) initial value = 0 0 0
    16. 'root carbon cost of nutrient uptake'(g) initial value = 0 0 0
    17. 'root carbon costs'(g) initial value = 0 0 0
        1. 'paths' = rootCarbonCostOfExudates;rootCarbonCostOfNutrientUptake;rootCarbonCostOfBiologicalNitrogenFixation (noUnit)
    18. 'root dry weight'(g)
    19. 'root growth scaling factor'(100%) initial value = 0 0 0
    20. 'root growth scaling factor;major axis'(100%) initial value = 0 0 0
    21. 'root length'(cm)
    22. 'root longitudinal growth'(cm) initial value = 0 0 0
    23. 'root potential carbon sink for growth'(g) initial value = 0 0 0
    24. 'root potential carbon sink for growth;major axis'(g) initial value = 0 0 0
    25. 'root respiration'(g) initial value = 0 0 0
    26. 'root secondary potential carbon sink for growth'(g) initial value = 0 0 0
    27. 'root surface area'(cm2)
    28. 'root volume'(cm3)
    29. 'secondary root growth scaling factor'(100%) initial value = 0 0 0
    30. 'seed carbohydrate content'(100%)
    31. 'seed carbohydrate to CFactor'(100%)
    32. 'shoot dry weight'(g)
    33. 'shoot potential carbon sink for growth'(g) initial value = 0 0 0
    34. 'shoot respiration'(g) initial value = 0 0 0
    35. 'stress factor'(noUnit)
    36. 'stress factor:impact on:leaf area expantion rate'(noUnit)
    37. 'stress factor:impact on:leaf respiration'(noUnit)
    38. 'stress factor:impact on:photosynthesis'(noUnit)

    39. 'stress factor:impact on:root potential longitudinal growth'(noUnit)
    40. 'stress factor:impact on:root segment carbon cost of exudates'(noUnit)
    41. 'stress factor:impact on:root segment respiration'(noUnit)
    42. 'stress factor:impact on:root segment secondary growth'(noUnit)
    43. 'stress factor:impact on:stem respiration'(noUnit)
5. 'plants'(noUnit)
    1. 'maize'(noUnit)
        1. 'carbon allocation to roots'(g) initial value = 0 0 8.35422e-05
        2. 'carbon allocation to shoot'(g) initial value = 0 0 0
        3. 'carbon available for growth'(g) initial value = 0 0 8.35422e-05
        4. 'carbon reserves'(g) initial value = 0 0 0
        5. 'carbon to dry weight ratio'(100%)
        6. 'plant carbon balance'(g)
        7. 'plant carbon income'(g) initial value = 0 0 8.35422e-05
        8. 'plant dry weight'(g)
        9. 'plant position' = 0 -2 0 (cm)
            1. 'hypocotyl' = 0 0 0 (cm)
                1. 'branches'(noUnit)
                    1. 'braceroots'(noUnit)
                    2. 'braceroots2'(noUnit)
                    3. 'nodalroots'(noUnit)
                    4. 'nodalroots2'(noUnit)
                    5. 'nodalroots3'(noUnit)
                    6. 'nodalroots4'(noUnit)
                2. 'data points'(noUnit)
                    1. 'data point00000' = 0 0 0 (cm)
                        1. 'aerenchyma formation'
                        2. 'branching frequency multiplier'(noUnit)
                        3. 'combined root class ID'(noUnit)
                        4. 'parent root class ID'(noUnit)
                        5. 'root circumference'(cm)
                        6. 'root class ID'(noUnit)
                        7. 'root diameter'(cm) initial value = 0 0.091 0
                        8. 'root hair density'
                        9. 'root hair diameter'
                        10. 'root hair length'
                        11. 'root hair surface area'(cm2/cm)
                        12. 'root length to base'(cm)
                        13. 'root potential secondary growth'(cm) initial value = 0 0.091 0
                        14. 'root segment age'(day)
                        15. 'root segment carbon cost of exudates'(g) initial value = 0 0 0
                        16. 'root segment dry weight'(g)
                        17. 'root segment length'(cm)
                        18. 'root segment length duration'(cm.day) initial value = 0 0 0
                        19. 'root segment respiration'(g/day)
                        20. 'root segment secondary potential carbon sink for growth'(g) initial value = 0 0 0
                        21. 'root segment surface area'(cm2)
                        22. 'root segment volume'(cm3)
                        23. 'spatial root density'(cm/cm3)
                3. 'growth rate multiplier' = 1 (noUnit)
                4. 'growthpoint'(cm) initial position = 0 0 0 0 0 0 0
                    1. 'branching frequency multiplier'(noUnit)
                    2. 'root circumference'(cm)
                    3. 'root diameter'(cm)
                    4. 'root longitudinal growth'(cm) initial value = 0 0 0
                    5. 'root potential longitudinal growth'(cm) initial value = 0 0 0
                    6. 'root potential secondary growth'(cm)
                    7. 'root segment age' = 0 (day)
                    8. 'root segment carbon cost of exudates' = 0 (g)
                    9. 'root segment dry weight' = 0 (g)
                    10. 'root segment length' = 0 (cm)
                    11. 'root segment length duration' = 0 (cm.day)
                    12. 'root segment potential carbon sink for growth'(g) initial value = 0 0 0

                13. 'root segment respiration' = 0 (g)
                14. 'root segment secondary potential carbon sink for growth' = 0 (g)
                15. 'root segment surface area' = 0 (cm2)
                16. 'root segment volume' = 0 (cm3)
            5. 'root carbon cost of exudates'(g) initial value = 0 0 0
            6. 'root dry weight'(g)
            7. 'root length'(cm)
            8. 'root respiration'(g) initial value = 0 0 0
            9. 'root secondary potential carbon sink for growth'(g) initial value = 0 0 0
            10. 'root surface area'(cm2)
            11. 'root system carbon cost of exudates'(g) initial value = 0 0 0
            12. 'root system dry weight'(g)
            13. 'root system length'(cm)
            14. 'root system longitudinal growth'(cm) initial value = 0 0 0
            15. 'root system potential carbon sink for growth'(g) initial value = 0 0 0
            16. 'root system potential carbon sink for growth;major axis'(g) initial value = 0 0 0
                1. 'included root classes' = hypocotyl, primaryRoot, seminal, nodalroots, nodalroots1, nodalroots2, nodalroots3, nodalroots4, nodalroots5, braceroots, braceroots1, braceroots2, braceroots3, basalWhorl1, basalWhorl2, basalWhorl3, basalWhorl4 (noUnit)
            17. 'root system respiration'(g) initial value = 0 0 0
            18. 'root system secondary potential carbon sink for growth'(g) initial value = 0 0 0
            19. 'root system surface area'(cm2)
            20. 'root system volume'(cm3)
            21. 'root type' = hypocotyl (noUnit)
            22. 'root volume'(cm3)
        2. 'primary root' = 0 0 0 (cm)
            1. 'branches'(noUnit)
                1. 'lateral'(noUnit)
                2. 'seminal'(noUnit)
            2. 'data points'(noUnit)
                1. 'data point00000' = 0 0 0 (cm)
                    1. 'aerenchyma formation'
                    2. 'branching frequency multiplier'(noUnit)
                    3. 'combined root class ID'(noUnit)
                    4. 'parent root class ID'(noUnit)
                    5. 'root circumference'(cm)
                    6. 'root class ID'(noUnit)
                    7. 'root diameter'(cm) initial value = 0 0.043 0
                    8. 'root hair density'
                    9. 'root hair diameter'
                    10. 'root hair length'
                    11. 'root hair surface area'(cm2/cm)
                    12. 'root length to base'(cm)
                    13. 'root potential secondary growth'(cm) initial value = 0 0.043 0
                    14. 'root segment age'(day)
                    15. 'root segment carbon cost of exudates'(g) initial value = 0 0 0
                    16. 'root segment dry weight'(g)
                    17. 'root segment length'(cm)
                    18. 'root segment length duration'(cm.day) initial value = 0 0 0
                    19. 'root segment respiration'(g/day)
                    20. 'root segment secondary potential carbon sink for growth'(g) initial value = 0 0 0
                    21. 'root segment surface area'(cm2)
                    22. 'root segment volume'(cm3)
                    23. 'spatial root density'(cm/cm3)
            3. 'growth rate multiplier' = 1 (noUnit)
            4. 'growthpoint'(cm) initial position = 0 0 0 0 0 0 0
                1. 'branching frequency multiplier'(noUnit)
                2. 'gravitropism'
                    1. 'multiplier'(noUnit)
                3. 'root circumference'(cm)
                4. 'root diameter'(cm)
                5. 'root longitudinal growth'(cm) initial value = 0 0 1.36
                6. 'root potential longitudinal growth'(cm) initial value = 0 0 1.36

                1. 'rate multiplier'(noUnit)
            7. 'root potential secondary growth'(cm)
            8. 'root segment age' = 0 (day)
            9. 'root segment carbon cost of exudates' = 0 (g)
            10. 'root segment dry weight' = 0 (g)
            11. 'root segment length' = 0 (cm)
            12. 'root segment length duration' = 0 (cm.day)
            13. 'root segment potential carbon sink for growth'(g) initial value = 0 0 8.35422e-05
            14. 'root segment respiration' = 0 (g)
            15. 'root segment secondary potential carbon sink for growth' = 0 (g)
            16. 'root segment specific weight' = 0 (g/cm3)
            17. 'root segment surface area' = 0 (cm2)
            18. 'root segment volume' = 0 (cm3)
        5. 'root carbon cost of exudates'(g) initial value = 0 0 0
        6. 'root dry weight'(g)
        7. 'root length'(cm)
        8. 'root respiration'(g) initial value = 0 0 0
        9. 'root secondary potential carbon sink for growth'(g) initial value = 0 0 0
        10. 'root surface area'(cm2)
        11. 'root system carbon cost of exudates'(g) initial value = 0 0 0
        12. 'root system dry weight'(g)
        13. 'root system length'(cm)
        14. 'root system longitudinal growth'(cm) initial value = 0 0 1.36
        15. 'root system potential carbon sink for growth'(g) initial value = 0 0 8.35422e-05
        16. 'root system potential carbon sink for growth;major axis'(g) initial value = 0 0 8.35422e-05
            1. 'included root classes' = hypocotyl, primaryRoot, seminal, nodalroots, nodalroots1, nodalroots2, nodalroots3, nodalroots4, nodalroots5, braceroots, braceroots1, braceroots2, braceroots3, basalWhorl1, basalWhorl2, basalWhorl3, basalWhorl4 (noUnit)
        17. 'root system respiration'(g) initial value = 0 0 0
        18. 'root system secondary potential carbon sink for growth'(g) initial value = 0 0 0
        19. 'root system surface area'(cm2)
        20. 'root system volume'(cm3)
        21. 'root type' = primaryRoot (noUnit)
        22. 'root volume'(cm3)
    3. 'shoot' = 0 0 0 (cm)
        1. 'area per plant'
        2. 'carbon allocation to leafs'(g) initial value = 0 0 0
        3. 'carbon allocation to stems'(g) initial value = 0 0 0
        4. 'extinction coefficient'
        5. 'leaf area'(cm2) initial value = 0 0 0
        6. 'leaf area index'(cm2/cm2)
        7. 'leaf area reduction coefficient'(cm2/cm2)
        8. 'leaf dry weight'(g) initial value = 0 0 0
        9. 'leaf potential carbon sink for growth'(g) initial value = 0 0 0
        10. 'leaf respiration'(g) initial value = 0 0 0
        11. 'light interception'(umol/cm2/day)
        12. 'photosynthesis'(g) initial value = 0 0 0
        13. 'potential leaf area'(cm2) initial value = 0 0 0
        14. 'relative carbon allocation to leafs'(100%)
        15. 'relative carbon allocation to stems'(100%)
        16. 'stem dry weight'(g) initial value = 0 0 0
        17. 'stem potential carbon sink for growth'(g) initial value = 0 0 0
        18. 'stem respiration'(g) initial value = 0 0 0
        19. 'stress adjusted potential leaf area'(cm2) initial value = 0 0 0
10. 'plant potential carbon sink for growth'(g) initial value = 0 0 8.35422e-05
11. 'plant respiration'(g) initial value = 0 0 0
12. 'plant type' = Ames21803 (noUnit)
13. 'planting time' = 0 (day)
14. 'relative carbon allocation to roots'(100%)
15. 'relative carbon allocation to shoot'(100%)
16. 'reserves'(g) initial value = 0 0.015 -0.000355552
17. 'root carbon cost of biological nitrogen fixation'(g) initial value = 0 0 0
18. 'root carbon cost of exudates'(g) initial value = 0 0 0

            19. 'root carbon cost of nutrient uptake'(g) initial value = 0 0 0
            20. 'root carbon costs'(g) initial value = 0 0 0
                1. 'paths' = rootCarbonCostOfExudates;rootCarbonCostOfNutrientUptake;rootCarbonCostOfBiologicalNitrogenFixation (noUnit)
            21. 'root dry weight'(g)
            22. 'root growth scaling factor'(100%) initial value = 0 1 0
            23. 'root growth scaling factor;major axis'(100%) initial value = 0 1 0
            24. 'root length'(cm)
            25. 'root longitudinal growth'(cm) initial value = 0 0 1.36
            26. 'root potential carbon sink for growth'(g) initial value = 0 0 8.35422e-05
            27. 'root potential carbon sink for growth;major axis'(g) initial value = 0 0 8.35422e-05
            28. 'root respiration'(g) initial value = 0 0 0
            29. 'root secondary potential carbon sink for growth'(g) initial value = 0 0 0
            30. 'root surface area'(cm2)
            31. 'root volume'(cm3)
            32. 'secondary root growth scaling factor'(100%) initial value = 0 1 0
            33. 'seed carbohydrate content'(100%)
            34. 'seed carbohydrate to CFactor'(100%)
            35. 'shoot dry weight'(g)
            36. 'shoot potential carbon sink for growth'(g) initial value = 0 0 0
            37. 'shoot respiration'(g) initial value = 0 0 0
            38. 'stress factor'(noUnit)
            39. 'stress factor:impact on:leaf area expantion rate'(noUnit)
            40. 'stress factor:impact on:leaf respiration'(noUnit)
            41. 'stress factor:impact on:photosynthesis'(noUnit)
            42. 'stress factor:impact on:root potential longitudinal growth'(noUnit)
            43. 'stress factor:impact on:root segment carbon cost of exudates'(noUnit)
            44. 'stress factor:impact on:root segment respiration'(noUnit)
            45. 'stress factor:impact on:root segment secondary growth'(noUnit)
            46. 'stress factor:impact on:stem respiration'(noUnit)
6. 'root type parameters'(noUnit)
    1. 'Ames to 1803'(noUnit)
        1. 'braceroots'(noUnit)
            1. 'aerenchyma formation' x,y pairs :{ 0 0 3 0 5 0.1 10 0.25 20 0.393 1000 0.393 }
            2. 'bottom boundary' = 0 (noUnit)
            3. 'bounce of the side' = 0 (noUnit)
            4. 'branch list'(noUnit)
                1. 'lateral of crown roots'(noUnit)
                    1. 'allow branches to form above ground' = 0 (noUnit)
                    2. 'branching frequency'(cm)=f{'uniform distribution'} minimum=0.100000 maximum=0.300000
                    3. 'branching spatial offset' = 12 (cm)
                    4. 'length root tip' = 10.93 (cm)
                    5. 'number of branches/whorl' = 1 (#)
            5. 'branching angle' = 140 (degrees)
            6. 'cannotgrowup' = 1 (noUnit)
            7. 'copy defaults from' = ../defaultsMajorAxis (noUnit)
            8. 'density' = 0.094 (g/cm3)
            9. 'diameter' x,y pairs :{ 0 0.4 8 0.4 15 0.15 24 0.1 100 0.1 }
            10. 'gravitropism.v2'(cm)=f{'uniform distribution'} minimum=-0.010000 maximum=-0.005000
            11. 'growth rate' x,y pairs :{ 0 0.01 5 1 10 4.5 17 4.5 22 0 1000 0 }
            12. 'length multiplier to diameter multiplier' x,y pairs :{ 0 0.25 1 1 2 1.5 3 1.8 4 2 100 2 1000 2 }
            13. 'length root tip without xylem vessels' = 2 (cm)
            14. 'local resource responses'(noUnit)
                1. 'impact on:branching frequency'(noUnit)
                    1. 'aggregation function' = maxRelativeDeviationFromOne (noUnit)
                    2. 'impact by:nitrate' x,y pairs :{ 0 1 2000 1 }
                    3. 'impact by:phosphorus' x,y pairs :{ 0 0.2 0.015 1 1000 1 }
                    4. 'impact by:potassium' x,y pairs :{ 0 1 1000 1 }
                2. 'impact on:gravitropism'(noUnit)
                    1. 'aggregation function' = maxRelativeDeviationFromOne (noUnit)
                    2. 'impact by:nitrate' x,y pairs :{ 0 1.5 100 1 2000 1 }
                    3. 'impact by:phosphorus' x,y pairs :{ 0 0.5 0.015 2 1000 0.5 }

            4. 'impact by:potassium' x,y pairs :{ 0 1 1000 1 }
        3. 'impact on:root potential longitudinal growth'(noUnit)
            1. 'aggregation function' = maxRelativeDeviationFromOne (noUnit)
            2. 'impact by:nitrate' x,y pairs :{ 0 1 2000 1 }
            3. 'impact by:phosphorus' x,y pairs :{ 0 0.2 0.015 1 1000 1 }
            4. 'impact by:potassium' x,y pairs :{ 0 1 1000 1 }
    15. 'longitudinal growth rate multiplier'(cm)=f{'uniform distribution'} minimum=0.700000 maximum=1.000000
    16. 'nitrate'(noUnit)
        1. 'Cmin' = 0.001 (umol/ml)
        2. 'Imax' x,y pairs :{ 0 1.21 2 2.1 40 2.1 }
        3. 'Km' x,y pairs :{ 0 0.0157 2 0.0522 40 0.0522 }
        4. 'minimal nutrient concentration' = 600 (umol/g)
        5. 'optimal nutrient concentration' = 1200 (umol/g)
    17. 'number of xylem poles' = 40 (noUnit)
    18. 'phosphorus'(noUnit)
        1. 'Cmin' = 0.0002 (umol/ml)
        2. 'Efflux' = 1e-06 (umol/cm/day)
        3. 'Imax' = 0.0555 (umol/cm2/day)
        4. 'Km' = 0.00545 (umol/ml)
        5. 'minimal nutrient concentration' = 30 (umol/g)
        6. 'optimal nutrient concentration' = 60 (umol/g)
    19. 'potassium'(noUnit)
        1. 'Cmin' = 0.002 (umol/ml)
        2. 'Efflux' = 1e-06 (umol/cm/day)
        3. 'Imax' = 0.467 (umol/cm2/day)
        4. 'Km' = 0.014 (umol/ml)
        5. 'minimal nutrient concentration' = 117 (umol/g)
        6. 'optimal nutrient concentration' = 234 (umol/g)
    20. 'radial hydraulic conductivity' x,y pairs :{ 0 0 1 0.000216 10 0.000216 20 0.00025 30 0.000216 40 0.0001 60 0 }
    21. 'reduction in respiration due to aerenchyma' x,y pairs :{ 0 0 0.3 0.7 0.6 1 }
    22. 'regular topology' = 4 (noUnit)
    23. 'relative carbon cost of exudation' x,y pairs :{ 0 5e-06 100 5e-06 }
    24. 'relative respiration' x,y pairs :{ 0 0.09 2 0.04 6 0.04 1000 0.04 }
    25. 'root class ID' = 102 (noUnit)
    26. 'root hair density' x,y pairs :{ 0 2000 1 2000 2 2000 10 2000 30 0 2000 0 }
    27. 'root hair diameter' = 0.0005 (cm)
    28. 'root hair length' x,y pairs :{ 0 0 1 0 2 0.028 2000 0.028 }
    29. 'soil impedence.v2'(cm)=f{'uniform distribution'} minimum=-0.030000 maximum=0.030000
    30. 'top boundary' = 1 (noUnit)
2. 'braceroots2'(noUnit)
    1. 'aerenchyma formation' x,y pairs :{ 0 0 3 0 5 0.1 10 0.25 20 0.393 1000 0.393 }
    2. 'bottom boundary' = 0 (noUnit)
    3. 'bounce of the side' = 0 (noUnit)
    4. 'branch list'(noUnit)
        1. 'lateral of crown roots'(noUnit)
            1. 'allow branches to form above ground' = 0 (noUnit)
            2. 'branching frequency'(cm)=f{'uniform distribution'} minimum=0.100000 maximum=0.400000
            3. 'branching spatial offset' = 15 (cm)
            4. 'length root tip' = 10.93 (cm)
            5. 'number of branches/whorl' = 1 (#)
    5. 'branching angle' = 130 (degrees)
    6. 'cannotgrowup' = 1 (noUnit)
    7. 'copy defaults from' = ../defaultsMajorAxis (noUnit)
    8. 'density' = 0.094 (g/cm3)
    9. 'diameter' x,y pairs :{ 0 0.5 9 0.5 16 0.2 24 0.1 100 0.1 }
    10. 'gravitropism.v2'(cm)=f{'uniform distribution'} minimum=-0.010000 maximum=-0.005000
    11. 'growth rate' x,y pairs :{ 0 0.01 5 1 10 4.5 17 4.5 22 0 1000 0 }
    12. 'length multiplier to diameter multiplier' x,y pairs :{ 0 0.25 1 1 2 1.5 3 1.8 4 2 100 2 1000 2 }
    13. 'length root tip without xylem vessels' = 2 (cm)
    14. 'local resource responses'(noUnit)
        1. 'impact on:branching frequency'(noUnit)

            1. 'aggregation function' = maxRelativeDeviationFromOne (noUnit)
            2. 'impact by:nitrate' x,y pairs :{ 0 1 2000 1 }
            3. 'impact by:phosphorus' x,y pairs :{ 0 0.2 0.015 1 1000 1 }
            4. 'impact by:potassium' x,y pairs :{ 0 1 1000 1 }
        2. 'impact on:gravitropism'(noUnit)
            1. 'aggregation function' = maxRelativeDeviationFromOne (noUnit)
            2. 'impact by:nitrate' x,y pairs :{ 0 1.5 100 1 2000 1 }
            3. 'impact by:phosphorus' x,y pairs :{ 0 0.5 0.015 2 1000 0.5 }
            4. 'impact by:potassium' x,y pairs :{ 0 1 1000 1 }
        3. 'impact on:root potential longitudinal growth'(noUnit)
            1. 'aggregation function' = maxRelativeDeviationFromOne (noUnit)
            2. 'impact by:nitrate' x,y pairs :{ 0 1 2000 1 }
            3. 'impact by:phosphorus' x,y pairs :{ 0 0.2 0.015 1 1000 1 }
            4. 'impact by:potassium' x,y pairs :{ 0 1 1000 1 }
    15. 'longitudinal growth rate multiplier'(cm)=f{'uniform distribution'} minimum=0.700000 maximum=1.000000
    16. 'nitrate'(noUnit)
        1. 'Cmin' = 0.001 (umol/ml)
        2. 'Imax' x,y pairs :{ 0 1.21 2 2.1 40 2.1 }
        3. 'Km' x,y pairs :{ 0 0.0157 2 0.0522 40 0.0522 }
        4. 'minimal nutrient concentration' = 600 (umol/g)
        5. 'optimal nutrient concentration' = 1200 (umol/g)
    17. 'number of xylem poles' = 48 (noUnit)
    18. 'phosphorus'(noUnit)
        1. 'Cmin' = 0.0002 (umol/ml)
        2. 'Efflux' = 1e-06 (umol/cm/day)
        3. 'Imax' = 0.0555 (umol/cm2/day)
        4. 'Km' = 0.00545 (umol/ml)
        5. 'minimal nutrient concentration' = 30 (umol/g)
        6. 'optimal nutrient concentration' = 60 (umol/g)
    19. 'potassium'(noUnit)
        1. 'Cmin' = 0.002 (umol/ml)
        2. 'Efflux' = 1e-06 (umol/cm/day)
        3. 'Imax' = 0.467 (umol/cm2/day)
        4. 'Km' = 0.014 (umol/ml)
        5. 'minimal nutrient concentration' = 117 (umol/g)
        6. 'optimal nutrient concentration' = 234 (umol/g)
    20. 'radial hydraulic conductivity' x,y pairs :{ 0 0 1 0.000216 10 0.000216 20 0.00025 30 0.000216 40 0.0001 60 0 }
    21. 'reduction in respiration due to aerenchyma' x,y pairs :{ 0 0 0.3 0.7 0.6 1 }
    22. 'regular topology' = 3 (noUnit)
    23. 'relative carbon cost of exudation' x,y pairs :{ 0 5e-06 100 5e-06 }
    24. 'relative respiration' x,y pairs :{ 0 0.09 2 0.04 6 0.04 1000 0.04 }
    25. 'root class ID' = 102 (noUnit)
    26. 'root hair density' x,y pairs :{ 0 2000 1 2000 2 2000 10 2000 30 0 2000 0 }
    27. 'root hair diameter' = 0.0005 (cm)
    28. 'root hair length' x,y pairs :{ 0 0 1 0 2 0.028 2000 0.028 }
    29. 'soil impedence.v2'(cm)=f{'uniform distribution'} minimum=-0.030000 maximum=0.030000
    30. 'top boundary' = 1 (noUnit)
3. 'defaults'(noUnit)
    1. 'bottom boundary' = 0 (noUnit)
    2. 'bounce of the side' = 0 (noUnit)
    3. 'cannotgrowup' = 1 (noUnit)
    4. 'local resource responses'(noUnit)
        1. 'impact on:branching frequency'(noUnit)
            1. 'aggregation function' = maxRelativeDeviationFromOne (noUnit)
            2. 'impact by:nitrate' x,y pairs :{ 0 1 2000 1 }
            3. 'impact by:phosphorus' x,y pairs :{ 0 0.5 0.015 1 1000 1 }
            4. 'impact by:potassium' x,y pairs :{ 0 1 1000 1 }
        2. 'impact on:gravitropism'(noUnit)
            1. 'aggregation function' = maxRelativeDeviationFromOne (noUnit)
            2. 'impact by:nitrate' x,y pairs :{ 0 1 2000 1 }
            3. 'impact by:phosphorus' x,y pairs :{ 0 1 1000 1 }

            4. 'impact by:potassium' x,y pairs :{ 0 1 1000 1 }
        3. 'impact on:root potential longitudinal growth'(noUnit)
            1. 'aggregation function' = maxRelativeDeviationFromOne (noUnit)
            2. 'impact by:nitrate' x,y pairs :{ -10 1.5 50 1.5 100 1 2000 1 }
            3. 'impact by:phosphorus' x,y pairs :{ 0 0.2 0.015 1 1000 1 }
            4. 'impact by:potassium' x,y pairs :{ 0 1 1000 1 }
    5. 'phosphorus'(noUnit)
        1. 'Cmin' = 0.0002 (umol/ml)
        2. 'Efflux' = 1e-06 (umol/cm/day)
        3. 'Imax' = 0.0555 (umol/cm2/day)
        4. 'Km' = 0.00545 (umol/ml)
        5. 'minimal nutrient concentration' = 30 (umol/g)
        6. 'optimal nutrient concentration' = 60 (umol/g)
    6. 'potassium'(noUnit)
        1. 'Cmin' = 0.002 (umol/ml)
        2. 'Efflux' = 1e-06 (umol/cm/day)
        3. 'Imax' = 0.467 (umol/cm2/day)
        4. 'Km' = 0.014 (umol/ml)
        5. 'minimal nutrient concentration' = 117 (umol/g)
        6. 'optimal nutrient concentration' = 234 (umol/g)
    7. 'relative carbon cost of exudation' x,y pairs :{ 0 5e-06 100 5e-06 }
    8. 'relative respiration' x,y pairs :{ 0 0.09 2 0.04 6 0.04 1000 0.04 }
    9. 'top boundary' = 1 (noUnit)
4. 'defaults major axis'(noUnit)
    1. 'bottom boundary' = 0 (noUnit)
    2. 'bounce of the side' = 0 (noUnit)
    3. 'cannotgrowup' = 1 (noUnit)
    4. 'copy defaults from' = ../defaults (noUnit)
    5. 'local resource responses'(noUnit)
        1. 'impact on:branching frequency'(noUnit)
            1. 'aggregation function' = maxRelativeDeviationFromOne (noUnit)
            2. 'impact by:nitrate' x,y pairs :{ 0 1 2000 1 }
            3. 'impact by:phosphorus' x,y pairs :{ 0 0.2 0.015 1 1000 1 }
            4. 'impact by:potassium' x,y pairs :{ 0 1 1000 1 }
        2. 'impact on:gravitropism'(noUnit)
            1. 'aggregation function' = maxRelativeDeviationFromOne (noUnit)
            2. 'impact by:nitrate' x,y pairs :{ 0 1.5 100 1 2000 1 }
            3. 'impact by:phosphorus' x,y pairs :{ 0 0.5 0.015 2 1000 0.5 }
            4. 'impact by:potassium' x,y pairs :{ 0 1 1000 1 }
        3. 'impact on:root potential longitudinal growth'(noUnit)
            1. 'aggregation function' = maxRelativeDeviationFromOne (noUnit)
            2. 'impact by:nitrate' x,y pairs :{ 0 1 2000 1 }
            3. 'impact by:phosphorus' x,y pairs :{ 0 0.2 0.015 1 1000 1 }
            4. 'impact by:potassium' x,y pairs :{ 0 1 1000 1 }
    6. 'phosphorus'(noUnit)
        1. 'Cmin' = 0.0002 (umol/ml)
        2. 'Efflux' = 1e-06 (umol/cm/day)
        3. 'Imax' = 0.0555 (umol/cm2/day)
        4. 'Km' = 0.00545 (umol/ml)
        5. 'minimal nutrient concentration' = 30 (umol/g)
        6. 'optimal nutrient concentration' = 60 (umol/g)
    7. 'potassium'(noUnit)
        1. 'Cmin' = 0.002 (umol/ml)
        2. 'Efflux' = 1e-06 (umol/cm/day)
        3. 'Imax' = 0.467 (umol/cm2/day)
        4. 'Km' = 0.014 (umol/ml)
        5. 'minimal nutrient concentration' = 117 (umol/g)
        6. 'optimal nutrient concentration' = 234 (umol/g)
    8. 'relative carbon cost of exudation' x,y pairs :{ 0 5e-06 100 5e-06 }
    9. 'relative respiration' x,y pairs :{ 0 0.09 2 0.04 6 0.04 1000 0.04 }
    10. 'top boundary' = 1 (noUnit)
5. 'finelateral'(noUnit)
    1. 'aerenchyma formation' x,y pairs :{ 0 0 3 0 5 0.1 10 0.25 20 0.393 1000 0.393 }

    2. 'branch list'(noUnit)
        1. 'finelateral2'(noUnit)
            1. 'allow branches to form above ground' = 0 (noUnit)
            2. 'branching frequency'(cm)=f{'uniform distribution'} minimum=0.400000 maximum=0.600000
            3. 'length root tip' = 1.5 (cm)
    3. 'branching angle' = 62.83 (degrees)
    4. 'density' = 0.094 (g/cm3)
    5. 'diameter' = 0.025 (cm)
    6. 'gravitropism.v2' = 0 0 0 (cm)
    7. 'growth rate' x,y pairs :{ 0 0.01 1 0.35 6 0 1000 0 }
    8. 'length multiplier to diameter multiplier' x,y pairs :{ 0 0.25 1 1 2 1.5 3 1.8 4 2 100 2 1000 2 }
    9. 'length root tip without xylem vessels' = 2 (cm)
    10. 'longitudinal growth rate multiplier'(cm)=f{'normal distribution'} minimum=0.500000 maximum=1.500000 mean=1.000000 stdev=0.100000
    11. 'nitrate'(noUnit)
        1. 'Cmin' = 0.0017 (umol/ml)
        2. 'Imax' = 1.27 (umol/cm2/day)
        3. 'Km' = 0.0027 (umol/ml)
        4. 'minimal nutrient concentration' = 600 (umol/g)
        5. 'optimal nutrient concentration' = 1200 (umol/g)
    12. 'number of xylem poles' = 4 (noUnit)
    13. 'radial hydraulic conductivity' x,y pairs :{ 0 0 1 0.000416 60 0.000416 }
    14. 'reduction in respiration due to aerenchyma' x,y pairs :{ 0 0 0.3 0.7 0.6 1 }
    15. 'relative carbon cost of exudation' x,y pairs :{ 0 5e-06 100 1e-06 }
    16. 'root class ID' = 98 (noUnit)
    17. 'root hair density' x,y pairs :{ 0 2000 1 2000 2 2000 10 2000 30 0 2000 0 }
    18. 'root hair diameter' = 0.0005 (cm)
    19. 'root hair length' x,y pairs :{ 0 0 1 0 2 0.028 2000 0.028 }
    20. 'soil impedence.v2'(cm)=f{'uniform distribution'} minimum=-0.050000 maximum=0.050000
6. 'finelateral2'(noUnit)
    1. 'aerenchyma formation' x,y pairs :{ 0 0 3 0 5 0.1 10 0.25 20 0.393 1000 0.393 }
    2. 'branch list'(noUnit)
    3. 'branching angle' = 62.83 (degrees)
    4. 'density' = 0.094 (g/cm3)
    5. 'diameter' = 0.015 (cm)
    6. 'gravitropism.v2' = 0 0 0 (cm)
    7. 'growth rate' x,y pairs :{ 0 0.001 1 0.28 4 0 1000 0 }
    8. 'length multiplier to diameter multiplier' x,y pairs :{ 0 0.25 1 1 2 1.5 3 1.8 4 2 100 2 1000 2 }
    9. 'length root tip without xylem vessels' = 2 (cm)
    10. 'longitudinal growth rate multiplier'(cm)=f{'normal distribution'} minimum=0.500000 maximum=1.500000 mean=1.000000 stdev=0.100000
    11. 'nitrate'(noUnit)
        1. 'Cmin' = 0.0017 (umol/ml)
        2. 'Imax' = 1.27 (umol/cm2/day)
        3. 'Km' = 0.0027 (umol/ml)
        4. 'minimal nutrient concentration' = 600 (umol/g)
        5. 'optimal nutrient concentration' = 1200 (umol/g)
    12. 'number of xylem poles' = 4 (noUnit)
    13. 'radial hydraulic conductivity' x,y pairs :{ 0 0 1 0.000416 60 0.000416 }
    14. 'reduction in respiration due to aerenchyma' x,y pairs :{ 0 0 0.3 0.7 0.6 1 }
    15. 'relative carbon cost of exudation' x,y pairs :{ 0 5e-06 100 1e-06 }
    16. 'root class ID' = 98 (noUnit)
    17. 'root hair density' x,y pairs :{ 0 2000 1 2000 2 2000 10 2000 30 0 2000 0 }
    18. 'root hair diameter' = 0.0005 (cm)
    19. 'root hair length' x,y pairs :{ 0 0 1 0 2 0.028 2000 0.028 }
    20. 'soil impedence.v2'(cm)=f{'uniform distribution'} minimum=-0.050000 maximum=0.050000
7. 'hypocotyl'(noUnit)
    1. 'aerenchyma formation' x,y pairs :{ 0 0 100 0 }
    2. 'bottom boundary' = 0 (noUnit)
    3. 'bounce of the side' = 0 (noUnit)
    4. 'branch list'(noUnit)
        1. 'braceroots'(noUnit)
            1. 'allometric scaling' = 1 (noUnit)

        2. 'branching spatial offset' = 4 (cm)
        3. 'branching time offset' = 25 (day)
        4. 'max number of branches' = 14 (#)
        5. 'number of branches/whorl' = 14 (#)
    2. 'braceroots2'(noUnit)
        1. 'allometric scaling' = 1 (noUnit)
        2. 'branching delay' = 14 (day)
        3. 'branching frequency' = 5 (cm)
        4. 'branching spatial offset' = 7 (cm)
        5. 'branching time offset' = 36 (day)
        6. 'number of branches/whorl' = 20 (#)
    3. 'nodalroots'(noUnit)
        1. 'branching spatial offset' = 1.5 (cm)
        2. 'branching time offset' = 7 (day)
        3. 'max number of branches' = 3 (#)
        4. 'number of branches/whorl' = 3 (#)
    4. 'nodalroots2'(noUnit)
        1. 'allometric scaling' = 1 (noUnit)
        2. 'branching spatial offset' = 1.9 (cm)
        3. 'branching time offset' = 12 (day)
        4. 'max number of branches' = 3 (#)
        5. 'number of branches/whorl' = 3 (#)
    5. 'nodalroots3'(noUnit)
        1. 'allometric scaling' = 1 (noUnit)
        2. 'branching spatial offset' = 2.1 (cm)
        3. 'branching time offset' = 17 (day)
        4. 'max number of branches' = 3 (#)
        5. 'number of branches/whorl' = 3 (#)
    6. 'nodalroots4'(noUnit)
        1. 'allometric scaling' = 1 (noUnit)
        2. 'branching spatial offset' = 2.3 (cm)
        3. 'branching time offset' = 23 (day)
        4. 'max number of branches' = 6 (#)
        5. 'number of branches/whorl' = 6 (#)
5. 'cannotgrowup' = 0 (noUnit)
6. 'copy defaults from' = ../defaultsMajorAxis (noUnit)
7. 'density' = 0.094 (g/cm3)
8. 'diameter' = 0.091 (cm)
9. 'gravitropism.v2' = 0 1 0 (cm)
10. 'growth rate' x,y pairs :{ 0 0 1 0 2 0.45 3 1.18 4 1.29 5 0 1000 0 }
11. 'length root tip without xylem vessels' = 2 (cm)
12. 'local resource responses'(noUnit)
    1. 'impact on:branching frequency'(noUnit)
        1. 'aggregation function' = maxRelativeDeviationFromOne (noUnit)
        2. 'impact by:nitrate' x,y pairs :{ 0 1 10000 1 }
        3. 'impact by:phosphorus' x,y pairs :{ 0 1 1000 1 }
        4. 'impact by:potassium' x,y pairs :{ 0 1 1000 1 }
    2. 'impact on:gravitropism'(noUnit)
        1. 'aggregation function' = maxRelativeDeviationFromOne (noUnit)
        2. 'impact by:nitrate' x,y pairs :{ 0 1 10000 1 }
        3. 'impact by:phosphorus' x,y pairs :{ 0 1 1000 1 }
        4. 'impact by:potassium' x,y pairs :{ 0 1 1000 1 }
    3. 'impact on:root potential longitudinal growth'(noUnit)
        1. 'aggregation function' = maxRelativeDeviationFromOne (noUnit)
        2. 'impact by:nitrate' x,y pairs :{ 0 1 10000 1 }
        3. 'impact by:phosphorus' x,y pairs :{ 0 1 1000 1 }
        4. 'impact by:potassium' x,y pairs :{ 0 1 1000 1 }
13. 'nitrate'(noUnit)
    1. 'Cmin' = 0 (umol/ml)
    2. 'Imax' = 0 (umol/cm2/day)
    3. 'Km' = 1 (umol/ml)
    4. 'minimal nutrient concentration' = 600 (umol/g)
    5. 'optimal nutrient concentration' = 1200 (umol/g)

        14. 'number of xylem poles' = 61 (noUnit)
        15. 'phosphorus'(noUnit)
            1. 'Cmin' = 0.0002 (umol/ml)
            2. 'Efflux' = 1e-06 (umol/cm/day)
            3. 'Imax' = 0.0555 (umol/cm2/day)
            4. 'Km' = 0.00545 (umol/ml)
            5. 'minimal nutrient concentration' = 30 (umol/g)
            6. 'optimal nutrient concentration' = 60 (umol/g)
        16. 'potassium'(noUnit)
            1. 'Cmin' = 0.002 (umol/ml)
            2. 'Efflux' = 1e-06 (umol/cm/day)
            3. 'Imax' = 0.467 (umol/cm2/day)
            4. 'Km' = 0.014 (umol/ml)
            5. 'minimal nutrient concentration' = 117 (umol/g)
            6. 'optimal nutrient concentration' = 234 (umol/g)
        17. 'radial hydraulic conductivity' x,y pairs :{ 0 0 60 0 }
        18. 'reduction in respiration due to aerenchyma' x,y pairs :{ 0 0 0.3 0.7 0.6 1 }
        19. 'relative carbon cost of exudation' x,y pairs :{ 0 0 100 0 }
        20. 'relative respiration' x,y pairs :{ 0 0.09 2 0.04 6 0.04 1000 0.04 }
        21. 'root class ID' = 97 (noUnit)
        22. 'root hair density' x,y pairs :{ 0 0 2000 0 }
        23. 'root hair diameter' = 0.0005 (cm)
        24. 'root hair length' x,y pairs :{ 0 0 1 0 2 0.028 2000 0.028 }
        25. 'soil impedence.v2'(cm)=f{'uniform distribution'} minimum=-0.300000 maximum=0.300000
        26. 'top boundary' = 0 (noUnit)
    8. 'lateral'(noUnit)
        1. 'aerenchyma formation' x,y pairs :{ 0 0 3 0 5 0.1 10 0.25 20 0.393 1000 0.393 }
        2. 'bottom boundary' = 0 (noUnit)
        3. 'bounce of the side' = 0 (noUnit)
        4. 'branch list'(noUnit)
            1. 'finelateral'(noUnit)
                1. 'allow branches to form above ground' = 0 (noUnit)
                2. 'branching frequency'(cm)=f{'uniform distribution'} minimum=0.150000 maximum=0.350000
                3. 'length root tip' = 4 (cm)
        5. 'branching angle' = 90 (degrees)
        6. 'cannotgrowup' = 1 (noUnit)
        7. 'density' = 0.094 (g/cm3)
        8. 'diameter' = 0.04 (cm)
        9. 'gravitropism.v2' = 0 0 0 (cm)
        10. 'growth rate' x,y pairs :{ 0 0.01 1 0.2 3 0.4 7 1 11 0 1000 0 }
        11. 'length multiplier to diameter multiplier' x,y pairs :{ 0 0.25 1 1 2 1.5 3 1.8 4 2 100 2 1000 2 }
        12. 'length root tip without xylem vessels' = 2 (cm)
        13. 'local resource responses'(noUnit)
            1. 'impact on:branching frequency'(noUnit)
                1. 'aggregation function' = maxRelativeDeviationFromOne (noUnit)
                2. 'impact by:nitrate' x,y pairs :{ 0 1 2000 1 }
                3. 'impact by:phosphorus' x,y pairs :{ 0 0.5 0.015 1 1000 1 }
                4. 'impact by:potassium' x,y pairs :{ 0 1 1000 1 }
            2. 'impact on:gravitropism'(noUnit)
                1. 'aggregation function' = maxRelativeDeviationFromOne (noUnit)
                2. 'impact by:nitrate' x,y pairs :{ 0 1 2000 1 }
                3. 'impact by:phosphorus' x,y pairs :{ 0 1 1000 1 }
                4. 'impact by:potassium' x,y pairs :{ 0 1 1000 1 }
            3. 'impact on:root potential longitudinal growth'(noUnit)
                1. 'aggregation function' = maxRelativeDeviationFromOne (noUnit)
                2. 'impact by:nitrate' x,y pairs :{ -10 1.5 50 1.5 100 1 2000 1 }
                3. 'impact by:phosphorus' x,y pairs :{ 0 0.2 0.015 1 1000 1 }
                4. 'impact by:potassium' x,y pairs :{ 0 1 1000 1 }
        14. 'longitudinal growth rate multiplier'(cm)=f{'lognormal distribution'} minimum=0.100000 maximum=2.000000
        15. 'nitrate'(noUnit)
            1. 'Cmin' = 0.0017 (umol/ml)
            2. 'Imax' = 1.27 (umol/cm2/day)

        3. 'Km' = 0.0027 (umol/ml)
        4. 'minimal nutrient concentration' = 600 (umol/g)
        5. 'optimal nutrient concentration' = 1200 (umol/g)
    16. 'number of xylem poles' = 4 (noUnit)
    17. 'phosphorus'(noUnit)
        1. 'Cmin' = 0.0002 (umol/ml)
        2. 'Efflux' = 1e-06 (umol/cm/day)
        3. 'Imax' = 0.0555 (umol/cm2/day)
        4. 'Km' = 0.00545 (umol/ml)
        5. 'minimal nutrient concentration' = 30 (umol/g)
        6. 'optimal nutrient concentration' = 60 (umol/g)
    18. 'potassium'(noUnit)
        1. 'Cmin' = 0.002 (umol/ml)
        2. 'Efflux' = 1e-06 (umol/cm/day)
        3. 'Imax' = 0.467 (umol/cm2/day)
        4. 'Km' = 0.014 (umol/ml)
        5. 'minimal nutrient concentration' = 117 (umol/g)
        6. 'optimal nutrient concentration' = 234 (umol/g)
    19. 'radial hydraulic conductivity' x,y pairs :{ 0 0 1 0.000416 60 0.000416 }
    20. 'reduction in respiration due to aerenchyma' x,y pairs :{ 0 0 0.3 0.7 0.6 1 }
    21. 'relative carbon cost of exudation' x,y pairs :{ 0 5e-06 100 3e-06 }
    22. 'relative respiration' x,y pairs :{ 0 0.09 2 0.04 6 0.04 1000 0.04 }
    23. 'root class ID' = 98 (noUnit)
    24. 'root hair density' x,y pairs :{ 0 2000 1 2000 2 2000 10 2000 30 0 2000 0 }
    25. 'root hair diameter' = 0.0005 (cm)
    26. 'root hair length' x,y pairs :{ 0 0 1 0 2 0.028 2000 0.028 }
    27. 'soil impedence.v2'(cm)=f{'uniform distribution'} minimum=-0.100000 maximum=0.100000
    28. 'top boundary' = 1 (noUnit)
9. 'lateral of crown roots'(noUnit)
    1. 'aerenchyma formation' x,y pairs :{ 0 0 3 0 5 0.1 10 0.25 20 0.393 1000 0.393 }
    2. 'branch list'(noUnit)
        1. 'lateral'(noUnit)
            1. 'allow branches to form above ground' = 0 (noUnit)
            2. 'branching frequency'(cm)=f{'uniform distribution'} minimum=0.250000 maximum=0.350000
            3. 'length root tip' = 5 (cm)
    3. 'branching angle' = 90 (degrees)
    4. 'density' = 0.094 (g/cm3)
    5. 'diameter' = 0.07 (cm)
    6. 'gravitropism.v2' = 0 0 0 (cm)
    7. 'growth rate' x,y pairs :{ 0 0.1 1 0.5 3 1.2 12 1.2 18 0 1000 0 }
    8. 'length multiplier to diameter multiplier' x,y pairs :{ 0 0.25 1 1 2 1.5 3 1.8 4 2 100 2 1000 2 }
    9. 'length root tip without xylem vessels' = 2 (cm)
    10. 'longitudinal growth rate multiplier'(cm)=f{'normal distribution'} minimum=0.100000 maximum=1.000000 mean=0.400000 stdev=0.300000
    11. 'nitrate'(noUnit)
        1. 'Cmin' = 0.0017 (umol/ml)
        2. 'Imax' = 1.27 (umol/cm2/day)
        3. 'Km' = 0.0027 (umol/ml)
        4. 'minimal nutrient concentration' = 600 (umol/g)
        5. 'optimal nutrient concentration' = 1200 (umol/g)
    12. 'number of xylem poles' = 4 (noUnit)
    13. 'radial hydraulic conductivity' x,y pairs :{ 0 0 1 0.000216 60 0.000216 }
    14. 'reduction in respiration due to aerenchyma' x,y pairs :{ 0 0 0.3 0.7 0.6 1 }
    15. 'relative carbon cost of exudation' x,y pairs :{ 0 5e-06 100 4e-06 }
    16. 'root class ID' = 98 (noUnit)
    17. 'root hair density' x,y pairs :{ 0 2000 1 2000 2 2000 10 2000 30 0 2000 0 }
    18. 'root hair diameter' = 0.0005 (cm)
    19. 'root hair length' x,y pairs :{ 0 0 1 0 2 0.028 2000 0.028 }
    20. 'soil impedence.v2'(cm)=f{'uniform distribution'} minimum=-0.050000 maximum=0.050000
10. 'nodalroots'(noUnit)
    1. 'aerenchyma formation' x,y pairs :{ 0 0 3 0 5 0.1 10 0.25 20 0.393 1000 0.393 }
    2. 'bottom boundary' = 0 (noUnit)
    3. 'bounce of the side' = 0 (noUnit)

4. 'branch list'(noUnit)
    1. 'lateral'(noUnit)
        1. 'allow branches to form above ground' = 0 (noUnit)
        2. 'branching frequency'(cm)=f{'uniform distribution'} minimum=0.121000 maximum=0.321000
        3. 'length root tip' = 10.93 (cm)
5. 'branching angle' = 125.67 (degrees)
6. 'cannotgrowup' = 1 (noUnit)
7. 'copy defaults from' = ../defaultsMajorAxis (noUnit)
8. 'density' = 0.094 (g/cm3)
9. 'diameter' x,y pairs :{ 0 0.069 8 0.063 13 0.062 18 0.061 }
10. 'gravitropism.v2'(cm)=f{'uniform distribution'} minimum=-0.010000 maximum=-0.005000
11. 'growth rate' x,y pairs :{ 0 2.97 1 2.98 3 3 4 3 5 3.01 7 3.03 10 3.05 12 3.07 15 3.09 17 3.1 18 3.11 40 3.11 }
12. 'length multiplier to diameter multiplier' x,y pairs :{ 0 0.25 1 1 2 1.5 3 1.8 4 2 100 2 1000 2 }
13. 'length root tip without xylem vessels' = 2 (cm)
14. 'local resource responses'(noUnit)
    1. 'impact on:branching frequency'(noUnit)
        1. 'aggregation function' = maxRelativeDeviationFromOne (noUnit)
        2. 'impact by:nitrate' x,y pairs :{ 0 1 2000 1 }
        3. 'impact by:phosphorus' x,y pairs :{ 0 0.2 0.015 1 1000 1 }
        4. 'impact by:potassium' x,y pairs :{ 0 1 1000 1 }
    2. 'impact on:gravitropism'(noUnit)
        1. 'aggregation function' = maxRelativeDeviationFromOne (noUnit)
        2. 'impact by:nitrate' x,y pairs :{ 0 1.5 100 1 2000 1 }
        3. 'impact by:phosphorus' x,y pairs :{ 0 0.5 0.015 2 1000 0.5 }
        4. 'impact by:potassium' x,y pairs :{ 0 1 1000 1 }
    3. 'impact on:root potential longitudinal growth'(noUnit)
        1. 'aggregation function' = maxRelativeDeviationFromOne (noUnit)
        2. 'impact by:nitrate' x,y pairs :{ 0 1 2000 1 }
        3. 'impact by:phosphorus' x,y pairs :{ 0 0.2 0.015 1 1000 1 }
        4. 'impact by:potassium' x,y pairs :{ 0 1 1000 1 }
15. 'longitudinal growth rate multiplier'(cm)=f{'normal distribution'} minimum=0.600000 maximum=1.200000 mean=1.000000 stdev=0.100000
16. 'nitrate'(noUnit)
    1. 'Cmin' = 0.001 (umol/ml)
    2. 'Imax' x,y pairs :{ 0 1.21 2 2.1 40 2.1 }
    3. 'Km' x,y pairs :{ 0 0.0157 2 0.0522 40 0.0522 }
    4. 'minimal nutrient concentration' = 600 (umol/g)
    5. 'optimal nutrient concentration' = 1200 (umol/g)
17. 'number of xylem poles' = 10 (noUnit)
18. 'phosphorus'(noUnit)
    1. 'Cmin' = 0.0002 (umol/ml)
    2. 'Efflux' = 1e-06 (umol/cm/day)
    3. 'Imax' = 0.0555 (umol/cm2/day)
    4. 'Km' = 0.00545 (umol/ml)
    5. 'minimal nutrient concentration' = 30 (umol/g)
    6. 'optimal nutrient concentration' = 60 (umol/g)
19. 'potassium'(noUnit)
    1. 'Cmin' = 0.002 (umol/ml)
    2. 'Efflux' = 1e-06 (umol/cm/day)
    3. 'Imax' = 0.467 (umol/cm2/day)
    4. 'Km' = 0.014 (umol/ml)
    5. 'minimal nutrient concentration' = 117 (umol/g)
    6. 'optimal nutrient concentration' = 234 (umol/g)
20. 'radial hydraulic conductivity' x,y pairs :{ 0 0 1 0.000216 10 0.000216 20 0.00025 30 0.000216 40 0.0001 60 0 }
21. 'reduction in respiration due to aerenchyma' x,y pairs :{ 0 0 0.3 0.7 0.6 1 }
22. 'regular topology' = 3 (noUnit)
23. 'relative carbon cost of exudation' x,y pairs :{ 0 5e-06 100 5e-06 }
24. 'relative respiration' x,y pairs :{ 0 0.09 2 0.04 6 0.04 1000 0.04 }
25. 'root class ID' = 101 (noUnit)
26. 'root hair density' x,y pairs :{ 0 2000 1 2000 2 2000 10 2000 30 0 2000 0 }
27. 'root hair diameter' = 0.0005 (cm)
28. 'root hair length' x,y pairs :{ 0 0 1 0 2 0.028 2000 0.028 }

29. 'soil impedence.v2'(cm)=f{'uniform distribution'} minimum=-0.020000 maximum=0.020000
30. 'top boundary' = 1 (noUnit)
31. 'topology offset' = 0 (noUnit)

11. 'nodalroots2'(noUnit)
    1. 'aerenchyma formation' x,y pairs :{ 0 0 3 0 5 0.1 10 0.25 20 0.393 1000 0.393 }
    2. 'bottom boundary' = 0 (noUnit)
    3. 'bounce of the side' = 0 (noUnit)
    4. 'branch list'(noUnit)
        1. 'lateral'(noUnit)
            1. 'allow branches to form above ground' = 0 (noUnit)
            2. 'branching frequency'(cm)=f{'uniform distribution'} minimum=0.293000 maximum=0.493000
            3. 'length root tip' = 10.93 (cm)
    5. 'branching angle' = 119.95 (degrees)
    6. 'cannotgrowup' = 1 (noUnit)
    7. 'copy defaults from' = ../defaultsMajorAxis (noUnit)
    8. 'density' = 0.094 (g/cm3)
    9. 'diameter' x,y pairs :{ 0 0.1 8 0.084 13 0.079 100 0.079 }
    10. 'gravitropism.v2'(cm)=f{'uniform distribution'} minimum=-0.010000 maximum=-0.005000
    11. 'growth rate' x,y pairs :{ 0 1.42 1 1.69 2 1.96 3 2.23 4 2.5 5 2.77 7 3.31 10 4.12 12 4.66 13 4.93 40 4.93 }
    12. 'length multiplier to diameter multiplier' x,y pairs :{ 0 0.25 1 1 2 1.5 3 1.8 4 2 100 2 1000 2 }
    13. 'length root tip without xylem vessels' = 2 (cm)
    14. 'local resource responses'(noUnit)
        1. 'impact on:branching frequency'(noUnit)
            1. 'aggregation function' = maxRelativeDeviationFromOne (noUnit)
            2. 'impact by:nitrate' x,y pairs :{ 0 1 2000 1 }
            3. 'impact by:phosphorus' x,y pairs :{ 0 0.2 0.015 1 1000 1 }
            4. 'impact by:potassium' x,y pairs :{ 0 1 1000 1 }
        2. 'impact on:gravitropism'(noUnit)
            1. 'aggregation function' = maxRelativeDeviationFromOne (noUnit)
            2. 'impact by:nitrate' x,y pairs :{ 0 1.5 100 1 2000 1 }
            3. 'impact by:phosphorus' x,y pairs :{ 0 0.5 0.015 2 1000 0.5 }
            4. 'impact by:potassium' x,y pairs :{ 0 1 1000 1 }
        3. 'impact on:root potential longitudinal growth'(noUnit)
            1. 'aggregation function' = maxRelativeDeviationFromOne (noUnit)
            2. 'impact by:nitrate' x,y pairs :{ 0 1 2000 1 }
            3. 'impact by:phosphorus' x,y pairs :{ 0 0.2 0.015 1 1000 1 }
            4. 'impact by:potassium' x,y pairs :{ 0 1 1000 1 }
    15. 'longitudinal growth rate multiplier'(cm)=f{'normal distribution'} minimum=0.600000 maximum=1.200000 mean=1.000000 stdev=0.100000
    16. 'nitrate'(noUnit)
        1. 'Cmin' = 0.001 (umol/ml)
        2. 'Imax' x,y pairs :{ 0 1.21 2 2.1 40 2.1 }
        3. 'Km' x,y pairs :{ 0 0.0157 2 0.0522 40 0.0522 }
        4. 'minimal nutrient concentration' = 600 (umol/g)
        5. 'optimal nutrient concentration' = 1200 (umol/g)
    17. 'number of xylem poles' = 18 (noUnit)
    18. 'phosphorus'(noUnit)
        1. 'Cmin' = 0.0002 (umol/ml)
        2. 'Efflux' = 1e-06 (umol/cm/day)
        3. 'Imax' = 0.0555 (umol/cm2/day)
        4. 'Km' = 0.00545 (umol/ml)
        5. 'minimal nutrient concentration' = 30 (umol/g)
        6. 'optimal nutrient concentration' = 60 (umol/g)
    19. 'potassium'(noUnit)
        1. 'Cmin' = 0.002 (umol/ml)
        2. 'Efflux' = 1e-06 (umol/cm/day)
        3. 'Imax' = 0.467 (umol/cm2/day)
        4. 'Km' = 0.014 (umol/ml)
        5. 'minimal nutrient concentration' = 117 (umol/g)
        6. 'optimal nutrient concentration' = 234 (umol/g)
    20. 'radial hydraulic conductivity' x,y pairs :{ 0 0 1 0.000216 10 0.000216 20 0.00025 30 0.000216 40 0.0001 60 0 }
    21. 'reduction in respiration due to aerenchyma' x,y pairs :{ 0 0 0.3 0.7 0.6 1 }

    22. 'regular topology' = 0 (noUnit)
    23. 'relative carbon cost of exudation' x,y pairs :{ 0 5e-06 100 5e-06 }
    24. 'relative respiration' x,y pairs :{ 0 0.09 2 0.04 6 0.04 1000 0.04 }
    25. 'root class ID' = 101 (noUnit)
    26. 'root hair density' x,y pairs :{ 0 2000 1 2000 2 2000 10 2000 30 0 2000 0 }
    27. 'root hair diameter' = 0.0005 (cm)
    28. 'root hair length' x,y pairs :{ 0 0 1 0 2 0.028 2000 0.028 }
    29. 'soil impedence.v2'(cm)=f{'uniform distribution'} minimum=-0.020000 maximum=0.020000
    30. 'top boundary' = 1 (noUnit)
    31. 'topology offset' = 0 (noUnit)
12. 'nodalroots3'(noUnit)
    1. 'aerenchyma formation' x,y pairs :{ 0 0 3 0 5 0.1 10 0.25 20 0.393 1000 0.393 }
    2. 'bottom boundary' = 0 (noUnit)
    3. 'bounce of the side' = 0 (noUnit)
    4. 'branch list'(noUnit)
        1. 'lateral'(noUnit)
            1. 'allow branches to form above ground' = 0 (noUnit)
            2. 'branching frequency'(cm)=f{'uniform distribution'} minimum=0.100000 maximum=0.300000
            3. 'length root tip' = 10.93 (cm)
    5. 'branching angle' = 110.8 (degrees)
    6. 'cannotgrowup' = 1 (noUnit)
    7. 'copy defaults from' = ../defaultsMajorAxis (noUnit)
    8. 'density' = 0.094 (g/cm3)
    9. 'diameter' x,y pairs :{ 0 0.125 8 0.098 100 0.098 }
    10. 'gravitropism.v2'(cm)=f{'uniform distribution'} minimum=-0.010000 maximum=-0.005000
    11. 'growth rate' x,y pairs :{ 0 0.78 1 1.12 2 1.56 3 1.95 4 2.34 5 2.72 8 3.89 40 3.89 }
    12. 'length multiplier to diameter multiplier' x,y pairs :{ 0 0.25 1 1 2 1.5 3 1.8 4 2 100 2 1000 2 }
    13. 'length root tip without xylem vessels' = 2 (cm)
    14. 'local resource responses'(noUnit)
        1. 'impact on:branching frequency'(noUnit)
            1. 'aggregation function' = maxRelativeDeviationFromOne (noUnit)
            2. 'impact by:nitrate' x,y pairs :{ 0 1 2000 1 }
            3. 'impact by:phosphorus' x,y pairs :{ 0 0.2 0.015 1 1000 1 }
            4. 'impact by:potassium' x,y pairs :{ 0 1 1000 1 }
        2. 'impact on:gravitropism'(noUnit)
            1. 'aggregation function' = maxRelativeDeviationFromOne (noUnit)
            2. 'impact by:nitrate' x,y pairs :{ 0 1.5 100 1 2000 1 }
            3. 'impact by:phosphorus' x,y pairs :{ 0 0.5 0.015 2 1000 0.5 }
            4. 'impact by:potassium' x,y pairs :{ 0 1 1000 1 }
        3. 'impact on:root potential longitudinal growth'(noUnit)
            1. 'aggregation function' = maxRelativeDeviationFromOne (noUnit)
            2. 'impact by:nitrate' x,y pairs :{ 0 1 2000 1 }
            3. 'impact by:phosphorus' x,y pairs :{ 0 0.2 0.015 1 1000 1 }
            4. 'impact by:potassium' x,y pairs :{ 0 1 1000 1 }
    15. 'longitudinal growth rate multiplier'(cm)=f{'normal distribution'} minimum=0.600000 maximum=1.200000 mean=1.000000 stdev=0.100000
    16. 'nitrate'(noUnit)
        1. 'Cmin' = 0.001 (umol/ml)
        2. 'Imax' x,y pairs :{ 0 1.21 2 2.1 40 2.1 }
        3. 'Km' x,y pairs :{ 0 0.0157 2 0.0522 40 0.0522 }
        4. 'minimal nutrient concentration' = 600 (umol/g)
        5. 'optimal nutrient concentration' = 1200 (umol/g)
    17. 'number of xylem poles' = 24 (noUnit)
    18. 'phosphorus'(noUnit)
        1. 'Cmin' = 0.0002 (umol/ml)
        2. 'Efflux' = 1e-06 (umol/cm/day)
        3. 'Imax' = 0.0555 (umol/cm2/day)
        4. 'Km' = 0.00545 (umol/ml)
        5. 'minimal nutrient concentration' = 30 (umol/g)
        6. 'optimal nutrient concentration' = 60 (umol/g)
    19. 'potassium'(noUnit)
        1. 'Cmin' = 0.002 (umol/ml)
        2. 'Efflux' = 1e-06 (umol/cm/day)

        3. 'Imax' = 0.467 (umol/cm2/day)
        4. 'Km' = 0.014 (umol/ml)
        5. 'minimal nutrient concentration' = 117 (umol/g)
        6. 'optimal nutrient concentration' = 234 (umol/g)
    20. 'radial hydraulic conductivity' x,y pairs :{ 0 0 1 0.000216 10 0.000216 20 0.00025 30 0.000216 40 0.0001 60 0 }
    21. 'reduction in respiration due to aerenchyma' x,y pairs :{ 0 0 0.3 0.7 0.6 1 }
    22. 'regular topology' = 0 (noUnit)
    23. 'relative carbon cost of exudation' x,y pairs :{ 0 5e-06 100 5e-06 }
    24. 'relative respiration' x,y pairs :{ 0 0.09 2 0.04 6 0.04 1000 0.04 }
    25. 'root class ID' = 101 (noUnit)
    26. 'root hair density' x,y pairs :{ 0 2000 1 2000 2 2000 10 2000 30 0 2000 0 }
    27. 'root hair diameter' = 0.0005 (cm)
    28. 'root hair length' x,y pairs :{ 0 0 1 0 2 0.028 2000 0.028 }
    29. 'soil impedence.v2'(cm)=f{'uniform distribution'} minimum=-0.020000 maximum=0.020000
    30. 'top boundary' = 1 (noUnit)
    31. 'topology offset' = 0 (noUnit)
13. 'nodalroots4'(noUnit)
    1. 'aerenchyma formation' x,y pairs :{ 0 0 3 0 5 0.1 10 0.25 20 0.393 1000 0.393 }
    2. 'bottom boundary' = 0 (noUnit)
    3. 'bounce of the side' = 0 (noUnit)
    4. 'branch list'(noUnit)
        1. 'lateral'(noUnit)
            1. 'allow branches to form above ground' = 0 (noUnit)
            2. 'branching frequency'(cm)=f{'uniform distribution'} minimum=0.100000 maximum=0.300000
            3. 'length root tip' = 10.93 (cm)
    5. 'branching angle' = 141.88 (degrees)
    6. 'cannotgrowup' = 1 (noUnit)
    7. 'copy defaults from' = ../defaultsMajorAxis (noUnit)
    8. 'density' = 0.094 (g/cm3)
    9. 'diameter' x,y pairs :{ 0 0.2 10 0.11 100 0.11 }
    10. 'gravitropism.v2'(cm)=f{'uniform distribution'} minimum=-0.010000 maximum=-0.005000
    11. 'growth rate' x,y pairs :{ 0 0.01 1 1 3 4.5 28 4.5 38 0 1000 0 }
    12. 'length multiplier to diameter multiplier' x,y pairs :{ 0 0.25 1 1 2 1.5 3 1.8 4 2 100 2 1000 2 }
    13. 'length root tip without xylem vessels' = 2 (cm)
    14. 'local resource responses'(noUnit)
        1. 'impact on:branching frequency'(noUnit)
            1. 'aggregation function' = maxRelativeDeviationFromOne (noUnit)
            2. 'impact by:nitrate' x,y pairs :{ 0 1 2000 1 }
            3. 'impact by:phosphorus' x,y pairs :{ 0 0.2 0.015 1 1000 1 }
            4. 'impact by:potassium' x,y pairs :{ 0 1 1000 1 }
        2. 'impact on:gravitropism'(noUnit)
            1. 'aggregation function' = maxRelativeDeviationFromOne (noUnit)
            2. 'impact by:nitrate' x,y pairs :{ 0 1.5 100 1 2000 1 }
            3. 'impact by:phosphorus' x,y pairs :{ 0 0.5 0.015 2 1000 0.5 }
            4. 'impact by:potassium' x,y pairs :{ 0 1 1000 1 }
        3. 'impact on:root potential longitudinal growth'(noUnit)
            1. 'aggregation function' = maxRelativeDeviationFromOne (noUnit)
            2. 'impact by:nitrate' x,y pairs :{ 0 1 2000 1 }
            3. 'impact by:phosphorus' x,y pairs :{ 0 0.2 0.015 1 1000 1 }
            4. 'impact by:potassium' x,y pairs :{ 0 1 1000 1 }
    15. 'longitudinal growth rate multiplier'(cm)=f{'normal distribution'} minimum=0.600000 maximum=1.200000 mean=1.000000 stdev=0.100000
    16. 'nitrate'(noUnit)
        1. 'Cmin' = 0.001 (umol/ml)
        2. 'Imax' x,y pairs :{ 0 1.21 2 2.1 40 2.1 }
        3. 'Km' x,y pairs :{ 0 0.0157 2 0.0522 40 0.0522 }
        4. 'minimal nutrient concentration' = 600 (umol/g)
        5. 'optimal nutrient concentration' = 1200 (umol/g)
    17. 'number of xylem poles' = 32 (noUnit)
    18. 'phosphorus'(noUnit)
        1. 'Cmin' = 0.0002 (umol/ml)
        2. 'Efflux' = 1e-06 (umol/cm/day)

        3. 'Imax' = 0.0555 (umol/cm2/day)
        4. 'Km' = 0.00545 (umol/ml)
        5. 'minimal nutrient concentration' = 30 (umol/g)
        6. 'optimal nutrient concentration' = 60 (umol/g)
    19. 'potassium'(noUnit)
        1. 'Cmin' = 0.002 (umol/ml)
        2. 'Efflux' = 1e-06 (umol/cm/day)
        3. 'Imax' = 0.467 (umol/cm2/day)
        4. 'Km' = 0.014 (umol/ml)
        5. 'minimal nutrient concentration' = 117 (umol/g)
        6. 'optimal nutrient concentration' = 234 (umol/g)
    20. 'radial hydraulic conductivity' x,y pairs :{ 0 0 1 0.000216 10 0.000216 20 0.00025 30 0.000216 40 0.0001 60 0 }
    21. 'reduction in respiration due to aerenchyma' x,y pairs :{ 0 0 0.3 0.7 0.6 1 }
    22. 'relative carbon cost of exudation' x,y pairs :{ 0 5e-06 100 5e-06 }
    23. 'relative respiration' x,y pairs :{ 0 0.09 2 0.04 6 0.04 1000 0.04 }
    24. 'root class ID' = 101 (noUnit)
    25. 'root hair density' x,y pairs :{ 0 2000 1 2000 2 2000 10 2000 30 0 2000 0 }
    26. 'root hair diameter' = 0.0005 (cm)
    27. 'root hair length' x,y pairs :{ 0 0 1 0 2 0.028 2000 0.028 }
    28. 'soil impedence.v2'(cm)=f{'uniform distribution'} minimum=-0.020000 maximum=0.020000
    29. 'top boundary' = 1 (noUnit)
14. 'primary root'(noUnit)
    1. 'aerenchyma formation' x,y pairs :{ 0 0 3 0 5 0.1 10 0.25 20 0.393 1000 0.393 }
    2. 'bottom boundary' = 0 (noUnit)
    3. 'bounce of the side' = 0 (noUnit)
    4. 'branch list'(noUnit)
        1. 'lateral'(noUnit)
            1. 'allow branches to form above ground' = 0 (noUnit)
            2. 'branching frequency'(cm)=f{'uniform distribution'} minimum=0.064000 maximum=0.264000
            3. 'length root tip' = 10.93 (cm)
        2. 'seminal'(noUnit)
            1. 'allow branches to form above ground' = 0 (noUnit)
            2. 'branching spatial offset' = 0 (cm)
            3. 'branching time offset' = 1 (day)
            4. 'number of branches/whorl' = 0 (#)
    5. 'branching angle' = 0 (degrees)
    6. 'cannotgrowup' = 1 (noUnit)
    7. 'copy defaults from' = ../defaultsMajorAxis (noUnit)
    8. 'density' = 0.094 (g/cm3)
    9. 'diameter' = 0.043 (cm)
    10. 'gravitropism.v2'(cm)=f{'uniform distribution'} minimum=-0.015000 maximum=-0.005000
    11. 'growth rate' x,y pairs :{ 0 1.36 5 1.82 10 2.27 15 2.73 20 3.18 25 3.64 40 3.64 }
    12. 'length multiplier to diameter multiplier' x,y pairs :{ 0 0.25 1 1 2 1.5 3 1.8 4 2 100 2 1000 2 }
    13. 'length root tip without xylem vessels' = 2 (cm)
    14. 'local resource responses'(noUnit)
        1. 'impact on:branching frequency'(noUnit)
            1. 'aggregation function' = maxRelativeDeviationFromOne (noUnit)
            2. 'impact by:nitrate' x,y pairs :{ 0 1 2000 1 }
            3. 'impact by:phosphorus' x,y pairs :{ 0 0.2 0.015 1 1000 1 }
            4. 'impact by:potassium' x,y pairs :{ 0 1 1000 1 }
        2. 'impact on:gravitropism'(noUnit)
            1. 'aggregation function' = maxRelativeDeviationFromOne (noUnit)
            2. 'impact by:nitrate' x,y pairs :{ 0 1.5 100 1 2000 1 }
            3. 'impact by:phosphorus' x,y pairs :{ 0 0.5 0.015 2 1000 0.5 }
            4. 'impact by:potassium' x,y pairs :{ 0 1 1000 1 }
        3. 'impact on:root potential longitudinal growth'(noUnit)
            1. 'aggregation function' = maxRelativeDeviationFromOne (noUnit)
            2. 'impact by:nitrate' x,y pairs :{ 0 1 2000 1 }
            3. 'impact by:phosphorus' x,y pairs :{ 0 0.2 0.015 1 1000 1 }
            4. 'impact by:potassium' x,y pairs :{ 0 1 1000 1 }
    15. 'nitrate'(noUnit)
        1. 'Cmin' = 0.001 (umol/ml)

        2. 'Imax' x,y pairs :{ 0 2.3 2 1.92 40 1.92 }
        3. 'Km' x,y pairs :{ 0 0.0105 2 0.0161 40 0.0161 }
        4. 'minimal nutrient concentration' = 600 (umol/g)
        5. 'optimal nutrient concentration' = 1200 (umol/g)
    16. 'number of xylem poles' = 8 (noUnit)
    17. 'phosphorus'(noUnit)
        1. 'Cmin' = 0.0002 (umol/ml)
        2. 'Efflux' = 1e-06 (umol/cm/day)
        3. 'Imax' = 0.0555 (umol/cm2/day)
        4. 'Km' = 0.00545 (umol/ml)
        5. 'minimal nutrient concentration' = 30 (umol/g)
        6. 'optimal nutrient concentration' = 60 (umol/g)
    18. 'potassium'(noUnit)
        1. 'Cmin' = 0.002 (umol/ml)
        2. 'Efflux' = 1e-06 (umol/cm/day)
        3. 'Imax' = 0.467 (umol/cm2/day)
        4. 'Km' = 0.014 (umol/ml)
        5. 'minimal nutrient concentration' = 117 (umol/g)
        6. 'optimal nutrient concentration' = 234 (umol/g)
    19. 'radial hydraulic conductivity' x,y pairs :{ 0 0 1 0.000216 10 0.000216 20 0.000216 30 0.000116 40 5e-05 60 0 }
    20. 'reduction in respiration due to aerenchyma' x,y pairs :{ 0 0 0.3 0.7 0.6 1 }
    21. 'relative carbon cost of exudation' x,y pairs :{ 0 5e-06 100 5e-06 }
    22. 'relative respiration' x,y pairs :{ 0 0.09 2 0.04 6 0.04 1000 0.04 }
    23. 'root class ID' = 100 (noUnit)
    24. 'root hair density' x,y pairs :{ 0 2000 1 2000 2 2000 10 2000 30 0 2000 0 }
    25. 'root hair diameter' = 0.0005 (cm)
    26. 'root hair length' x,y pairs :{ 0 0 1 0 2 0.028 2000 0.028 }
    27. 'soil impedence.v2'(cm)=f{'uniform distribution'} minimum=-0.050000 maximum=0.050000
    28. 'top boundary' = 1 (noUnit)
15. 'resources'(noUnit)
    1. 'Cto dry weight ratio' = 0.45 (100%)
    2. 'carbon allocation to leafs factor' x,y pairs :{ 0 1 10 0.7 20 0.45 33 0.42 40 0.4 60 0.4 }
    3. 'carbon allocation to roots factor' x,y pairs :{ 0 1 1 1 6 0.4 20 0.2 40 0.17 80 0.17 }
    4. 'carbon cost of nitrate uptake' = 1.392e-05 (g/umol)
    5. 'max carbon allocation to shoot' = 0.82 (100%)
    6. 'nitrate'(noUnit)
        1. 'initial nutrient uptake' = 20.07 (umol)
    7. 'phosphorus'(noUnit)
        1. 'initial nutrient uptake' = 2.96 (umol)
    8. 'potassium'(noUnit)
        1. 'initial nutrient uptake' = 27 (umol)
    9. 'reserve allocation rate' x,y pairs :{ 0 0.01 1 0.02 2 0.04 3 0.04 10 0.2 11 0.2 1000 0.2 }
    10. 'seed carbohydrate content' = 0.5292 (100%)
    11. 'seed carbohydrate to CFactor' = 0.444 (100%)
    12. 'seed reserve duration' = 100 (day)
    13. 'seed size' = 0.015 (g)
16. 'seminal'(noUnit)
    1. 'aerenchyma formation' x,y pairs :{ 0 0 3 0 5 0.1 10 0.25 20 0.393 1000 0.393 }
    2. 'bottom boundary' = 0 (noUnit)
    3. 'bounce of the side' = 0 (noUnit)
    4. 'branch list'(noUnit)
        1. 'lateral'(noUnit)
            1. 'allow branches to form above ground' = 0 (noUnit)
            2. 'branching frequency'(cm)=f{'uniform distribution'} minimum=0.182000 maximum=0.482000
            3. 'length root tip' = 10.93 (cm)
    5. 'branching angle' = 119.64 (degrees)
    6. 'cannotgrowup' = 1 (noUnit)
    7. 'copy defaults from' = ../defaultsMajorAxis (noUnit)
    8. 'density' = 0.094 (g/cm3)
    9. 'diameter' = 0.074 (cm)
    10. 'gravitropism.v2'(cm)=f{'uniform distribution'} minimum=-0.035000 maximum=-0.025000

11. 'growth rate' x,y pairs :{ 0 1.29 1 1.51 2 1.72 3 1.94 4 2.15 5 2.37 7 2.8 10 3.44 12 3.87 15 4.52 17 4.95 20 5.59 22 6.02 25 6.67 40 6.67 }
12. 'length multiplier to diameter multiplier' x,y pairs :{ 0 0.25 1 1 2 1.5 3 1.8 4 2 100 2 1000 2 }
13. 'length root tip without xylem vessels' = 2 (cm)
14. 'local resource responses'(noUnit)
    1. 'impact on:branching frequency'(noUnit)
        1. 'aggregation function' = maxRelativeDeviationFromOne (noUnit)
        2. 'impact by:nitrate' x,y pairs :{ 0 1 2000 1 }
        3. 'impact by:phosphorus' x,y pairs :{ 0 0.2 0.015 1 1000 1 }
        4. 'impact by:potassium' x,y pairs :{ 0 1 1000 1 }
    2. 'impact on:gravitropism'(noUnit)
        1. 'aggregation function' = maxRelativeDeviationFromOne (noUnit)
        2. 'impact by:nitrate' x,y pairs :{ 0 1.5 100 1 2000 1 }
        3. 'impact by:phosphorus' x,y pairs :{ 0 0.5 0.015 2 1000 0.5 }
        4. 'impact by:potassium' x,y pairs :{ 0 1 1000 1 }
    3. 'impact on:root potential longitudinal growth'(noUnit)
        1. 'aggregation function' = maxRelativeDeviationFromOne (noUnit)
        2. 'impact by:nitrate' x,y pairs :{ 0 1 2000 1 }
        3. 'impact by:phosphorus' x,y pairs :{ 0 0.2 0.015 1 1000 1 }
        4. 'impact by:potassium' x,y pairs :{ 0 1 1000 1 }
15. 'longitudinal growth rate multiplier'(cm)=f{'normal distribution'} minimum=0.600000 maximum=1.200000 mean=1.000000 stdev=0.100000
16. 'nitrate'(noUnit)
    1. 'Cmin' = 0.001 (umol/ml)
    2. 'Imax' x,y pairs :{ 0 2.3 2 1.92 40 1.92 }
    3. 'Km' x,y pairs :{ 0 0.0105 2 0.0161 40 0.0161 }
    4. 'minimal nutrient concentration' = 600 (umol/g)
    5. 'optimal nutrient concentration' = 1200 (umol/g)
17. 'number of xylem poles' = 6 (noUnit)
18. 'phosphorus'(noUnit)
    1. 'Cmin' = 0.0002 (umol/ml)
    2. 'Efflux' = 1e-06 (umol/cm/day)
    3. 'Imax' = 0.0555 (umol/cm2/day)
    4. 'Km' = 0.00545 (umol/ml)
    5. 'minimal nutrient concentration' = 30 (umol/g)
    6. 'optimal nutrient concentration' = 60 (umol/g)
19. 'potassium'(noUnit)
    1. 'Cmin' = 0.002 (umol/ml)
    2. 'Efflux' = 1e-06 (umol/cm/day)
    3. 'Imax' = 0.467 (umol/cm2/day)
    4. 'Km' = 0.014 (umol/ml)
    5. 'minimal nutrient concentration' = 117 (umol/g)
    6. 'optimal nutrient concentration' = 234 (umol/g)
20. 'radial hydraulic conductivity' x,y pairs :{ 0 0 1 0.000216 10 0.000216 20 0.00025 30 0.000216 40 0.0001 60 0 }
21. 'reduction in respiration due to aerenchyma' x,y pairs :{ 0 0 0.3 0.7 0.6 1 }
22. 'regular topology' = 1 (noUnit)
23. 'relative carbon cost of exudation' x,y pairs :{ 0 5e-06 100 5e-06 }
24. 'relative respiration' x,y pairs :{ 0 0.09 2 0.04 6 0.04 1000 0.04 }
25. 'root class ID' = 99 (noUnit)
26. 'root hair density' x,y pairs :{ 0 2000 1 2000 2 2000 10 2000 30 0 2000 0 }
27. 'root hair diameter' = 0.0005 (cm)
28. 'root hair length' x,y pairs :{ 0 0 1 0 2 0.028 2000 0.028 }
29. 'soil impedence.v2'(cm)=f{'uniform distribution'} minimum=-0.040000 maximum=0.040000
30. 'top boundary' = 1 (noUnit)

17. 'shoot'(noUnit)
    1. 'aerenchyma photosynthesis mitigation' = 0.5 (100%)
    2. 'area per plant' = 1600 (cm2)
    3. 'extinction coefficient' = 0.85 (noUnit)
    4. 'leaf area expantion rate' x,y pairs :{ 0 0 2 0 5 0 6 0.134 7 0.87 8 1.61 9 2.34 10 3.08 11 3.81 12 4.55 13 4.87 14 7.009 15 9.148 16 11.287 17 13.426 18 15.565 19 17.704 20 19.843 21 21.982 22 24.121 23 26.26 24 28.399 25 30.538 40 30.538 }
    5. 'light use efficiency' = 3.8e-07 (g/umol)

6. 'nitrate'(noUnit)
    1. 'leaf minimal nutrient concentration' x,y pairs :{ 0 1200 80 800 }
    2. 'leaf optimal nutrient concentration' x,y pairs :{ 0 2500 80 1500 }
    3. 'stem minimal nutrient concentration' = 400 (umol/g)
    4. 'stem optimal nutrient concentration' = 800 (umol/g)
7. 'phosphorus'(noUnit)
    1. 'leaf minimal nutrient concentration' = 35 (umol/g)
    2. 'leaf optimal nutrient concentration' = 70 (umol/g)
    3. 'stem minimal nutrient concentration' = 15 (umol/g)
    4. 'stem optimal nutrient concentration' = 30 (umol/g)
8. 'potassium'(noUnit)
    1. 'leaf minimal nutrient concentration' = 273 (umol/g)
    2. 'leaf optimal nutrient concentration' = 508 (umol/g)
    3. 'stem minimal nutrient concentration' = 117 (umol/g)
    4. 'stem optimal nutrient concentration' = 250 (umol/g)
9. 'relative potential transpiration' = 100 (cm3/g)
10. 'relative respiration rate leafs' = 0.04 (g/g/day)
11. 'relative respiration rate stems' = 0.02 (g/g/day)
12. 'specific leaf area' x,y pairs :{ 0 0.0015 24 0.0026 50 0.0032 100 0.0032 }

18. 'stress impact factors'(noUnit)
    1. 'impact on:branching frequency multiplier'(noUnit)
        1. 'impact by:nitrate' x,y pairs :{ 0 1 0.5 1 1 0 }
        2. 'impact by:phosphorus' x,y pairs :{ 0 1 0.5 1 1 0 }
        3. 'impact by:potassium' x,y pairs :{ 0 1 0.5 1 1 0 }
    2. 'impact on:gravitropism multiplier'(noUnit)
        1. 'impact by:nitrate' x,y pairs :{ 0 1 0.5 1 1 0 }
        2. 'impact by:phosphorus' x,y pairs :{ 0 1 0.5 1 1 0 }
        3. 'impact by:potassium' x,y pairs :{ 0 1 0.5 1 1 0 }
    3. 'impact on:leaf area expantion rate'(noUnit)
        1. 'impact by:nitrate' x,y pairs :{ 0 0 0.3 0.1 1 1 }
        2. 'impact by:phosphorus' x,y pairs :{ 0 0 1 1 }
        3. 'impact by:potassium' x,y pairs :{ 0 0 0.2 0.5 1 1 }
    4. 'impact on:leaf respiration'(noUnit)
        1. 'impact by:nitrate' x,y pairs :{ 0 1 1 1 }
        2. 'impact by:phosphorus' x,y pairs :{ 0 1 1 1 }
        3. 'impact by:potassium' x,y pairs :{ 0 1 1 1 }
    5. 'impact on:photosynthesis'(noUnit)
        1. 'impact by:nitrate' x,y pairs :{ 0 0 0.4 0.5 1 1 }
        2. 'impact by:phosphorus' x,y pairs :{ 0 0.5 0.5 1 1 1 }
        3. 'impact by:potassium' x,y pairs :{ 0 0 1 1 }
    6. 'impact on:root potential longitudinal growth'(noUnit)
        1. 'impact by:nitrate' x,y pairs :{ 0 0 0.5 1 1 1 }
        2. 'impact by:phosphorus' x,y pairs :{ 0 1 0.1 1 1 1 }
        3. 'impact by:potassium' x,y pairs :{ 0 0 0.5 1 1 1 }
    7. 'impact on:root potential longitudinal growth multiplier'(noUnit)
        1. 'impact by:nitrate' x,y pairs :{ 0 1 0.5 1 1 0 }
        2. 'impact by:phosphorus' x,y pairs :{ 0 1 0.5 1 1 0 }
        3. 'impact by:potassium' x,y pairs :{ 0 1 0.5 1 1 0 }
    8. 'impact on:root segment carbon cost of exudates'(noUnit)
        1. 'impact by:nitrate' x,y pairs :{ 0 1 1 1 }
        2. 'impact by:phosphorus' x,y pairs :{ 0 1 1 1 }
        3. 'impact by:potassium' x,y pairs :{ 0 1 1 1 }
    9. 'impact on:root segment respiration'(noUnit)
        1. 'impact by:nitrate' x,y pairs :{ 0 1 1 1 }
        2. 'impact by:phosphorus' x,y pairs :{ 0 1 1 1 }
        3. 'impact by:potassium' x,y pairs :{ 0 1 1 1 }
    10. 'impact on:root segment secondary growth'(noUnit)
        1. 'impact by:nitrate' x,y pairs :{ 0 0 1 1 }
        2. 'impact by:phosphorus' x,y pairs :{ 0 0 1 1 }
        3. 'impact by:potassium' x,y pairs :{ 0 0 1 1 }
    11. 'impact on:stem respiration'(noUnit)
        1. 'impact by:nitrate' x,y pairs :{ 0 1 1 1 }
        2. 'impact by:phosphorus' x,y pairs :{ 0 1 1 1 }

3. 'impact by:potassium' x,y pairs :{ 0 1 1 1 }

7. 'shoot template'(noUnit)
    1. 'area per plant'
    2. 'carbon allocation to leafs'(g) initial value = 0 0 0
    3. 'carbon allocation to stems'(g) initial value = 0 0 0
    4. 'extinction coefficient'
    5. 'leaf area'(cm2) initial value = 0 0 0
    6. 'leaf area index'(cm2/cm2)
    7. 'leaf area reduction coefficient'(cm2/cm2)
    8. 'leaf dry weight'(g) initial value = 0 0 0
    9. 'leaf potential carbon sink for growth'(g) initial value = 0 0 0
    10. 'leaf respiration'(g) initial value = 0 0 0
    11. 'light interception'(umol/cm2/day)
    12. 'photosynthesis'(g) initial value = 0 0 0
    13. 'potential leaf area'(cm2) initial value = 0 0 0
    14. 'relative carbon allocation to leafs'(100%)
    15. 'relative carbon allocation to stems'(100%)
    16. 'stem dry weight'(g) initial value = 0 0 0
    17. 'stem potential carbon sink for growth'(g) initial value = 0 0 0
    18. 'stem respiration'(g) initial value = 0 0 0
    19. 'stress adjusted potential leaf area'(cm2) initial value = 0 0 0
8. 'sibling root template'(noUnit)
    1. 'branches'(noUnit)
    2. 'data points'(noUnit)
    3. 'growthpoint'(cm) initial position = 0 0 0 0 0 0 0
        1. 'branching frequency multiplier'(noUnit)
        2. 'gravitropism'
            1. 'multiplier'(noUnit)
        3. 'root circumference'(cm)
        4. 'root diameter'(cm)
        5. 'root longitudinal growth'(cm) initial value = 0 0 0
        6. 'root potential longitudinal growth'(cm) initial value = 0 0 0
            1. 'rate multiplier'(noUnit)
        7. 'root potential secondary growth'(cm)
        8. 'root segment age' = 0 (day)
        9. 'root segment carbon cost of exudates' = 0 (g)
        10. 'root segment dry weight' = 0 (g)
        11. 'root segment length' = 0 (cm)
        12. 'root segment length duration' = 0 (cm.day)
        13. 'root segment potential carbon sink for growth'(g) initial value = 0 0 0
        14. 'root segment respiration' = 0 (g)
        15. 'root segment secondary potential carbon sink for growth' = 0 (g)
        16. 'root segment specific weight' = 0 (g/cm3)
        17. 'root segment surface area' = 0 (cm2)
        18. 'root segment volume' = 0 (cm3)
    4. 'root carbon cost of exudates'(g) initial value = 0 0 0
    5. 'root dry weight'(g)
    6. 'root length'(cm)
    7. 'root respiration'(g) initial value = 0 0 0
    8. 'root secondary potential carbon sink for growth'(g) initial value = 0 0 0
    9. 'root surface area'(cm2)
    10. 'root system carbon cost of exudates'(g) initial value = 0 0 0
    11. 'root system dry weight'(g)
    12. 'root system length'(cm)
    13. 'root system longitudinal growth'(cm) initial value = 0 0 0
    14. 'root system potential carbon sink for growth'(g) initial value = 0 0 0
    15. 'root system potential carbon sink for growth;major axis'(g) initial value = 0 0 0
        1. 'included root classes' = hypocotyl, primaryRoot, seminal, nodalroots, nodalroots1, nodalroots2, nodalroots3, nodalroots4, nodalroots5, braceroots, braceroots1, braceroots2, braceroots3, basalWhorl1, basalWhorl2, basalWhorl3, basalWhorl4 (noUnit)
    16. 'root system respiration'(g) initial value = 0 0 0
    17. 'root system secondary potential carbon sink for growth'(g) initial value = 0 0 0
    18. 'root system surface area'(cm2)

    19. 'root system volume'(cm3)
    20. 'root volume'(cm3)
9. 'simulation controls'(noUnit)
    1. 'Simula stochastic::number of samples' = 1 (noUnit)
    2. 'aerenchyma'(noUnit)
        1. 'include photosynthesis effects' = 0 (noUnit)
        2. 'reduce respiration' = 0 (noUnit)
        3. 'remobilize p' = 0 (noUnit)
    3. 'integration parameters'(noUnit)
        1. 'default spatial integration length' = 1 (cm)
    4. 'output parameters'(noUnit)
        1. 'RSML'(noUnit)
            1. 'requested variables' = rootLength, rootSurfaceArea (noUnit)
            2. 'run' = 1 (noUnit)
            3. 'time interval' = 2 (noUnit)
        2. 'VTU'(noUnit)
            1. 'include point data' = 1 (noUnit)
            2. 'include roots' = 1 (noUnit)
            3. 'include shoots' = 0 (noUnit)
            4. 'include VTUFor depletion zones' = 1 (noUnit)
            5. 'run' = 1 (noUnit)
            6. 'time interval' = 1 (noUnit)
        3. 'defaults'(noUnit)
            1. 'end time' = 25 (noUnit)
            2. 'start time' = 0 (noUnit)
            3. 'time interval' = 1 (noUnit)
        4. 'garbage collection'(noUnit)
            1. 'lack time' = 1 (noUnit)
            2. 'run' = 1 (noUnit)
            3. 'start time' = 5 (noUnit)
            4. 'time interval' = 1 (noUnit)
        5. 'model dump'(noUnit)
            1. 'end time' = 1 (noUnit)
            2. 'run' = 1 (noUnit)
            3. 'start time' = 0 (noUnit)
        6. 'probe all objects'(noUnit)
            1. 'run' = 0 (noUnit)
            2. 'time interval' = 1 (noUnit)
        7. 'raster image'(noUnit)
            1. 'edge smoothing' = 0 (cm)
            2. 'include binary raw image' = 0 (noUnit)
            3. 'include VTKImage' = 0 (noUnit)
            4. 'overlap function' = additive (noUnit)
            5. 'radius comes from' = rootDiameter,phosphorus/radiusDepletionZone (noUnit)
            6. 'root diameter scaling factor' = 1 (100%)
            7. 'run' = 0 (noUnit)
            8. 'start time' = 5 (noUnit)
            9. 'substeps' = 1 (#)
            10. 'time interval' = 5 (noUnit)
            11. 'voxelsize' = 0.1 (cm)
        8. 'root statistics'(noUnit)
            1. 'requested variables' = rootLength, rootSurfaceArea (noUnit)
            2. 'run' = 0 (noUnit)
            3. 'time interval' = 5 (noUnit)
        9. 'table'(noUnit)
            1. 'run' = 1 (noUnit)
            2. 'searching depth' = 5 (noUnit)
            3. 'skip these variables' = primaryRoot, hypocotyl, (noUnit)
        10. 'table of root nodes'(noUnit)
            1. 'run' = 0 (noUnit)
            2. 'time interval' = 8 (noUnit)
        11. 'vtp'(noUnit)
            1. 'run' = 1 (noUnit)

            2. 'time interval' = 1 (noUnit)

10. 'soil'(noUnit)
    1. 'Root length profile_00-10'(cm)
        1. 'y1' = 0 (noUnit)
        2. 'y2' = -10 (noUnit)
    2. 'Root length profile_10-20'(cm)
        1. 'y1' = -10 (noUnit)
        2. 'y2' = -20 (noUnit)
    3. 'Root length profile_ to 0-30'(cm)
        1. 'y1' = -20 (noUnit)
        2. 'y2' = -30 (noUnit)
    4. 'Root length profile_30-40'(cm)
        1. 'y1' = -30 (noUnit)
        2. 'y2' = -40 (noUnit)
    5. 'Root length profile_ for 0-50'(cm)
        1. 'y1' = -40 (noUnit)
        2. 'y2' = -50 (noUnit)
    6. 'Root length profile_50-60'(cm)
        1. 'y1' = -50 (noUnit)
        2. 'y2' = -60 (noUnit)
    7. 'Root length profile_60-70'(cm)
        1. 'y1' = -60 (noUnit)
        2. 'y2' = -70 (noUnit)
    8. 'Root length profile_70-80'(cm)
        1. 'y1' = -70 (noUnit)
        2. 'y2' = -80 (noUnit)
    9. 'Root length profile_80-90'(cm)
        1. 'y1' = -80 (noUnit)
        2. 'y2' = -90 (noUnit)
    10. 'Root length profile_90+'(cm)
        1. 'y1' = -90 (noUnit)
        2. 'y2' = -1000 (noUnit)
    11. 'between the row coring'(noUnit)
        1. '05.500,00.000'(cm)
            1. 'center' = 5.5 0 0 (cm)
            2. 'coring depth' = -200 (cm)
            3. 'radius' = 4.5 (cm)
            4. 'vertical spacing' = 10 (cm)
    12. 'in the row coring'(noUnit)
        1. '00.000,00.000'(cm)
            1. 'center' = 0 0 0 (cm)
            2. 'coring depth' = -200 (cm)
            3. 'radius' = 4.5 (cm)
            4. 'vertical spacing' = 10 (cm)
